# Cortical-like dynamics in recurrent circuits optimized for sampling-based probabilistic inference

**DOI:** 10.1101/696088

**Authors:** Rodrigo Echeveste, Laurence Aitchison, Guillaume Hennequin, Máté Lengyel

**Affiliations:** Computational and Biological Learning Lab, Dept. of Engineering, University of Cambridge, Cambridge, UK; Research Institute for Signals, Systems and Computational Intelligence sinc(i) (FICH-UNL/CONICET), Santa Fe, Argentina; Center for Cognitive Computation, Department of Cognitive Science, Central European University, Budapest, Hungary

## Abstract

Sensory cortices display a suite of ubiquitous dynamical features, such as ongoing noise variability, transient overshoots, and oscillations, that have so far escaped a common, principled theoretical account. We developed a unifying model for these phenomena by training a recurrent excitatory–inhibitory neural circuit model of a visual cortical hypercolumn to perform sampling-based probabilistic inference. The optimized network displayed several key biological properties, including divisive normalization, as well as stimulus-modulated noise variability, inhibition-dominated transients at stimulus onset, and strong gamma oscillations. These dynamical features had distinct functional roles in speeding up inferences and made predictions that we confirmed in novel analyses of awake monkey recordings. Our results suggest that the basic motifs of cortical dynamics emerge as a consequence of the efficient implementation of the same computational function—fast sampling-based inference—and predict further properties of these motifs that can be tested in future experiments.

---

The dynamics of sensory cortices exhibit a set of features that appear ubiquitously across species and experimental conditions. Responses vary over time and across trials even when the same static stimulus is presented^1^, and these intrinsic variations have both systematic and seemingly random components (so-called noise variability). The most prominent systematic patterns of neural activity are strong, inhibition-dominated transients at stimulus onset2 (or, equivalently, strong adaptation following stimulus onset), and stimulus-dependent population oscillations in the gamma band (20–80 Hz)^3,4^. The extent and pattern of noise variability is also stimulus-dependent: variability is quenched at stimulus onset^1^, decreasing gradually with stimulus contrast in the primary visual cortex (V1)^5,6^, and is further modulated by the content of the stimulus, e.g. the orientation or direction of drifting gratings for cells in V1 or in the middle temporal visual area (MT)^7,8^.

While the mechanisms giving rise to these dynamical phenomena are increasingly well understood^8–10^, their functional significance remains largely unknown and controversial, with several candidate functional roles having been proposed for each of them. For example, cortical gamma oscillations have been suggested to be a substrate for binding different sources of information about a feature (known as binding by synchrony)^11,12^, to mediate information routing (communication by synchrony)^13^, or to enable a temporal code of spikes relative to the oscillation phase^14^. Additionally, transient overshoots have been proposed to carry novelty or prediction error signals^15^. Noise variability, when considered to have any function at all rather than being mere nuisance^16^, has been argued to bear signatures of specific probabilistic computations in the cortex^6,17,18^. However, it is unclear whether these explanations can be reconciled, as each of them only accounts for select aspects of the data, and has been challenged by alternative accounts^3, 19–21^.

Here, we present a unifying model in which all of these dynamical phenomena emerge as a consequence of the efficient implementation of the same computational function: probabilistic inference. Probabilistic inference provides a principled solution for the fundamental requirement of perception to continually fuse partial and noisy information from multiple sources (including multiple sensory cues, modalities, and forms of memory)^22,23^. Formally, the result of this fusion is a posterior distribution. The posterior distribution expresses the probability that relevant features in the environment that are not directly accessible to the brain (e.g. the three-dimensional shapes of objects) may take any particular configuration given information that is directly available to our senses (e.g. photons absorbed in the retina). Behavioral evidence in several domains, including near-optimal performance in multi-sensory integration, decision making, motor control, and learning suggests that the brain represents posterior distributions at least approximately^24^. There have also been several proposals for how the neural responses of sensory cortical populations may implement these probabilistic representations^6–17,25^. While these models successfully explained important aspects of *stationary* response distributions (e.g. tuning curves, Fano factors, noise correlations), they have so far fallen short of accounting for the rich intrinsic *dynamics* of sensory cortical areas.

To bring together dynamics (cortical-like activity patterns) and function (representing posterior distributions) in a principled manner, we optimized a biologically constrained recurrent neural network for performing probabilistic inference. In particular, the biological constraints included a separation of excitatory and inhibitory cells, and nonsaturating firing rates in the physiological regime. On each trial, the network received a different visual stimulus as input, and it was required to perform inference by representing in its responses the posterior distribution that would be inferred for the same stimulus by a Bayesian ideal observer. Specifically, network dynamics had to produce activities which represented statistical samples from the posterior distribution. Importantly, this required the network to modulate not only the mean but also the variability of its responses in a stimulus-dependent manner. Such a sampling-based probabilistic representation of uncertainty has been shown to account for stimulus- and task-dependent aspects of *stationary* variability in V1^6,26,27^, thus offering a promising computational target for a network used to study the temporal *dynamics* of V1 responses.

The optimized neural circuit exhibited a number of appealing computational and dynamical features. Computationally, after training on a reduced stimulus set, the network exhibited strong forms of generalization by producing near-optimal response distributions to novel inputs which required qualitatively different responses. Furthermore, the network discovered out-of-equilibrium dynamics, a strategy currently employed by modern machine learning algorithms to produce samples that become statistically independent on short timescales^28^. Biologically, the circuit achieved divisive normalization of its outputs and displayed marked transients at stimulus onset, as well as strong gamma oscillations. Both the magnitude of transients and the frequency of gamma oscillations scaled with stimulus contrast. Crucially, these dynamical phenomena did not emerge in a control network trained to match posterior mean responses only, without a need to modulate the variability of its responses. Indeed, further analyses of transients and oscillations in the optimized network revealed distinct functional roles for them. Our results also allowed us to predict novel properties of cortical dynamics. For example, we predicted that onset transients should be tuned to stimuli, which we confirmed by performing novel analyses of published V1 recordings in the awake monkey^29^. In addition, our model also made specific predictions about the stimulus tuning of excitatory–inhibitory lags and the distribution of gamma power across the different modes of network dynamics. Both can be readily tested in future experiments.

In summary, we constructed the first biologically constrained recurrent neural network performing sampling-based probabilistic inference that explained a plethora of electrophysiological observations in sensory cortices. Our model thus provides a unifying theoretical account of the basic motifs of sensory cortical dynamics.

## Results

### Optimizing a recurrent neural circuit for probabilistic inference

To study neural circuit dynamics implementing probabilistic inference, we used a novel combination of two well-established, though hitherto unrelated computational approaches: Bayesian ideal observers and the training of recurrent neural networks. First, we used a Bayesian ideal observer model to specify the computational goal of perceptual inference in a simplified visual task. Performing inference requires an internal model that encapsulates one’s assumptions about how the inputs to be processed have been generated by the environment. For this, we adopted the Gaussian scale mixture (GSM) model (Fig. 1a, Online Methods), a generative model that has been shown to capture the statistics of natural image patches^30^. Conversely, inference under the GSM model has been shown to account for behavioral and neural data (for stationary responses) in visual perception^6,31,32^. The GSM model assumes that an image patch is generated as a linear combination of oriented Gabor filter-like visual features (“projective fields”), each present with a different intensity (the latent variables of the model). The image patch is further scaled by a single global “contrast” variable. Following previous approaches to studying V1 dynamics^8,33,34^, we focused on modelling a single hypercolumn. Therefore, we chose the projective fields of the GSM latent variables to only differ in their orientation so that they formed a ring topology (Fig. 1a, Fig. S1a). The ideal observer was obtained by a Bayesian inversion of this model (Online Methods). Thus, for every image patch taken as sensory input, the ideal observer yielded a high-dimensional posterior distribution quantifying the probability that any particular joint combination of intensities for the Gabor-like projective fields may have generated the input (Fig. 1b).

**Fig. 1.**
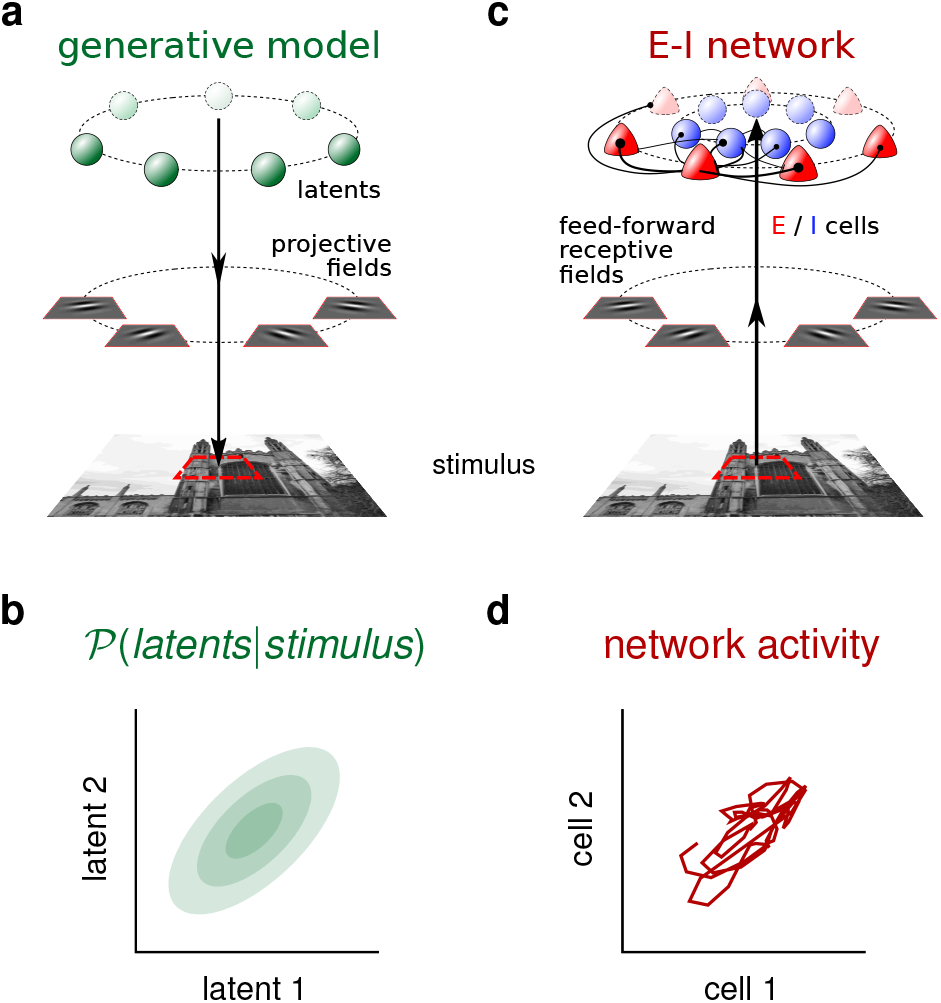
The statistical generative model, and the corresponding neural circuit implementing sampling-based probabilistic inference. **a,** Sketch of the Gaussian scale mixture (GSM) generative model. An image patch is constructed as a linear combination of a fixed set of localized, oriented, Gabor filter-like features (projective fields, differing only in their orientations, uniformly spread between −90° and 90°), with stimulus-specific feature intensities (latent variables) drawn from a multivariate Gaussian distribution. The resulting image patch is scaled by a global contrast variable and corrupted by noise (not shown). (Stimulus shown is for illustration only: the GSM model employed here was not sufficiently complex to generate photorealistic images. For a sample of generated image patches see Fig. 3c and Fig. S1c.) **b,** 2-dimensional projection of the posterior distribution over latent variables given a visual stimulus, computed by the Bayesian ideal observer under the generative model. **c,** An excitatory–inhibitory (E–I) neural network receiving an image patch as an input, filtered by feed-forward receptive fields identical to the projective fields of the generative model in **a**. The activity of each E cell represents the value of one latent variable in the generative model. As an illustration of the ring topology of the network, the outgoing connections of one E-I cell pair are shown (connection strength is indicated by line width and “synapse” size, see details in text). **d,** Responses of the two E cells corresponding to the latent variables shown in **b**. The response trajectory samples from the corresponding posterior distribution over time given the same stimulus.

Second, to model cortical circuit dynamics, we used a canonical, rate-based stochastic recurrent neural network model, the stochastic variant of the stabilized supralinear network (SSN)^8,35,36^ (Online Methods). The network was constrained to exhibit some basic biological features that have been shown to have fundamental consequences for cortical dynamics: the presence of separate but inter-connected excitatory (E) and inhibitory (I) populations of neurons (Fig. 1c), supralinear (expansive) input/output functions^35,37^, and some finite and stimulus-independent process noise^8,38^ incorporating intrinsic and extrinsic forms of neural variability.

We trained this network to perform sampling-based inference under the GSM. For this, the membrane potential response of each excitatory cell at any point in time was taken to represent a possible level of intensity of the corresponding projective field of the GSM model. The activities of inhibitory neurons were treated as auxiliary variables which were not explicitly constrained by the computational objective. The network was optimized to produce distributions of excitatory neural activities that matched the posteriors computed by the GSM-based ideal observer up to second-order statistics (Online Methods). In other words, for each stimulus in a small training set, the network was required to use its stochastic dynamics to visit different parts of state space over time with a frequency proportional to the posterior distribution corresponding to the same stimulus (Fig. 1d). Critically, as process noise in the network was stimulus-independent, the network had to use its recurrent dynamics to shape this variability appropriately for matching the target posteriors for each input. The training objective also included terms encouraging fast circuit dynamics (Online Methods). Thus, the network had to generate fast fluctuations with the correct Optimizing our network was challenging because modulating response variability (to match the stimulus-dependent posterior covariances of the ideal observer model) requires strong and nonlinear recurrent interactions, but networks of strongly connected excitatory neurons – especially with supralinear input/output functions – are prone to becoming unstable^8,39^. In such networks, it is non-trivial to find parameter regimes in which stability is preserved and thus optimization can proceed^8^. We therefore chose a reduced parametrization of our network, analogous to classical “ring” models of a V1 hyper-column^8–33,34^. Recurrent connection strength between any two cells (and the covariability of their process noise) only depended on the angular distance between their preferred stimuliand their respective cell types (E or I; Fig. S3a, Online Methods). The feed-forward receptive fields of the cells were fixed and identical to the projective fields of the corresponding latent variables of the ring-structured GSM (Fig. S1a).

### Inference and generalization in the optimized network

In line with neural recordings, activity in the optimized network was highly variable across time and trials, in response to both low-contrast (Fig. 2a, top) and high-contrast stimuli (Fig. 2a, bottom). Critically, the distributions of neural responses at the five training stimuli (the same image patch at five different contrast levels; Fig. 2b left) closely matched the corresponding GSM posteriors (Fig. 2b–d, compare red to green). Specifically, the mean activity of neurons increased while the variability of their responses decreased with contrast as well as with the match between stimulus orientation and their preferred orientation. This was consistent with the behaviour of the moments of the GSM posterior (Fig. 2c, and circles in Fig. 3a). Thus, the network had been trained successfully to perform sampling-based inference on these stimuli.

**Fig. 2.**
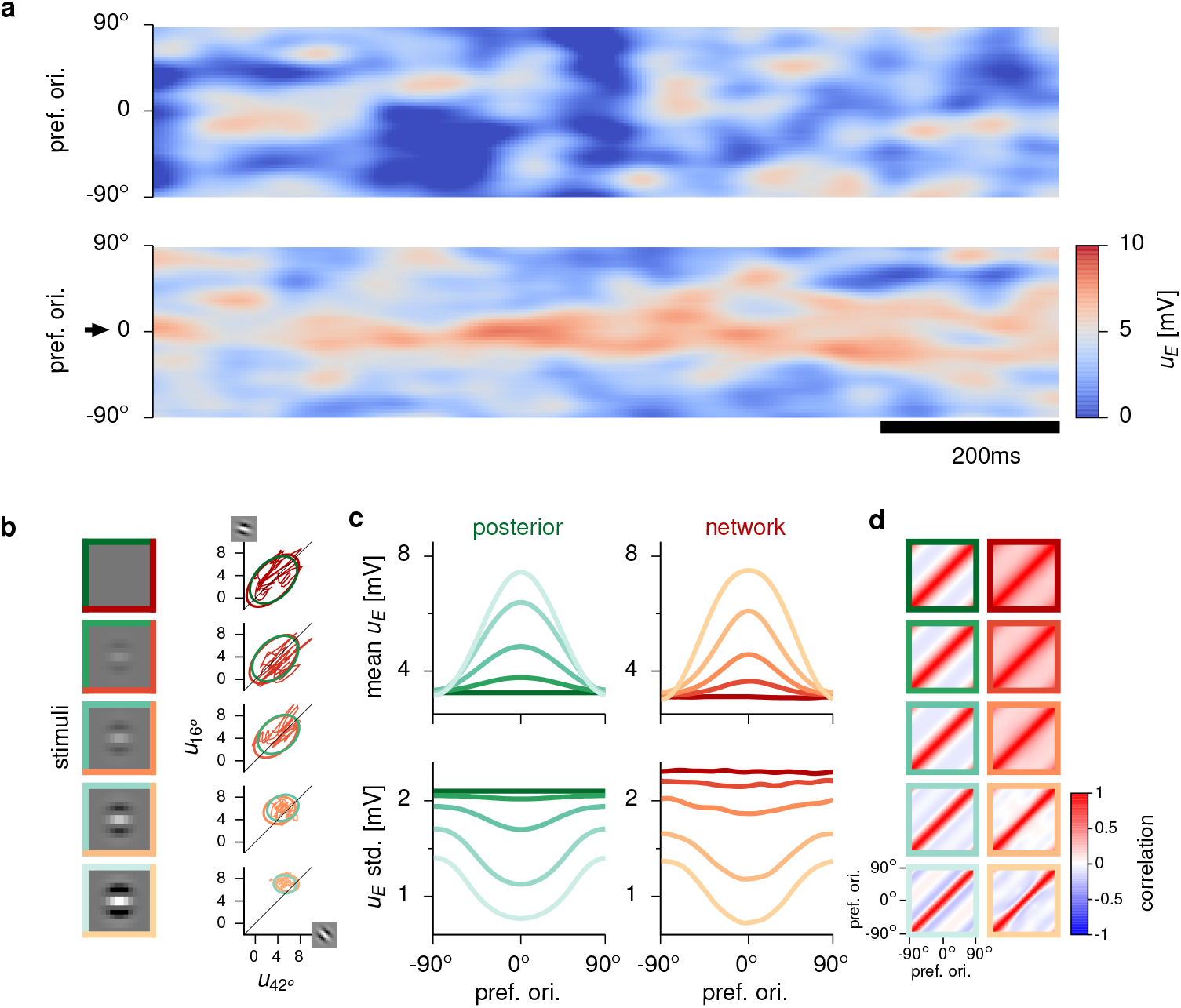
Inference and responses in the optimized network. **a,** Sample population activity of excitatory (E) cell membrane potentials *u*_E_ (color) at zero (top) and high (bottom) contrast. The high contrast stimulus has a dominant orientation at 0° (arrow). Neurons are ordered by preferred orientation. **b,** Left: stimuli in the training set (shade of frame color indicates contrast level, split green and red indicates that the same stimuli were used as input to the ideal observer and the neural network). Right: covariance ellipses (2 standard deviations) of the ideal observer’s posterior distributions (green) and of the network’s corresponding response distributions (red). Red trajectories show sample 500 ms-sequences of activities in the network. Projections for two representative latent variables / E cells are shown, with projective fields / receptive fields at preferred orientations 42° and 16° (insets at the end of axes). **c,** Mean (top) and standard deviation (bottom) of latent variables under the ideal observer’s posterior distribution (left, green) and of E cell membrane potentials **u**_E_ under the network’s stationary distribution (right, red), ordered by their preferred orientation, for each stimulus in the training set. **d,** Correlation matrices of the ideal observer’s posterior distributions (left, green) and the network’s stationary response distributions (right, red). Line colors in **c** and frame colors in **d** correspond to different contrast levels, same colors as stimulus frames in **b**.stimulus-dependent patterns of across-trial mean and covariance.

**Fig. 3.**
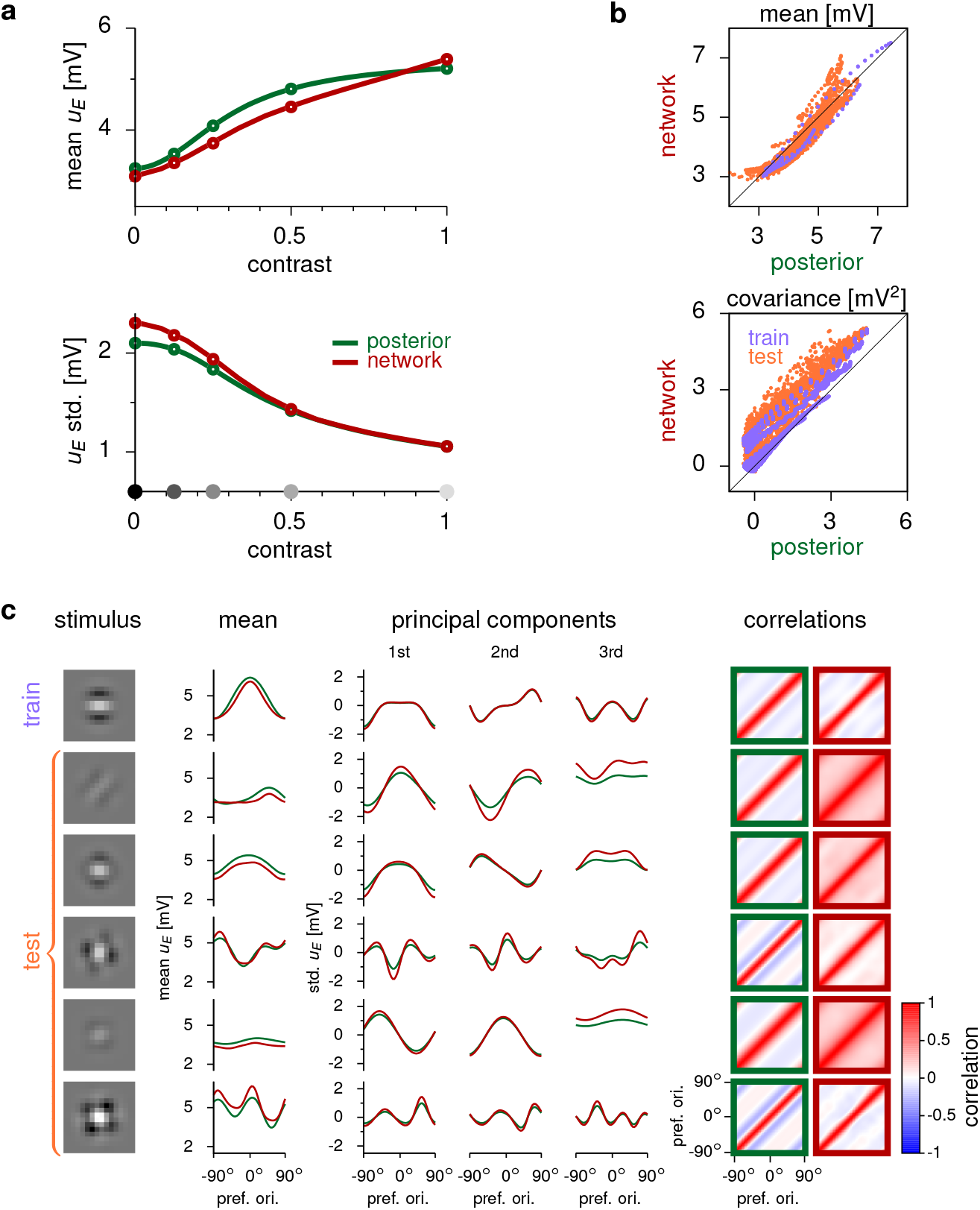
Generalization in the optimized network. **a,** Mean and standard deviation of latent variables (green) and stationary network responses (red) averaged over the population, as a function of contrast. Circles, and gray dots on x-axis indicate training contrast levels. The network correctly generalizes to untrained contrast levels (segments between circles). **b,** Stationary mean (top) and covariance (bottom) during network activity (y-axis) versus under the posterior (x-axis). Each dot corresponds to the response of an individual cell (top) or cell-pair (bottom) to one of the trained stimuli (purple) or one of the novel, untrained stimuli in the test set (orange). **c,** Examples of generalization in the network. Each row corresponds to a different stimulus, and shows the corresponding statistical moments of latent variables under the GSM posterior (green) and stationary responses in the network (red). As a reference, the top row shows one of the training stimuli. The bottom five rows show generalization to novel test stimuli. Left: example stimuli. Middle: GSM (green) and network means (red), and the first three principal components of the GSM covariance, scaled by the square root of the variance they explain of the GSM posterior (green) and of the network covariance (red). Right: correlation matrices of the ideal observer’s posterior distributions (left, green frames) and the network’s response distributions (right, red frames).

While the specific circularly symmetric architecture of the network automatically ensured exact generalization to stimuli that were rotated versions of training stimuli (Online Methods), we also tested the capacity of the network to represent the appropriate posterior distributions for genuinely novel stimuli. First, we employed the same image patch that was used to construct the training set but presented it at novel, intermediate contrast levels. The mean and variability of network responses smoothly interpolated between the corresponding target moments, closely following the behavior of the GSM posterior (Fig. 3a, solid curves between circles). Next, we presented the network with 500 entirely novel image patches randomly generated from the GSM (Online Methods, Fig. S1c). Overall, both the means (Fig. 3b, top) and covariances of network responses (Fig. 3b, bottom) matched those of the target posteriors. This match was similarly good for test stimuli (Fig. 3b, orange) as for training stimuli (Fig. 3b, purple). Critically, while the inputs of the training set included a single dominant orientation, many test stimuli had a more complex structure, with more than one dominant orientation (Fig. 3c, first column). Consequently, the corresponding GSM posteriors that the network was required to match became qualitatively different. Specifically, both the mean activity profiles across the population (Fig. 3c, 2nd column) and the principal components (PCs) of the noise covariances (Fig. 3c, remaining columns) became multimodal and highly dependent on the stimulus (Fig. 3c, green; compare across rows). The network was able to match the required GSM posteriors with high accuracy even in these challenging cases (Fig. 3c, red). Thus, the optimized network performed approximate Bayesian inference over a wide array of stimuli by always sampling (approximately) from the appropriate, stimulus-dependent high-dimensional posterior distribution of the ideal observer.

### The optimized network performs fast sampling

Under sampling-based inference, the time it takes to accurately represent a posterior distribution by collecting successive samples is directly proportional to the timescale over which these samples are correlated^40^. For example, if neural responses were correlated on a 100 ms timescale, it would take on the order of a second to obtain 10 independent samples. In our optimized network, noise variability generated new, independent samples every few tens of milliseconds across all contrast levels. This was evident in its fast-decaying membrane potential autocorrelations (Fig. 4a, colored curves). These timescales were similar to the timescales of activity fluctuations observed in sensory cortical areas^18,41^. In fact, they were faster than what would have been expected in a disconnected network with the same membrane and input time constants (Fig. 4a, dashed curve). Sampling speed was even close to the theoretical limit of a network of infinitely fast neurons in which sampling speed is solely limited by the input time constant (Fig. 4a, dotted curve).

**Fig. 4.**
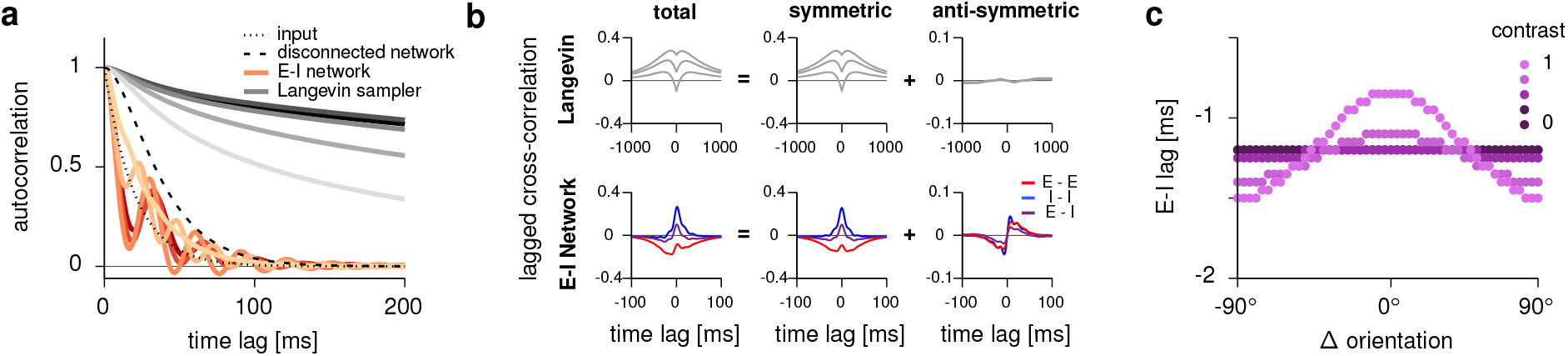
Temporal correlations in the optimized network. **a,** Membrane potential auto-correlations (population average) in the network for increasing levels of stimulus contrast (from dark to pale red; same colors as in Fig. 2b–d). The auto-correlation of a purely feed-forward network (with the same process noise) is shown for comparison (dashed black line), together with those of the process noise (dotted black line), and a collection of networks implementing Langevin sampling at each contrast level (from dark to light gray). **b,** Lagged cross-correlation (left) in the Langevin sampler (top) and in the optimized E–I network (bottom), decomposed into temporally symmetric (middle) and anti-symmetric components (right). Each line corresponds to a different cell pair (3 representative pairs shown), color encodes identity of participating cells (E or I, note that there is no separation of E and I cells in the Langevin networks). **c,** Lag between total E and I inputs to each E cell, as a function of stimulus orientation (relative to preferred orientation) at different contrast levels (colors).

To understand the algorithmic strategy underlying fast sampling in the optimized network, we compared its dynamics to a machine learning algorithm known as Langevin sampling (Online Methods). Langevin dynamics results in a random walk that spends more time in regions of activity space associated with high posterior probability. Sampling by Langevin dynamics was a relevant comparison for our network for two reasons. First, Langevin sampling is a popular, general-purpose algorithm in machine learning and as such can be used to benchmark the performance of our network. Second, most previous work suggested that stochastic recurrent neural networks (without a separation of E and I cells) may implement sampling by Langevin-like dynamics^42–44^. We found that for each input, Langevin dynamics was consistently an order of magnitude slower than our network (Fig. 4a, gray curves). The slowness of Langevin dynamics is known to arise from one of its critical features: time-reversible dynamics— i.e. that any time series of responses is as probable as its time-reversed counterpart^45^. This was indeed reflected in purely temporally symmetric cross correlograms (Fig. 4b, top). In contrast, our optimized network displayed a marked departure from time-reversibility, as evidenced by strong asymmetric components in its crosscorrelograms (Fig. 4b, bottom). Thus, our network fundamentally deviated from most previous proposals for how sampling might be implemented in neural circuits, and achieved an order of magnitude better performance.

The irreversibility of network dynamics implied sequentiality in the activation of particular pairs of neurons. In particular, we found that I cells typically lagged behind E cells. Moreover, for any cell, its total inhibitory input tended to also lag behind its overall excitatory input (Fig. 4c), consistent with known electrophysiology^46^. Interestingly, this lag was smaller for cells that were most strongly driven by the stimulus, and this modulation became stronger with increasing contrast. These form testable predictions of our model.

### Cortical-like dynamics in the optimized circuit

Having established that our network fulfilled its function of representing posterior distributions via sampling, we compared its dynamics with known physiological properties of V1. First, firing rates in the model had a physiologically realistic dynamic range and were tuned to stimulus orientation, similarly to neurons in macaque V1 (Fig. 5a, left-middle; Ref. 8, analysis of data recorded by Ref. 29). We also examined spike count statistics in the network, computed from firing rates assuming a doubly stochastic spike generation process (Online Methods). The quenching of membrane potential variability with increasing contrast that we noted earlier (Fig. 2d, bottom) gave rise to quenching of spike count variability, as quantified by the Fano factor (Fig. 5b, middle; Supplementary Material, Section S3.4). Furthermore, Fano factor suppression was stronger at the cell’s preferred orientation. These effects were qualitatively similar to experimental data from V1, though somewhat weaker in magnitude (Fig. 5b, left). Moreover, stationary responses in the net-work exhibited clear signatures of divisive normalization (Fig. S4), a canonical operation of cortical circuits^47^. All these results were expected for a network whose stationary membrane potential response distributions represent GSM posteriors^6^.

**Fig. 5.**
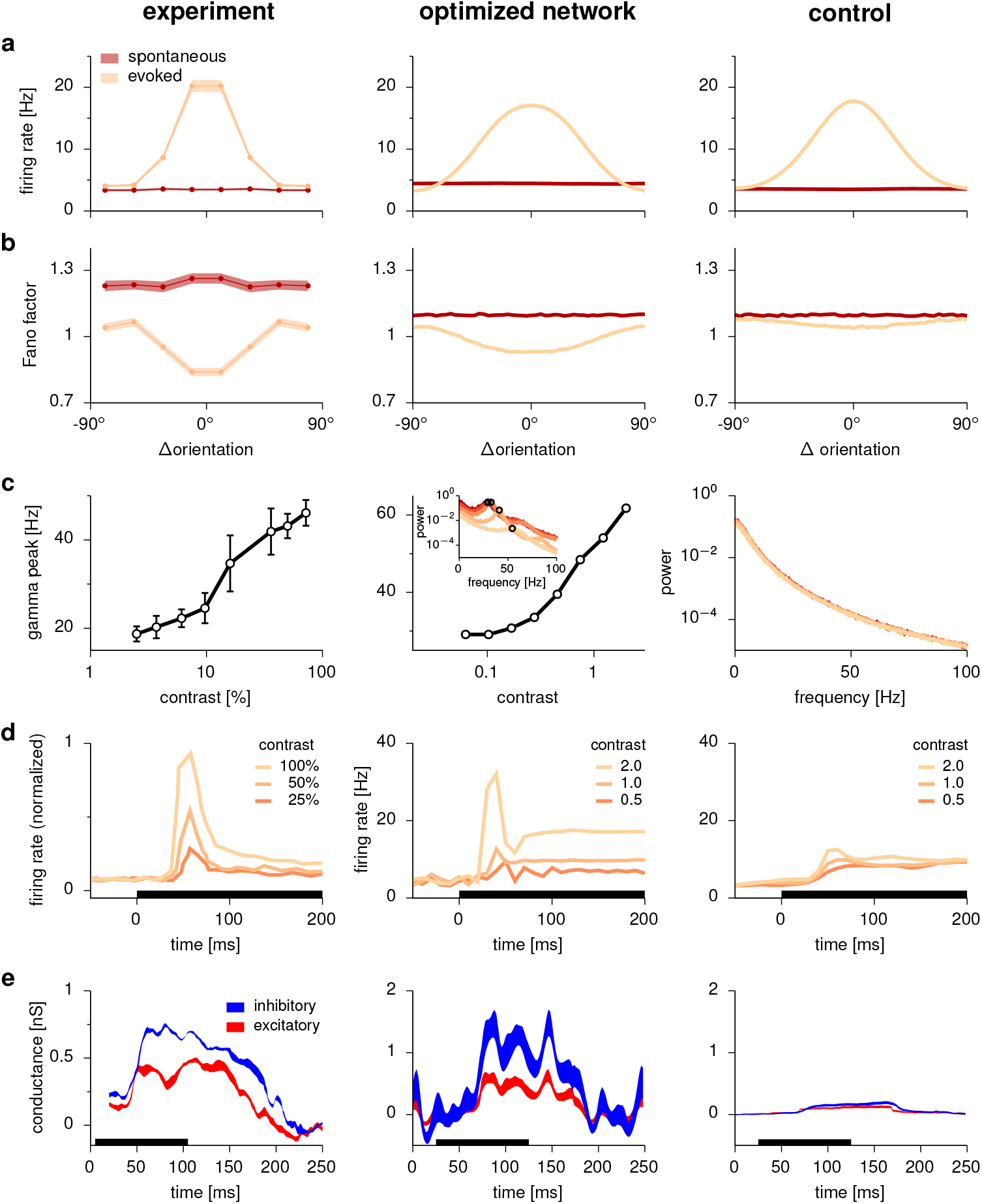
Cortical-like dynamics in the optimized network. Left: experimental data; middle: optimized network; right: control network trained to modulate its mean responses but not its variability. **a-b,** Mean firing rate (**a**) and Fano factor (**b**) of neurons as a function of stimulus orientation (relative to preferred orientation) during spontaneous (dark red) and evoked activity (light orange). Experimental results show mean ± s.e.m. **c,** Peak gamma frequency in the local field potential (LFP) power spectrum as a function of contrast. Inset for the optimized network and main panel for the control network show LFP power spectra at different contrast levels (colors as in Figs. 2 and 4). Note that no dependence of gamma frequency is shown for the control network as there were no discernible gamma peaks in its power spectra. **d,** Average rate response around stimulus onset at different contrast levels (colors). **e,** Excitatory and inhibitory conductance (mean ± s.e.m., relative to baseline, see Supplementary Material, Section S3.4 for details) during a transient stimulus response. Black bars in **d-e** show stimulus period. Panels **a-b** reproduce analyses from Ref. 8 of data from Ref. 29 (awake macaque V1). Experimental data in **c** was reproduced from Ref. 4 (awake macaque V1), in **d** reproduced from Ref. 3 (awake macaque V1), and in **e** reproduced from Ref. 2 (awake mouse V1).

Next, we focused on the dynamical properties of the optimized network. Recall that the optimization procedure only constrained the stationary response distributions and encouraged general temporal decorrelation on fast timescales, but did not otherwise prescribe any specific dynamics. Nevertheless, we found that the network exhibited a number of experimentally observed features of cortical dynamics. Specifically, the optimized circuit displayed strong gamma oscillations, with a peak frequency increasing with contrast, consistent with V1 recordings in the awake monkey^3,4^ (Fig. 5c, left-middle). Moreover, oscillations disappeared entirely when we voltage-clamped each cell in the E population to its mean voltage corresponding to the input (Fig. S5). Analogous voltage-clamp of the inhibitory population led to unstable dynamics (not shown). Thus, in the optimized network, gamma oscillations arose from interactions between E and I cells (i.e. the so-called “PING” mechanism21).

The network also showed strong transient responses such that average population rates had a marked contrast-dependent overshoot at stimulus onset, consistent with recordings in V1 3 (Fig. 5d, left-middle). Finally, using a conductance-based approximation of our current-based model (Supplementary Material, Section S3.4), we found that inhibition transiently dominated over excitation during stimulus presentation, as in V1 of the awake mouse2 (Fig. 5e, left-middle).

We next sought to establish whether these dynamical properties arose simply due to the biological and architectural constraints imposed on our network – or specifically due to the network being optimized for sampling-based inference. For this, we optimized a ‘control’ network in which single cell parameters (time constants and firing rate nonlinearities), overall network architecture, and receptive fields were identical to those used in the original network (Online Methods). Critically, just as the original network, this control network was also trained to match the means of the posterior distributions, but unlike the original network, it was not required to modulate its variability. Despite clear stimulus-dependent modulations in mean responses (as required by training; Fig. 5a, right), the control network exhibited only minimal modulations of both membrane potential variability (Fig. S11) and Fano factors (Fig. 5b, right; Fig. S13). This indicated that modulations of response variability seen in the original network, which are a hallmark of a sampling-based probabilistic inference strategy^6^, were not just a generic by-product of non-linear E–I dynamics^36^. Moreover, neither gamma oscillations nor marked inhibition-dominated transients emerged in the control network (Fig. 5c-e, right). This was still the case when we optimized the network for matching means and variances only, but not covariances as would be required for “full” probabilistic inference (Figs. S10 and S13). Furthermore, oscillations were also absent in a third control network specifically optimized to modulate its mean firing rates as before, while keeping its Fano factors constant (Figs. S12 and S13), as would be required by other, non sampling-based probabilistic representations^17^. These results suggest that the dynamical features observed in the original network could emerge as a consequence of the specific computation for which it was optimized. Conversely, training the network on the original cost function but without enforcing Dale’s principle as a constraint resulted in substantially poorer performance and a lack of oscillations and transients (Figs. S7, S8 and S13). Thus, rather than just being a hindrance, some biological constraints we considered may have actually helped achieving competent performance and thus contributed to the cortical-like dynamical features of our network.

### Oscillations improve mixing time

To study the potential functional benefits of oscillations, one would ideally like to perform a “knockout” experiment with a well-controlled removal of oscillations from the dynamics of the network, leaving all other features of the dynamics intact. Given the complex and high dimensional dynamics of our network, this approach seemed unfeasible. Therefore, we resorted to first studying the response of just a single neuron to obtain an analytical understanding of the general role of oscillations in sampling (Fig. 6a-b). These analyses readily generalized to oscillations in network-wide activity patterns rather than single neurons, thus providing insights into the high-dimensional dynamics of the full network (Fig. 6c-d).

**Fig. 6.**
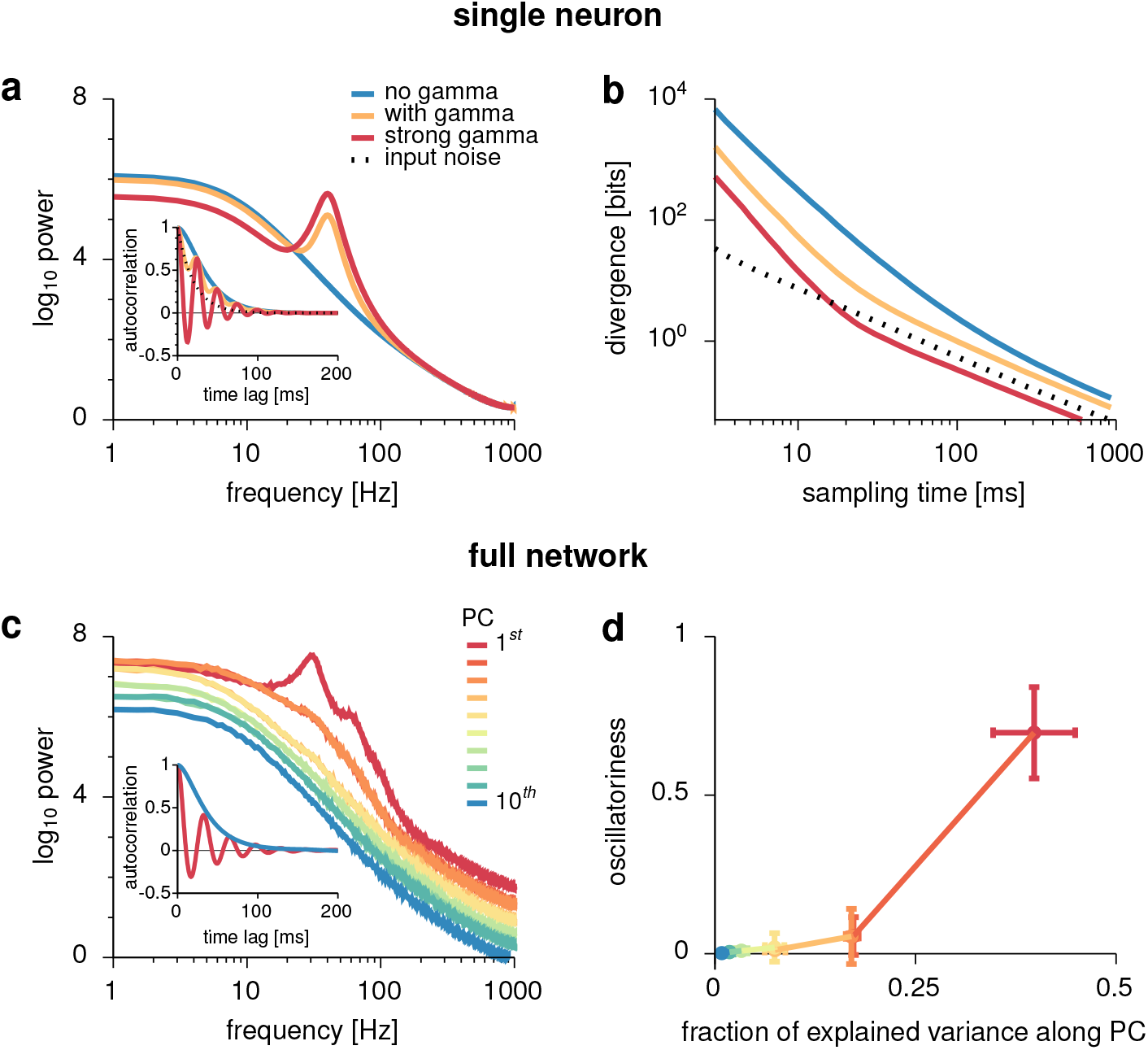
Oscillations improve mixing time. **a-b,** Analysis of oscillations in the response of a single neuron. **a,** Power spectra of three different neural responses (colored lines) with identical mean and variance but different degrees of oscillatoriness. Inset: autocorrelation functions. Black dotted line represents the autocorrelation of the process noise. **b,** Divergence between the distribution estimated from a finite sampling time (x-axis) and the true stationary distribution for the three systems (colors as in **a**). **c-d,** Analysis of oscillations in the full network. **c,** Power spectra of the network’s neural activity along the directions of the principal components (PCs) of its stationary response distribution, ordered by PC rank (colors). Inset: autocorrelation of neural activity along the directions of the 1st and 10th PCs (colors as in main plot). **d,** Oscillatoriness of the autocorrelogram along each principal component (colors as in **c**) as a function of the fraction of the total variance of responses they capture. Note that our measure of oscillatoriness is based on the relative contributions of an oscillatory vs. a non-oscillatory component in a parametric fit to the autocorrelogram, and as such it is invariant to the overall magnitude of fluctuations (which is factored out by using the autocorrelation rather than the autocovariance of responses, Supplementary Material, Section S3.5).

Assuming that the response of a neuron is statistically stationary and approximately normally distributed, it is fully characterized by its mean, variance, and autocorrelogram. As long as this neuron is part of a sampling-optimized network, the mean and variance of its response will be determined by matching the mean and variance of the target distribution sampled by the network (Fig. 2b-d). Although the autocorrelogram is not constrained by the objective of matching these target moments, it still contributes critically to the performance of the network. Specifically, it determines the speed with which the samples produced by the network dynamics become representative of the target distribution. In particular, it can be shown mathematically that the total area under the autocorrelogram directly scales “mixing time”: the time it takes for the dynamics to represent the target distribution to a given precision (Supplementary Material, Section S5.1). Therefore, to understand the specific role of oscillations in the performance of a sampling-optimized network, we compared idealized (stationary and normally distributed) neuronal responses, constructed to have the same mean and variance as responses in our network but different autocorrelation functions. We noted that the envelope of the autocorrelogram of any neuron in the original network would be ultimately constrained by the time constant of the process noise (Fig. 6a, inset, black dotted; see also Fig. 4). We therefore restricted all autocorrelograms in our analysis to have the same envelope as in the full network, and only varied their degree of “oscillatori-ness” (Fig. 6a, blue, orange, red). These oscillations were able to substantially reduce the area under the autocorrelogram (Fig. 6a, inset) and thus reduce mixing time (Supplementary Material, Section S5.2.1). This could be seen by the speed with which the distribution of responses measured over a finite time window converged to the target distribution (Fig. 6b). Importantly, oscillations will only decrease the area under the autocorrelogram, and thus mixing time, if at least one oscillation cycle fits under the envelope. This requires the oscillation period to be sufficiently shorter than the time constant of the envelope (approximately 35 ms), implying oscillations at 30 Hz or higher. Indeed, the lowest frequency we observed in the network was about 30 Hz (Fig. 5c, middle).

We next studied the organization of gamma oscillations in the multidimensional responses of the full network. Previous mathematical analyses suggested that fast convergence to a highdimensional target distribution requires temporally irreversible dynamics, such as those exhibited during oscillations^45^. Here, we were able to show that maximal sampling speed is achieved specifically when smaller response variance is associated with higher oscillation frequency (Supplementary Material, Section S5.2.3). In turn, as we showed above, variability is quenched with increasing contrast both in our network and in the cortex (Fig. 2b-c, Fig. 3a, Fig. 5b). This explains why the frequency of gamma oscillations increased with contrast in our network after optimization. These results suggest that contrast-modulated gamma oscillations observed in the cortex^3,4^ may reflect a speed-optimized sampling strategy (Fig. 5c). The mechanism by which oscillations speed up sampling in the response of a single cell (Fig. 6a-b) generalize to oscillations along any direction of the multidimensional state space of the full network, i.e. network-wide activity patterns. We thus wondered whether there were specific directions in the state space of the full network in which oscillations appeared or if they were distributed evenly across all directions. In line with intuition, our analyses predicted that oscillations in an efficiently sampling network should be predominantly expressed where they matter most: in the (stimulus-dependent) network-wide activity patterns capturing most of the overall response variability. Namely, for each stimulus, we expected the strongest oscillations along the top PCs of the corresponding stationary covariance (Supplementary Material, Section S5.2.4). This was indeed apparent in the power spectra of our network associated with the top 10 PCs (Fig. 6c), and the corresponding autocorrelograms that even showed negative-going lobes (Fig. 6c, inset). Specifically, there was a positive relationship between oscillatori-ness along successive PCs and the fraction of variance explained. This meant that the network oscillated more in the directions along which its responses had the largest variance (Fig. 6d). (Note that our measure of oscillatoriness was based on autocorrelograms, and therefore had no a *priori* dependence on response variance; Supplementary Material, Section S3.5.) In sum, the network used non-trivial temporal dynamics, in the form of contrast-dependent, pattern-selective gamma oscillations, to ensure that even short segments of its activity were sufficiently representative of the posterior distribution it represented for each stimulus.

### Transients support continual inference

The foregoing results showed that oscillations increased the effective speed of sampling in the stationary regime, i.e. once network responses had become representative of the target distribution (also known as “mixing speed”40). Complementing this, we found that transients in our network mitigated the other main temporal constraint of sampling: the so-called “burn-in” time it takes for responses to become representative in the first place^40^. We observed that, in line with experimental data, during stimulus onset, neural responses tended to overshoot the corresponding stationary response levels (Fig. 5d, Fig. 7a). One might naively expect such transients to be detrimental for representing a distribution, as they clearly deviate from the target (represented by the steady-state responses). However, in a realistic setting with a changing environment, distributions need to be represented continually, without waiting for the system to achieve steady state. Thus, we considered how a moving decoder of neural responses over a finite trailing time window approximated the target.

**Fig. 7.**
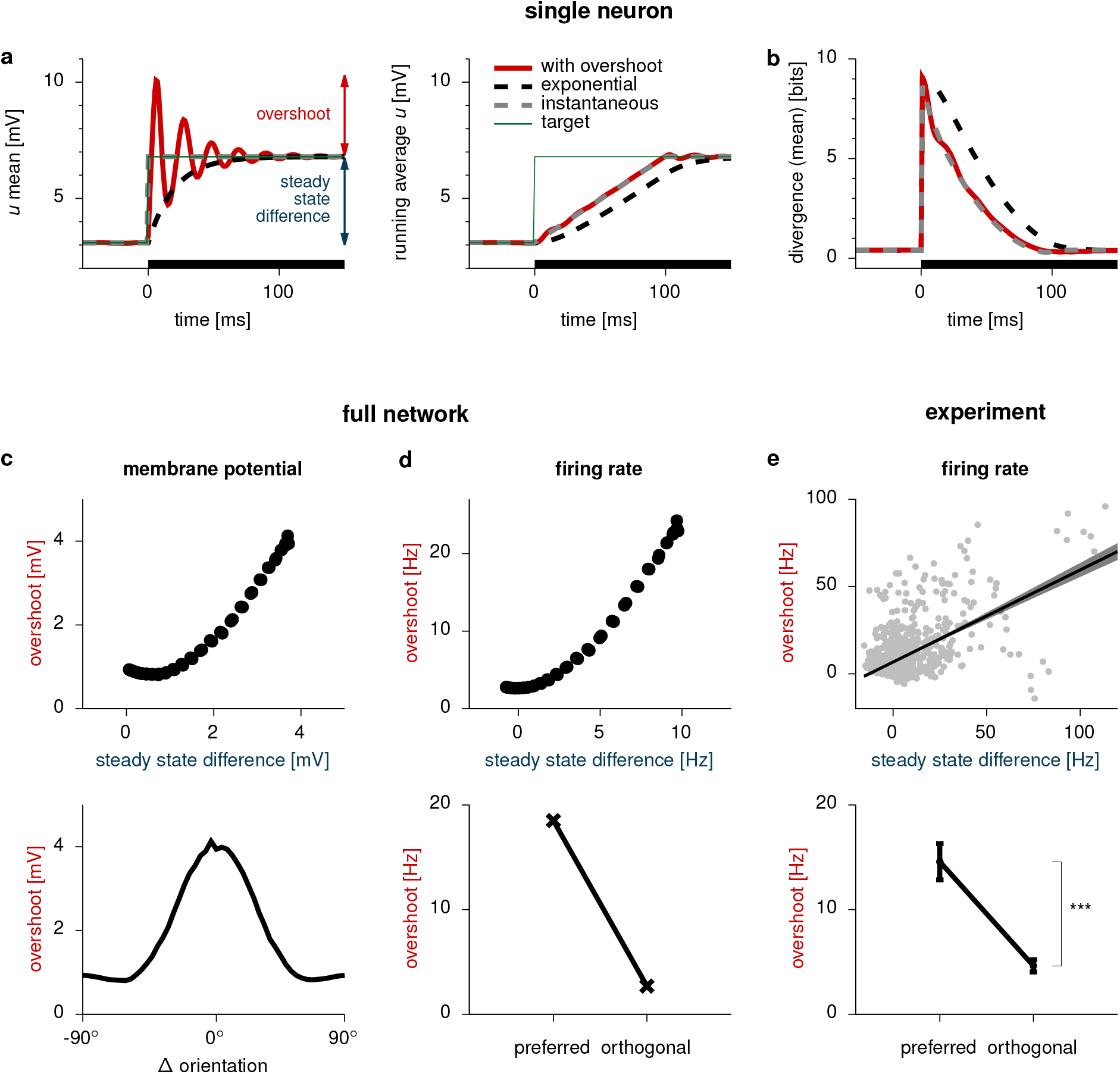
Transients support continual inference. **a-b,** Analysis of transients in the response of a single neuron. **a,** Temporal evolution of the mean (left) membrane potential (*u_E_*), and its running average (right), in three different neural responses (thick lines) with identical autocorrelations (matched to neural autocorrelations in the full network, Fig. S15a; cf. Fig. 6) but different time-dependent means (shown here) and variances (Fig. S15a). Thin green line shows the time-varying target mean. **b,** Divergence between the target distribution at a given point in time and the distribution represented by the neural activity sampled in the preceding 100 ms, for each of the three responses (colors as in **a**). The mean-dependent term of the divergence is shown here, which depends on the difference between the target mean and the running average of samples (shown in **a**, right; see Fig. S15b for the full divergence). Black bars in **a-b** show stimulus period. **c,** Top: overshoot magnitude versus steady state difference in membrane potentials (see **a** for legend). Each dot corresponds to the response of one cell to one particular stimulus. Bottom: overshoot magnitude as a function of stimulus orientation (relative to preferred orientation). **d,** Top: same as **c**, top, for firing rates. Bottom: average rate overshoot across cells whose preferred orientation is aligned with the stimulus (0 ± 30°), or near-orthogonal to it (90 ± 30°). **e,** Analysis of experimental recordings from awake macaque V1^29^. Top: overshoot magnitude versus steady state difference, as in **d**, top. Black line shows linear regression (±95% confidence bands); ***: *p* < 0.001 (**n** = 1280 cell-stimulus pairs; see also Fig. S15d-e). Bottom: average (±1 s.e.m) overshoot magnitude for preferred and orthogonal stimuli, as in **d**, bottom; ***: **p** < 0.001 (**n** = 864 cell-stimulus pairs).

As with oscillations, we performed this analysis in two steps. First, to isolate the potential functional benefits of transients, beyond those of oscillations we analyzed before, we once again considered the response of just a single idealized neuron that is part of a sampling-optimized network (Online Methods; Supplementary Material, Section S5.3.1). For this idealized neuron, we fixed the autocorrelogram as well as the before- and after-stimulus onset steady-state means and variances to those of an actual, representative neuron in our network. We then studied different ways in which this neuron could transition between these two steady states (Fig. 7a). We considered three possibilities: 1. as an upper bound on performance, instantaneously switching between the two steady-states (Fig. 7a, gray dashed); 2. exponentially approaching the new steady state with the characteristic time constant of the cells in the network, thus lacking overshoots (Fig. 7a, black dashed); and 3. undergoing overshoots as seen in our optimized network (Fig. 7a, red). We found that overshoots performed close to the upper bound, provided by instantaneous switching. In particular, they generated samples that allowed a substantially more accurate estimate of the target mean than that afforded by approaching the new steadystate exponentially without overshoots (Fig. 7a– b). (These results extended qualitatively to the case when the match in the full distributions was considered, Fig. S15b.) This was because without overshoots at stimulus onset, responses were still sampling from the distribution corresponding to the baseline input. Thus, including them in the estimation of the new stimulus-related mean inevitably biased the estimate to be too low. This bias was largely offset by the overshoot. Indeed, we were able to show analytically that the optimal way to compensate for this bias was to express transient overshoots at stimulus onset (Supplementary Material, Section S5.3.2, Fig. S15c). The intuition for this is that continual averaging of responses formally corresponds to a temporal convolution, and so the optimal response was the deconvolution of the target with the averaging kernel. Under basic smoothness constraints, the deconvolution of a step function with such an averaging kernel yields similar transients to those that we observed in the network (Fig. S15c).

The hypothesis of increased sampling accuracy by transient compensation made a distinct prediction (which was also supported by our mathematical analysis, Supplementary Material, Section S5.3.2). Namely, transient overshoots should scale with the change in steady state responses. Indeed, our network exhibited this effect in both membrane potentials and firing rates (Fig. 7c-d, top). Importantly, this also resulted in transients being orientation tuned, reflecting the tuning of stationary responses (Fig. 7c-d, bottom). While stimulus-onset transients have been widely observed^3,4^, previous reports did not analyze their stimulus tuning. Therefore, we conducted our own analyses of a previously published dataset of V1 responses in the awake monkey^29^. In line with the predictions of the model, the size of overshoots were orientation tuned (Fig. 7e, bottom) and, more generally, they scaled with the change in stationary responses *(n =* 1280 stimulus-cell pairs, coefficient of determination *R*^2^ ≃ 0.33, *p* < 0.001, Fig. 7e, top; these results were robust to excluding the outliers with high firing rates, e.g. above 60 Hz: *n* = 1263, *R*^2^ ≃ 0.27, *p* < 0.001, Fig. S15d-e).

## Discussion

We have shown that a canonical neural network model^8,35^ produces cortical-like dynamics when optimized to perform sampling-based probabilistic inference, but not when optimized to perform a non-probabilistic objective, or a non sampling-based probabilistic objective. Further controls demonstrated that these dynamics were not mere side products of the particular biological constraints or optimization approach we adopted. Instead, they played well-defined functional roles in performing inference.

### The Gaussian scale mixture model and the stochastic stabilized supralinear network

We used a canonical model of neural network dynamics (the stochastic SSN) to embody a set of biologically relevant constraints for cortical circuits. It was not trivial *a priori* that this model would be able to modulate its responses as necessary for successful sampling-based inference under a canonical generative model of visual image patches (the GSM). A hint that this might indeed be possible came from previous studies showing that both in the SSN^8^,35 and the GSM^6,48^, a range of parameters exists for which the average response or posterior mean monotonically increases while the variance decreases with increasing stimulus strength. Empirically, we found a good quantitative match that went beyond this coarse, qualitative trend, with the SSN also capturing much of the detailed structure of the GSM posteriors. However, this match was not perfect: for example, the GSM posteriors systematically showed negative correlations of larger magnitude than what the network was able to express (Fig. 2d and Fig. 3b-c). It might be possible to achieve a more accurate match by allowing negative input correlations, and in general a more flexible parameterization of the SSN. Nevertheless, basic information theoretic considerations suggest that the decrease of posterior variability with contrast can be expected to be a hallmark of any statistically efficient model of natural images, not just of the highly simplified (ring-structured) GSM we used here^18^. Indeed, the same qualitative behaviour was found earlier using a richer GSM model of V1 with a greater variety of basis functions^6^. Thus, once the optimization of larger-scale SSNs becomes feasible, we expect them to also show this behaviour.

Interestingly, the divisive normalization performed by the SSN has been proposed to be a canonical operation implemented throughout the cortex^47^. At the same time, stacked layers of subunits, each with a GSM-like separation of content- and style-like variables, have been suggested to form the basis of probabilistic generative models underlying deep learning for high-nuisance inference tasks^49^. Therefore, our results establishing the SSN as an appropriate approximate inference “engine” (a so-called recognition model) for the GSM suggest that, similarly, a cascade of circuits with SSN-like dynamics could perform efficient inference under more powerful generative models and thus account for computations beyond V1^50^.

### Function-optimized neural networks

Our approach is complementary to classical approaches for training neural network models that had shown how various steady-state properties of cortical responses (such as receptive fields, or trial-averaged activities) emerge from optimizing neural networks for some computationally well-defined objective (such as object recognition, memory, context-dependent decision making, or sensorimotor control)^39,51–55^. Notably, our sampling-based computational objective required our network to modulate not only the mean but also the variability of its responses in a stimulus-dependent manner. This made the training of networks significantly more challenging than conventional approaches training networks for deterministic targets without explicitly requiring them to modulate their variability^51,54,56,57^. In return, the dynamics of our network exhibited rich, stimulus-modulated patterns of variability. These responses captured a variety of ubiquitous features of the trial-by-trial behaviour of cortical responses (noise variability, transients, and oscillations) beyond the steady-state or trial-average properties that could be addressed by previous work.

Typically, previous network optimization approaches aimed to determine the types of dynamics that arise when a task is executed under minimal mechanistic constraints, using a neural network as a universal function approximator. As a result, they yielded fundamental insights about the macroscopic organization of network dynamics (e.g. the presence of line attractors57) but did not attempt to incorporate some of the most salient constraints on the detailed organization of cortical circuits. Specifically, they used networks that were either purely feed-forward^51^, utilized neuronal transfer functions that lacked the expansive nonlinearities characteristic of cortical neurons^51,53,55–57^, had no separation of E and I cells^51,55–57^, or had noiseless dynamics^56^.

In contrast, our goal was to study the emergence of (probabilistic) computations through dynamics and to connect these dynamics to experimental data at (or near) single cell resolution (e.g. the neuron- and stimulus-specific reduction of variability, or the lag between total inhibitory and excitatory inputs in individual cells). This required respecting all the aforementioned biological constraints. Nevertheless, this additional realism came at the cost of having to limit the number of optimized parameters to be far lower than standard approaches with feedforward networks or recurrent networks for which dynamical stability is more easily achieved. While this reduced parametrization made it easier to find stable solutions, it was still sufficiently expressive. In particular, we found that our results could not have been obtained without optimization (Figs. S6 and S7), or with optimizing other objective functions (Figs. 5 and S10-S13). Indeed, this parametrization still included networks that were unstable, or showed a decrease in mean responses and/or increase in variability with increasing stimulus strength (i.e. the opposite of what was required for matching the GSM), or were modulated in a non-monotonic way or only minimally altogether Fig. S6).

### Neural representations of uncertainty

Our approach markedly differed from previous work on the neural bases of probabilistic inference. Previous models were typically derived using a *top-down* approach (but see Refs. 54,58), using hand-designed network dynamics that explicitly mimicked specific existing approximate inference algorithms from machine learning based on sampling^42–45,59^ or other representations^17,25,42,60,61^. As a result, these models came with strong theoretical guarantees for their performance but often offered only a mostly phenomenological match to neural circuit dynamics. In particular, they did not respect some basic biological constraints (e.g. Dale’s principle^42,44,60,61^), or had to assume an unrealistically rapid and direct influence of stimuli on network parameters (e.g. synaptic weights^44,59^). In contrast, we used a more *bottom-up* approach, starting from known constraints of cortical circuit organization, and then optimizing the parameters of networks under such constraints to achieve efficient sampling-based probabilistic inference – without pre-specifying the details of the dynamics that needed to be implemented. While this approach cannot provide formal guarantees on performance, our optimized network “discovered” novel algorithmic motifs (oscillations and transients) for speeding up probabilistic inference. Although some of these motifs have been observed in previous work^59^, their function remained unclear as they were built-in by design rather than obtained as a result of optimization, or appeared purely epiphenomenal. In contrast, these motifs served computationally well-defined functions in our network.

The dynamics of our network may also provide useful clues for constructing novel machine learning algorithms. In general, the kind of time-irreversible, out-of-equilibrium dynamics we demonstrate for our network have only recently been appreciated in machine learning^28,45^. At the same time, sampling-based inference algorithms using second-order dynamics with so-called “momentum” variables, such as Hamiltonian Monte Carlo, have long been known to improve sampling speed^62^. Indeed, it might be interesting to explore how much the dynamics of our network can be interpreted as a neural implementation of Hamiltonian Monte Carlo^59^. Nevertheless, despite such second-order dynamical systems often exhibiting oscillations and transient overshoots, their sampling efficiency has usually been analyzed only in the more generic terms of the suppression of random walk-like behavior. In contrast, our analyses revealed specific roles for oscillations and transients. In fact, the setting of continual inference that we used to demonstrate the benefits of transients has not been considered in machine learning applications so far, although we expect it to be highly relevant for both biological and artificial cognition.

### Cortical variability, transients, and oscillations

Our work suggests a novel unifying function for three ubiquitous properties of sensory cortical responses: stimulus-modulated variability, transient overshoots, and gamma oscillations. In previous work, these phenomena have traditionally been studied in isolation and ascribed separate functional roles that have been difficult to reconcile. In particular, they have not been derived normatively, i.e. by starting from some functional objective and then optimizing that objective in a principled manner (but see e.g. Ref. 60). For example, cortical variability has most often been considered a nuisance, diminishing the accuracy of neural codes^16^. Theories postulating a functional role of variability in probabilistic computations have only considered the steady-state distribution of responses without making specific predictions about their dynamical features^6,17,63^. Conversely, transient responses prominently feature as central ingredients of models of predictive coding, where they signal novelty or deviations between predicted and observed states^60^. However, these theories did not address response variability.

Our work accounts for both transients and variability starting from a single principle, using only the equivalent of “internal representation neu-rons”64 of predictive coding but without invoking specific prediction error-coding neurons. In particular, our model correctly predicted a specific scaling relationship between transients and steady-state responses which we tested by novel analyses of experimental data (Fig. 7). Furthermore, our mathematical analysis suggested that prediction-error-like signals (more formally, responses that scale with the magnitude of change in the target distribution; Fig. S15c) are a generic signature of continual inference using sampling-based dynamics, and will thus not only appear at stimulus onsets but in any situation when predictions change temporally. A conclusive test of whether prediction-error-like responses in the cortex are due to this mechanism or classical predictive coding mechanisms will require more specific manipulations of prior expectations.

Gamma oscillations have also been proposed as a substrate for a number of functional roles in the past, related to how information is encoded, combined, or routed in the brain^11–14,65^. These putative functions need not be mutually exclusive to that played in our network. Nevertheless, some of these functions seem difficult to reconcile with specific experimental findings^3,19,20,66,67^. More generally, theories of gamma oscillations do not typically address transients. Although there are extensions of the predictive coding framework that do account for the presence of gamma oscillations, by attributing to it the representation of prediction errors^68^, these theories would also predict a tight coupling between gamma-band synchronization and firing rates (both related to prediction errors) which has not been confirmed experimentally^69^. Moreover, it is unclear whether these theories would also account for properties beyond the mere existence of gamma oscillations. These would include the frequency modulation by contrast^3,4^ that our model reproduced (Fig. 5), or indeed any aspect of the ubiquitous variability of cortical responses, and its modulation by stimuli, which our model also reproduced as a core feature (Figs. 2, 3 and 5). In contrast, our results show that variability, transients, and gamma oscillations can all emerge from the same functional objective: that neural circuits use an efficient sampling-based representation of uncertainty under time constraints.

The mechanism by which gamma oscillations are generated in the brain, particularly whether it involves interactions between E and I cells (‘PING’ mechanism) or among I cells only (‘ING’ mechanism), is a subject of current debate^21^. In our model, voltage-clamping of E cells eliminated gamma oscillations (Fig. S5), pointing to the ‘PING’ mechanism. However, our network only included a single inhibitory cell type, and heavily constrained connectivity, therefore it remains for future work to study how the precise mechanism of gamma generation depends on such architectural constraints.

### Hierarchical processing

Our model showed salient divisive normalization (Fig. S4) that is thought to underlie many non-classical receptive field effects in V1^47^. Yet, it only represented a single idealized V1 hypercolumn and thus could not address layer-and cell-type specific lateral and top-down processing which, for example, predictive coding models can more naturally capture^70^. In particular, recent experimental data shows (movement related) mismatch signals in layer 2/3 of mouse V1^71^, suggesting that these neurons may specifically represent errors between bottom-up visual input and top-down predictions. It will be interesting to see whether such neurons would also “automatically” emerge via our approach when optimizing a more complex architecture than what we used here, or if they would require special design constraints or decisions. More generally, a combination of sampling-based representations and predictive coding may be possible^72^. This could lead to computationally powerful representations that both encode uncertainty (which most predictive coding models ignore) and are suited for hierarchical processing (which many models of probabilistic representations eschew). Such a hybrid architecture might be able to account for specific forms of variability modulation by stimuli or topdown signals (at which sampling-based models excel^6,26,45^) as well as for prediction error-like signals (naturally captured by predictive coding models^60,70^).

Studying more hierarchical or spatially extended versions of our model may also allow us to study longer-range aspects of gamma oscillations, such as gamma synchronization^65^, and the dependence of gamma power on the structure of the stimulus at larger spatial scales^73^, which our model of a local hypercolumn could not address. If local features encoded by neurons in different hypercolumns form parts of the same higher-order feature, one expects these neurons to show correlated variability under a sampling-based representation^27^. This in turn may lead to the synchronization of gamma oscillations at their respective sites.

Finally, a fully hierarchical version of our model, including layers that directly control decisions or actions, could also allow end-to-end training for behaviorally relevant tasks, rather than the interim goal of representing uncertainty that we used here. This in turn would make it possible to evaluate whether variability is still used to represent uncertainty despite lacking an explicit objective for doing so. Preliminary results on unsupervised training suggest that this may be the case^58^. By providing a direct read-out of predicted behavior, such hierarchical networks will be ideal to study the link between the temporal dynamics of neural and behavioral forms of variability.

## Acknowledgements

This work was supported by the Wellcome Trust (New Investigator Award 095621/Z/11/Z and Investigator Award in Science 212262/Z/18/Z to M.L., and Seed Award 202111/Z/16/Z to G.H.), and the Human Frontiers Science Programme (Research Grant RGP0044/2018 to M.L.). We are grateful to A. Ecker, P. Berens, M. Bethge, and A. Tolias for making their data publicly available, to G. Orbán, A. Bernacchia, R. Haefner, and Y. Ahmadian for useful discussions, and to J.P. Stroud for detailed comments on the manuscript.

## Author Contributions

R.E., G.H. and M.L. designed the study. R.E. and G.H. developed the optimization approach, R.E. ran all numerical simulations, R.E. and G.H. analyzed experimental data, all authors performed analytical derivations, R.E. G.H. and M.L. interpreted results and wrote the paper, with comments from L.A.

## Online methods

### The generative model

Following Refs. 6,31, we adopted the Gaussian scale mixture model (GSM)30 as the generative model of natural image patches under which the primary visual cortex (V1) performs inference. Thus, an image patch **x** ∈ ℝ^*N*_x_^ was assumed to be constructed by linearly combining a set of local features, the columns of **A** ∈ ℝ^*N*_x_×*N*_y_^, weighted by a set of image patch-specific feature coefficients, **y** ∈ ℝ^*N*_y_^, and scaled by a single global (at the scale of the image patch) contrast variable, *z* ∈ ℝ, plus additive white Gaussian noise, resulting in the following likelihood for the feature coefficients **y**:

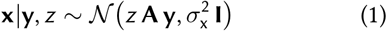

where the feature coefficients were assumed to be drawn from a multivariate Gaussian distribution (i.e. the prior of **y**):

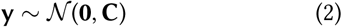

and *z* was assumed to be drawn from a Gamma prior: *z* ~ Γ(*K*, *ϑ*) (Table S1, see also Ref. 48).

To model inferences in a V1 hypercolumn, we chose the columns of **A** (the so-called projective fields of the latent variables) to be oriented Gabor filters that only differed by their orientation (evenly spaced between −90° and 90°, four examples are shown in Fig. 1a, see also Fig. S1a).The prior covariance matrix **C** was a circulant matrix whose elements varied smoothly as a function of the angular distance between the orientations of the projective fields of the corresponding latent variables, with positive and negative correlations between latent variables with similarly and orthogonally oriented projective fields in **A**, respectively (Fig. S1b).

The ideal observer’s posterior over latent spatial features **y** under the GSM for a given image patch, **x**, and a known contrast *z*, can be written as^48^:

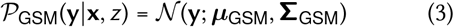

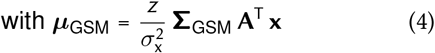

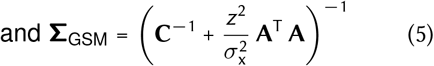

Although, in general, *z* would also need to be inferred, as *z* is just a single scalar, of which the inference pools information across all pixels in the input, we approximated the posterior over *z* with a delta distribution at *z*\*, the true value of *z* that was used to generate the input^48^. Thus, the final posterior over **y**, after marginalizing out the unknown *z*, was approximated by substituting *z*\* into Eq. 3:

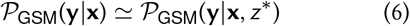

Following Ref. 6, membrane potentials, **u**, were taken to represent a weakly non-linear function of visual feature activations **y** (Supplementary Material, Section S1):

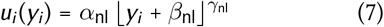

where ⌊·⌋ is the threshold-linear function, and *α*_nl_, *β*_nl_, and *γ*_nl_ are respectively the scaling, baseline, and power of the transformation (Table S1, Fig. S2a).

### Network dynamics and architecture

Our nonlinear, stochastic E/I network consisted of *N*_E_ excitatory and *N*_I_ inhibitory neurons. Following Ref. 8, we modelled the dynamics of each neuron *i* as:

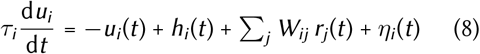

where *u_i_* represented the membrane potential of neuron *i, τ_i_* was its membrane time constant, *h_i_* its feedforward input, *η_i_* was process noise (incorporating intrinsic and extrinsic forms of neural variability), and *W_ij_* was the weight of the synapse connecting neuron *j* to neuron *i*. Firing rates *r_i_* were given by a supralinear transformation of membrane potentials:

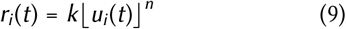

where *k* and *n* were respectively the scale and exponent of the firing rate nonlinearity *(Table S1)*. We reasoned that any network performing accurate sampling-based inference under our ring-structured GSM would need to exhibit the same circular symmetry. We therefore parametrized the recurrent connectivity of the network to be rotationally symmetric, such that neurons were arranged into pairs of E and I cells around a “ring” according to their preferred orientations *(Fig. 1c*) and the connectivity of the network (as well as the process noise covariance, see below) was a smoothly decaying (circular Gaussian) function of the tuning difference between two cells. Specifically, each quadrant of the weight matrix (E → E, E → I, I → E, and I → I) was defined as:

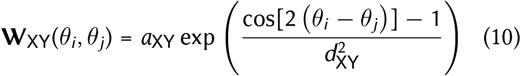

where *X, Y* ∈ {E; I} and *θ_i_* = *πi*/*N*_E/I_ was the orientation represented by the *i*^th^ E/I neuron. Thus, we did not optimize all elements of the weight matrix, but only the eight free parameters *a*_XY_ and *d*_XY_. We also constrained the *a*_XY_ amplitudes to be positive for *Y* = E and negative for *Y* = *I*, such that the network obeyed Dale’s principle. This circulant parametrization implied that training the network on one particular stimulus-posterior pair in effect trained the network on all possible rotations of this pair. This reduced the size of the training set necessary to achieve good generalization, and therefore sped up training.

Neurons in the network also received stimulus-independent process noise that was spatially and temporally correlated zero-mean Gaussian (e.g. modelling inputs from other brain areas, or intrinsic variability in the network):

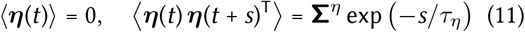

where *τ_η_* was the timescale of the process noise (Table S1) and **Σ**^*η*^ was the stationary (zero-lag) covariance matrix parametrized block-wise as:

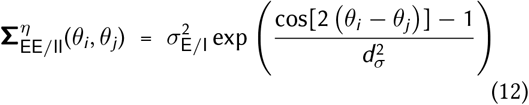

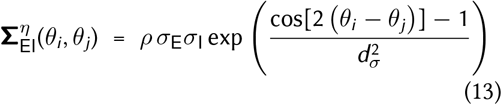

which introduced four additional free parameters: *σ*_E_ > 0, *σ*_I_ > 0, *ρ*, *d_σ_*.

As in standard models of V1 simple cells^74^, the stimulus-dependent input to each neuron was obtained by applying a linear filter **W**_ff_ to the stimulus followed by a static nonlinearity:

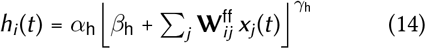

where **x**(*t*) was the stimulus (input image patch) received at time *t*, and *α*_h_, *β*_h_, and *γ*_h_ were respectively the scale, baseline, and exponent of the input nonlinearity (Table S1, Fig. S3b). Given the one-to-one correspondence between the latent variables of the GSM and excitatory-inhibitory neuron pairs of the network model (Fig. 1a and c), we determined the external input to each neuron via an input receptive field that was identical (up to a constant factor) to the projective field of the corresponding GSM latent variable, as this was suggested to be optimal for sampling by previous work^39^: **W**^ff^ = [**A A**]^T^/15, where **A** was the same matrix as in the generative model (Eq. 1), and [**A A**] denotes concatenating **A** with itself column-wise.

In summary, we optimized a total of 15 parameters: 8 describing the weight matrix **W** (Eq. 10), 4 describing **Σ**^*η*^ (Eqs. 12 and 13), and 3 specifying the mapping from stimuli to network inputs (Eq. 14).

### Training and test stimuli

The training set (Fig. 2b) consisted of five image patches, 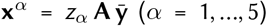 with the same dominant orientation (defined by 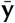, Supplementary Material, Section S2.1) but different contrast levels (*z_α_*), together with their corresponding target moments, which were the moments of the posterior over **u** as prescribed by the generative model (Eqs. 3–7):

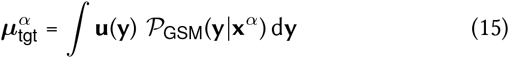

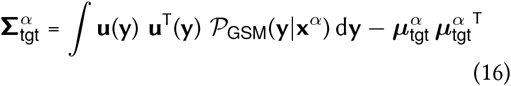

To test generalization in the network, we generated a set of 500 novel image patches with the GSM, which were thus not constrained to have a single dominant orientation (as the prior allowed multiple elements of **y** with different projective fields to be non-zero, Eq. 2). To be consistent with the training set, we did not include additive noise in **x**, and added a contrast-dependent baseline to **y** so that its mean was modulated by contrast in the same way as in the training set. For each image patch in the test set, we also computed the corresponding posterior moments (Eqs. 15 and 16) to evaluate the network’s test performance.

### Network optimization

The cost function 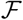 for which we optimized the network consisted of four terms for each input stimulus *α* in the training set:

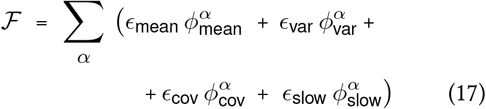

The first three terms of Eq. 17 penalized differences between the (across trial) moments of the network’s response distribution (mean, ***μ**^α^*(*t*), variance, ***σ**^α^*(*t*), and covariance, **Σ**^*α*^(*t*)) averaged over a finite time window ending at *T*_max_ = 500 ms after stimulus onset, and the respective target moments of the ideal observer’s corresponding posterior distributions (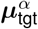 and 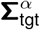 from Eqs. 15 and 16, and with 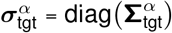),:

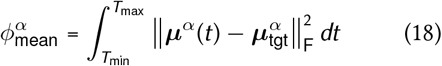

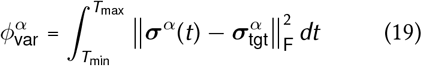

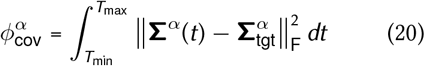

The last term of Eq. 17 was an additional slowness cost, penalizing the total lagged neural response autocorrelation, given by the diagonal of **C**(*τ*) = corr(**u**(*t*), **u**(*t + τ*)), within a *τ*_max_ = 100 ms time window:

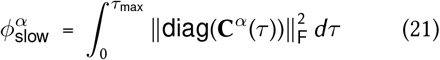

The coefficients *ϵ* controlled the relative importance of these terms (Table S1). The beginning of the averaging window, *T*_min_, was systematically changed (“annealed”) over optimization from *T*_min_ = 0 ms (stimulus onset) to *T*_max_ – 50 ms (Supplementary Material, Section S2.3). The finite length of the averaging window, and in particular including samples immediately or shortly following stimulus onset, encouraged fast sampling. Thus, setting the explicit slowness cost *ϵ*_slow_ = 0 did not qualitatively affect our results (Figs. S9 and S13). In the first control network (Fig. 5, right column; Figs. S11 and S13), we set *ϵ*_var_ = *ϵ*_cov_ = *ϵ*_slow_ = 0, but kept all other metaparameters and target means the same. In the second control network (Figs. S10 and S13), we set only *ϵ*_cov_ = *ϵ*_slow_ = 0, but left *ϵ*_var_ and other meta-parameters the same as in the original network. In the third control network (Figs. S12 and S13, right), all *ϵ*… parameters were the same as for the optimization of the original network, but the target covariances were modified to induce contrast-independent Fano factors (see Supplementary Material, Section S4.4).

To minimize the cost function in Eq. 17, we used a novel combination of stochastic and deterministic methods, both involving back-propagation through time75 (Supplementary Material, Section S2.2). The optimizer was written in OCaml and can be found online at bitbucket.org/RSE_ 1987/ssn_inference_optimizer.

As the cost function that we used (Eq. 17) was non-convex, we checked the robustness of our findings by performing 10 further optimization attempts from random initial conditions. No solutions achieved substantially lower costs, and those whose final cost was at least approximately as low as the network presented in the main text behaved qualitatively similarly (in particular, they showed contrast-dependent oscillations and transients, Fig. S7). Nevertheless, our results should not be taken to represent a global optimum of our cost function.

### Numerical experiments after training

To obtain a reliable estimate of the stationary moments of neural responses to a fixed input (Figs. 2 and 3), a total of 20,000 independent samples (taken 200 ms apart) were drawn from the network, not including transients, as neural activity evolved according to Eq. 8. Neural activities in Fig. 2a show 1 s of simulated network activity, convolved with a 20 ms sliding window to match the effects of spike binning to compute average rates in experiments. Neural trajectories in Fig. 2b correspond to the neural activity of two cells in the network with preferred orientations 42° (u,o) and 16° (*u_j_*), over a post-transient period of 500 ms. To illustrate both the degree of modulation of the posterior covariances and the match between posterior and network covariances in Fig. 3c, the top three PCs of each posterior covariance were computed. Neural activity was then projected onto each PC, and the amount of variance along each direction was computed. The middle plots of Fig. 3c present these posterior PCs scaled by either the square root of the total variance along that direction in the GSM (in green) or in the network (in red).

Autocorrelograms in Fig. 4a were computed in 500 non-overlapping windows of 2 s of simulated neural activity each (subsampled at 0.4 ms) after stimulus onset (not including transients), and then averaged across these windows. Autocorrelograms were first computed for individual cells’ membrane potentials and then averaged across all cells. Crosscorrelograms and E–I lags in Fig. 4b–c were computed from a single 400 s-long simulation after stimulus onset and transients (without subsampling). The E–I lag for each cell was determined as the location of the maximum in the anti-symmetric component of the cross-correlogram between its total E and I input. Langevin samplers in Fig. 4a–b corresponded to neural networks with linear, time-reversible dynamics, not respecting Dale’s principle. As variability in a linear network does not depend on the input (unlike in our nonlinear circuit model), we implemented a separate Langevin sampler for each input. Autocorrelograms and crosscorrelograms for the Langevin sampler were computed as for the original network.

Average firing rates in Fig. 5a were computed from the same neural traces used in Fig. 2 to compute **u** moments (here taking the average of **r** instead of **u** in Eq. 8). To compute Fano factors in Fig. 5b we considered a doubly stochastic Gamma process and computed spike-counts over 500,000 100 ms windows. The shape parameter K_ISI_ of the Gamma process (Table S1) was chosen to reproduce the experimentally found range of Fano factors which could be less than 1 (Fig. 5b), resulting in more regular spike trains than an inhomogenous Poisson process. Power spectra in Fig. 5c were computed from the (across-cell) average neural activity (membrane potentials), following standard approaches^8^, using the same samples as the autocorrelograms of Fig. 4a (see above). Gamma peak frequency was identified as the location of the local maximum (within the gamma band, 20– 80 Hz) of the power spectrum. Transients in Fig. 5d correspond to average firing rates across E cells and trials *(n* = 100). These were also averaged over 10 ms windows to mimic the resolution of the experimental results. To account for the response delays observed in experimental data, we simulated a random delay time (truncated Gaussian, with 45 ms mean and 5 ms s.d.) to each E–I cell pair in the network. Conductance changes (relative to baseline) in Fig. 5e correspond to across trial averages (n = 20) for a single neuron with preferred orientation aligned to that of the stimulus (see Supplementary Material, Section S3.4 for further details).

To study the effect of oscillations (Fig. 6a and b) and transient overshoots (Fig. 7a and b) on sampling accuracy, we employed a family of simplified systems which were constructed as 1-dimensional Gaussian processes76 designed to match the statistics (stationary mean and variance) of single neurons in the network, but allowing to parametrically and independently vary either the degree of oscillatoriness in the system (i.e. the kernel of the Gaussian process), or the temporal profile of the mean response (Supplementary Material, Section S5).

Autocorrelograms and power spectra of Fig. 6c were computed as in Fig. 4a and Fig. 5c, but for the directions in the space of neural responses that corresponded to the first ten PCs of neural variability. To quantify the degree of oscillatori-ness along each PC, we fitted the corresponding autocorrelogram with a function that explicitly included a, the degree of oscillatoriness of responses, as one of its parameters (Section S3.5 and eq. S12).

Overshoots in Fig. 7c and d were obtained using the same stimulus that was used to train the network at 0.7 contrast, and computed as the maximal across-trial average *(n* = 100) response of each E cell (membrane potential for c, firing rate for d), minus its stationary mean response, further averaged over 1000 delay configurations in our network (as for Fig. 5d, see above). Steadystate differences denote the magnitude of mean evoked responses of each cell with respect to its mean pre-stimulus response. Fig. 7e shows novel analyses of experimental recordings from awake macaque V1 during the presentation of moving gratings of different orientations29 (data released in the repository of Ref. 77). Following the same procedure as in Ref. 8, only cells that were significantly tuned (orientation tuning index greater than 0.75) and had an average evoked rate above 1 spike per second were included in the analysis. For each cell and each stimulus, a time-dependent firing rate trace was first obtained by averaging spikes across trials in a 50 ms sliding square window. From these traces, overshoots and steady-state differences were then computed (dots in Fig. 7e top) as average evoked responses excluding transients (*t* > 160 ms after stimulus onset) minus average baseline responses (computed from the 300 ms prior to stimulus presentation). The overshoot size is computed as the maximum of the response trace for *t* < 160 ms after stimulus onset, minus the same average evoked response previously computed. Linear regression was performed using SciPy’s ‘linregress’ function, which reports a two-sided p-value (null hypothesis: zero slope), using a Wald test with a t-distribution of the test statistic. Results in the bottom plots of Fig. 7d and e were computed by averaging stimuli presented at each neuron’s preferred orientation (±30°) or orthogonal to its preferred orientation (±30°). We tested for significance in overshoot tuning using SciPy’s ‘ttest_ind’ function (null hypothesis: identical average).

