## Supplementary Material for "Cortical-like dynamics in recurrent circuits optimized for sampling-based probabilistic inference"

**S1 Mapping from GSM variables to neural responses** [Figs. S1 and S2] . . . . . . . . . . . . . . . **2**

**S2 Training the neural network** . . . . . . . . . . . . . . . . . . . . . . . . . . . . . . . . . **2**

S2.1 Training set and cost function 2

S2.2 Computing the moments of neural responses 4

S2.3 Network optimization 5

**S3 Analyses of the optimized network** . . . . . . . . . . . . . . . . . . . . . . . . . . . . . **5**

S3.1 Parameters [Fig. S3] 5

S3.2 Divisive normalization [Fig. S4] 6

S3.3 Network mechanism underlying oscillations [Fig. S5] 7

S3.4 Computing Fano factors and conductances 7

S3.5 Quantifying oscillatoriness 8

**S4 Analyses of other networks** . . . . . . . . . . . . . . . . . . . . . . . . . . . . . . . . . **8**

S4.1 Random networks [Figs. S6 and S7] 8

S4.2 Other optimized networks 10

S4.3 Control network optimized without enforcing Dale's principle [Fig. S8] 10

S4.4 Control networks optimized for other objectives [Figs. S9–S13] 12

**S5 Mathematical analyses of oscillations and transients** . . . . . . . . . . . . . . . . . . . . . **15**

S5.1 Sampling accuracy of statistically stationary dynamics 16

S5.2 Oscillations [Fig. S14] 17

S5.3 Transients [Fig. S15] 21

**S6 Online repositories** [Table S1] . . . . . . . . . . . . . . . . . . . . . . . . . . . . . . . **24**

### S1 Mapping from GSM variables to neural responses

Our choice to have membrane potential values rather than firing rates in the network represent the latent variables of the GSM might seem unusual at first, as most computational theories assign meaningful representations to firing rates, not membrane potentials (Dayan and Abbott, 2001; but see e.g. Ujfalussy et al., 2015). However, note that in our case, non-negative firing rates were inappropriate for representing the original GSM variables which could be negative as well as positive. This mismatch could be addressed in either of two mathematically equivalent ways. Membrane potential variables (which can be positive or negative, i.e. above or below the resting value) could represent the GSM variables and firing rates could be computed "phenomenologically" as some transformation of membrane potentials (Eq. 9). Alternatively, we could have defined a transformed generative model in terms of variables that are directly represented by firing rates, with these variables then being appropriately (inversely) transformed into the original GSM variables. Importantly, as we have shown in Orbán et al. (2016), these two solutions are mathematically equivalent in that they imply the same predictive distribution over stimuli $\mathcal{P}(\mathbf{x})$ (which determines the model's statistical validity as a model of natural image patches) and ultimately the same target posteriors $\mathcal{P}\left(\mathbf{r}|\mathbf{x}\right)$ over the rates $\mathbf{r}$ for any stimulus $\mathbf{x}$ (which determine the model's predictions for firing rates). We chose the first solution for two reasons. First, following Hennequin et al. (2018), the form of network dynamics we used was primarily defined in terms of membrane potentials (Eq. 8), with rates derived "phenomenologically" (Eq. 9). Second, the GSM posteriors were approximately Gaussian, and as such well characterised by their first two moments. This made them suitable targets for the second-order moment-matching objective we used for training our network (Eqs. 17-20). In contrast, the second solution would have required matching highly non-Gaussian target distributions (Hennequin and Lengyel, 2016).

We also introduced a further transformation, such that even membrane potential values, $\mathbf{u}$, represented a (mildly) transformed version of the GSM latent variables. This was because under the GSM model we used (Eqs. 1-6, Fig. S1), the posterior variances over latent variables $\mathbf{y}$ only depended on contrast, but not on anything else in the stimulus. In particular, they did not depend on the orientation content of the stimulus. In contrast, empirical observations from V1 recordings indicate that Fano factors, related to $\mathbf{r}$ and thus ultimately $\mathbf{u}$ in our neural network model (Eqs. 8 and 9), do depend on stimulus orientation (Hennequin et al., 2018). Just as above, this mismatch could have also been addressed in either of two mathematically equivalent ways: via a non-linear transformation of the latent variables $\mathbf{y}$ once they have been inferred, or by modifying the generative model such that it was already defined in terms of these transformed variables $\mathbf{u}$. Once again, these two models are mathematically equivalent in terms of their predictive distribution over image patches and target posterior distributions over membrane potentials, and thus our choice was dictated by practical considerations. For mathematical convenience, we chose the former formalization as there is no *a priori* reason to believe that the brain should employ a linear encoding of the GSM variables. Thus, as described in the Online Methods, the original GSM latent variables $y_i$ were transformed into membrane potentials according to Eq. 7, with parameters $\alpha_{\mathrm{nl}}$, $\beta_{\mathrm{nl}}$, and $\gamma_{\mathrm{nl}}$ respectively controlling the scaling, baseline, and power of the transformation. To obtain the kind of variance modulation observed in experiments (variance smallest for neurons driven by their preferred inputs), we set $\gamma_{\mathrm{nl}} < 1$ (Table S1). This also yielded physiologically realistic ranges for the corresponding rates (Fig. 5a). We also note that with the chosen parameters, all the mean values of the original $\mathbf{y}$ distributions lay well above threshold (Fig. S2a), implying only a mild transformation of variables (compare Fig. S2b and c).

### S2 Training the neural network

The basic approach for training our network is explained in the Online Methods. Here we provide some further details.

#### S2.1 Training set and cost function

We trained the network to perform sampling-based probabilistic inference under the GSM. To that end, we first created a small training set comprising five image patches (Fig. 2b):

$$\mathbf{x}^{\alpha} = z_{\alpha}\,\mathbf{A}\,\bar{\mathbf{y}} \tag{S1}$$

with $\alpha = 1, \ldots, 5$ and $z_{\alpha} \in \{0, 0.125, 0.25, 0.5, 1.0\}$. Therefore, these stimuli had the same content $\bar{\mathbf{y}}$: a 27°-wide Gaussian function centered around 0° (i.e. a single dominant orientation), and differed only in their contrast

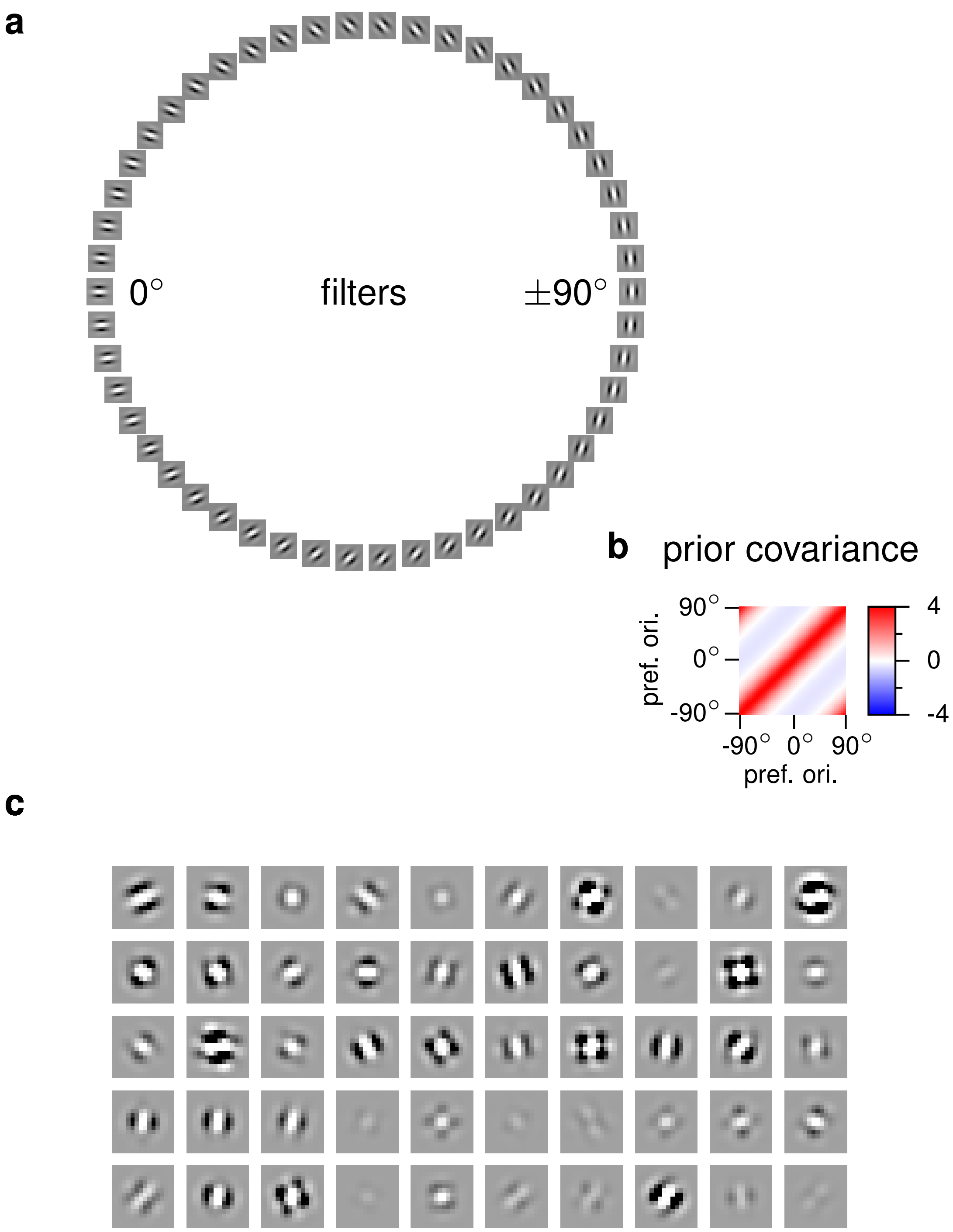


**Fig. S1 | GSM parameters and stimulus generation. a,** Filters in the GSM. Each filter image is a column of **A** (Eq. 1), also used as feedforward receptive fields in the network: $\mathbf{W}^{\text{ff}} = [\mathbf{A}\,\mathbf{A}]^{\text{T}}/15$ (Eq. 14; cf. Fig. 1a and c). **b,** Prior covariance in the GSM (**C**, Eq. 2). **c,** Sample stimuli generated by the GSM, also used for testing the network's generalization (Fig. 3b-c).

(Fig. 2b). As the parametrization of our network was rotationally invariant, such a stimulus was in fact representative of all image patches that could be obtained by rotating this patch around the center. For each training stimulus $\mathbf{x}^{\alpha}$, we computed the corresponding posterior distributions over $\mathbf{u}$ under the GSM (Eqs. 6 and 7). We called these distributions the "target distributions", and their corresponding means $\boldsymbol{\mu}^{\alpha}_{\text{tgt}}$ and covariances $\boldsymbol{\Sigma}^{\alpha}_{\text{tgt}}$, the "target moments" (Eqs. 15 and 16, Fig. 2c,d).

We minimized a cost function involving the distance between the statistical moments of the neural responses sampled over time during the presentation of each stimulus, and the target moments corresponding to the same

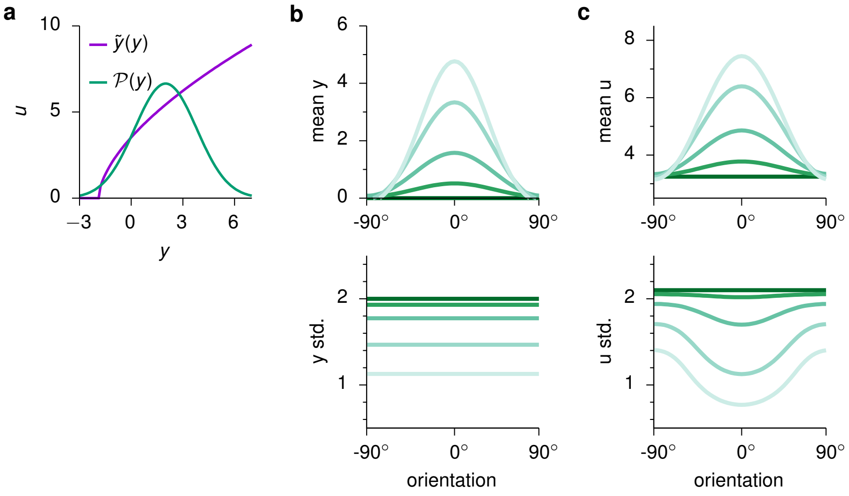


**Fig. S2 | Nonlinear transformation of the GSM variables. a,** Transformed variable $u$ as a function of the latent variable $y$ of the GSM (purple; Eq. 7), and a representative posterior distribution over $y$ (green). **b-c,** First two moments (top: mean; bottom: standard deviation) of the distributions over the original GSM variables $\mathbf{u}$ (**b**) and the transformed variables $\mathbf{y}$ (**c**). Colors indicate contrast, as in Fig. 2c.

stimulus. Each stimulus was presented $N_{\text{trials}}$ times independently, in which it was held constant over a period of 500 ms. Just before stimulus onset, neural responses were initialized by drawing from a Gaussian distribution $\mathcal{N}(\boldsymbol{\mu}_0, \boldsymbol{\Sigma}_0)$ (Table S1). For every time $t$ relative to stimulus onset, we denote the across-trial moments of neural responses by $\boldsymbol{\mu}^\alpha(t)$, $\boldsymbol{\Sigma}^\alpha(t)$ (see Section S2.2). For convenience, we also define $\boldsymbol{\sigma}^\alpha(t) = \text{diag}\big(\boldsymbol{\Sigma}^\alpha(t)\big)$.

The cost function took the form shown in the Online Methods (Eqs. 17-21) with parameters $\epsilon_{\text{mean}} \propto 1.0/(N_{\text{E}}\, T_{\text{max}})$, $\epsilon_{\text{var}} \propto 2.0/(N_{\text{E}}\, T_{\text{max}})$, and $\epsilon_{\text{cov}} \propto 1.0/(N_{\text{E}}^2\, T_{\text{max}})$ respectively controlling the relative importance of matching the means, variances, and covariances of the target distributions (see Table S1 for actual values).

### S2.2 Computing the moments of neural responses

The cost function defined by Eq. 17 required us to evaluate how the probability distribution over neural responses, $\mathbf{u}$, evolved over time for a given input and a given set of network parameters. In particular, we needed to track the time evolution of the mean and covariance of network activity (Section S2.1):

$$\boldsymbol{\mu}(t) = \langle \mathbf{u}(t) \rangle \tag{S2}$$

$$\boldsymbol{\Sigma}(t, s) = \left\langle \big(\mathbf{u}(t) - \boldsymbol{\mu}(t)\big)\, \big(\mathbf{u}(t+\Delta t) - \boldsymbol{\mu}(t+\Delta t)\big)^{\mathsf{T}} \right\rangle \tag{S3}$$

starting from an initial distribution with moments $\boldsymbol{\mu}_0$ and $\boldsymbol{\Sigma}_0$. Here, $\langle\cdot\rangle$ denotes trial-averaging. To do this, we employed two different methods. The first approach, which we refer to as the "stochastic method", consisted of approximating the averages via sampling, i.e. simulating stochastic network dynamics in a set of trials using the same stimulus (Eqs. 8, 9 and 14) and computing the across-trial sample mean and sample covariance at each time step.

The second approach, which we refer to as "assumed density filtering" (ADF), used deterministic equations of motion for computing the across-trial moments. First, based on Hennequin and Lengyel (2016), the following exact differential equations were used to describe the evolution of $\boldsymbol{\mu}(t)$ and $\boldsymbol{\Sigma}(t, \cdot)$:

$$\frac{\mathrm{d}\boldsymbol{\mu}(t)}{\mathrm{d}t} = \mathbf{T}^{-1}\, \big[-\boldsymbol{\mu}(t) + \mathbf{h}(t) + \mathbf{W}\, \boldsymbol{\nu}(t)\big] \tag{S4}$$

$$\frac{\mathrm{d}\boldsymbol{\Sigma}(t,0)}{\mathrm{d}t} = \big[\mathbf{T}^{-1}\, \boldsymbol{\Sigma}^*(t)\big] + \big[\mathbf{T}^{-1}\, \boldsymbol{\Sigma}^*(t)\big]^{\mathsf{T}} + \mathcal{J}(t)\; \boldsymbol{\Sigma}(t,0) + \boldsymbol{\Sigma}(t,0)\; \mathcal{J}(t)^{\mathsf{T}} \tag{S5}$$

$$\frac{\mathrm{d}\boldsymbol{\Sigma}^*(t)}{\mathrm{d}t} = -\frac{1}{\tau_\eta}\, \boldsymbol{\Sigma}^*(t) + \boldsymbol{\Sigma}^\eta\, \mathbf{T}^{-1} + \boldsymbol{\Sigma}^*(t)\; \mathcal{J}(t)^{\mathsf{T}} \tag{S6}$$

$$\frac{\mathrm{d}\boldsymbol{\Sigma}(t,\Delta t)}{\mathrm{d}\Delta t} = e^{-\Delta t/\tau_\eta}\, \big[\mathbf{T}^{-1}\, \boldsymbol{\Sigma}^*(t)\big]^{\mathsf{T}} + \boldsymbol{\Sigma}(t,\Delta t)\; \mathcal{J}(t+\Delta t)^{\mathsf{T}} \qquad \forall \Delta t > 0 \tag{S7}$$

where $\mathbf{T}$ is the diagonal matrix of membrane time constants (Table S1), $\boldsymbol{\nu} = \langle \mathbf{r} \rangle$ is the average firing rate of neurons, $\boldsymbol{\Sigma}(t, -\Delta t) = \boldsymbol{\Sigma}^{\mathsf{T}}(t, \Delta t)$ is the time-lagged cross covariance of membrane potentials in the network, $\boldsymbol{\Sigma}^* = \left\langle \boldsymbol{\eta}\,(\mathbf{u} - \boldsymbol{\mu})^{\mathsf{T}} \right\rangle$ is the instantaneous cross-covariance between membrane potentials and temporally correlated process noise $\boldsymbol{\eta}$ (Eq. 8) with instantaneous covariance $\boldsymbol{\Sigma}^{\eta} = \langle \boldsymbol{\eta}\,\boldsymbol{\eta}^{\mathsf{T}} \rangle$, and

$$\mathcal{J} = \mathbf{T}^{-1}\left[-\mathbf{I} + \mathbf{W}\,\mathrm{diag}\left(\frac{\partial \boldsymbol{\nu}}{\partial \boldsymbol{\mu}}\right)\right] \tag{S8}$$

is the Jacobian of Eq. S4 w.r.t. $\boldsymbol{\mu}$. Integrating Eqs. S4-S7 in turn required evaluating some nonlinear moments of $\mathbf{u}$, namely covariances between membrane potentials ($\mathbf{u}$) and firing rates ($\mathbf{r}$). For the SSN, these moments can be obtained in closed form, assuming the full joint (space-time) distribution of membrane potentials is Gaussian (Hennequin and Lengyel, 2016). Thus, in contrast to the first (stochatic) method which leads to unbiased, but potentially high-variance, estimation of the moments, the ADF method leads to zero-variance, but potentially biased estimates. To train the network, as described below, we combined the strengths of these two approaches.

#### S2.3 Network optimization

Training the network involved optimizing the recurrent connectivity, the stimulus-independent process noise covariance, and a small set of parameters describing the single, static nonlinear transformation of the feedforward receptive fields (see Online Methods). Given the initial conditions (and a fixed process noise history for the stochastic method), the equations become fully differentiable, therefore we used gradient-based methods for optimization. The optimization procedure was implemented in OCaml and automatic differentiation was employed to compute the gradient of the cost function defined in Eq. 17 by back-propagation through time (Werbos, 1990).

We trained the network in two stages. During the first stage, we employed the stochastic method using 50 trials for each training stimulus to compute the corresponding moments of network responses, and performed 250 iterations of the ADAM optimizer (Kingma and Ba, 2014). Both the network's initial conditions and the process noise were re-sampled for each trial and iteration. In this case $T_{\mathrm{min}}$ in Eqs. 18-20 was set to 0 ms for the first iteration, and gradually and linearly increased in each iteration towards $T_{\mathrm{max}}$, up to a maximum of $T_{\mathrm{max}} - 50$ ms. We found this annealing procedure helped keep the network stable and encouraged the discovery of fast convergence to the target distribution (i.e. short burn-in) during the initial phase of training, while progressively focusing on the stationary moments of the output distribution (after the initial burn-in period).

In the second stage, we refined the optimization using the ADF method together with an L-BFGS-B optimizer (Zhu et al., 1997), leaving the cost-integration time window at its minimum (50 ms, as reached by the end of the first phase). The slowness penalty cost in Eq. 21 was only applied during ADF-based optimization and, for simplicity, it was approximated using stationary lagged correlations predicted by the ADF-method (Eq. S7) in the limit of temporally white process noise (Hennequin and Lengyel, 2016).

### S3 Analyses of the optimized network

In the main text, we comprehensively analyze the behavior of the optimized network (Figs. 2–7). Here we present some further properties of it and the details of the analyses. See also Sections S5.2.1 and S5.3.1 for further details of the analyses of oscillations and transients.

#### S3.1 Parameters

After training, the connectivity profile of either excitatory or inhibitory cells in the optimized network was largely independent of whether the postsynaptic cell was excitatory or inhibitory (Fig. S3a, top). Overall, recurrent excitatory and inhibitory connections had similar tuning widths, with excitatory connections being slightly more broadly tuned than inhibitory ones (Fig. S3a, middle). Nevertheless, the net excitatory input to any one cell in the network was still more narrowly tuned than the net inhibitory input, due to the responses of presynaptic excitatory cells being more narrowly tuned than those of inhibitory ones (not shown).

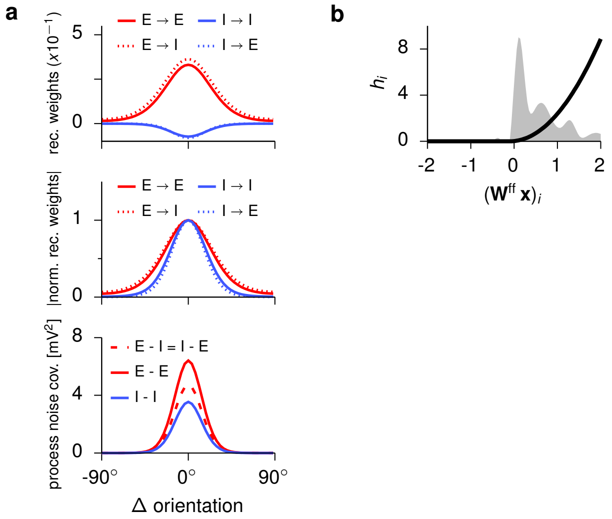


**Fig. S3 | Parameters of the optimized network. a,** Recurrent weights (top: raw weights; middle: normalized absolute values) and process noise covariance (bottom) after training. Weights and covariances are shown as one row (of each quadrant) of $\mathbf{W}$ (Eq. 8) and $\boldsymbol{\Sigma}^{\eta}$ (Eq. 11), respectively, as they are circulant. Thus, each line shows the weights connecting, or the covariance between, cells of different types (see legend) as a function of the difference in their preferred stimuli. **b,** Input nonlinearity (Eq. 14), converting feedforward receptive field activations $(\mathbf{W}^{\text{ff}}\,\mathbf{x})_i$ into network inputs $h_i$ (black). For comparison, the distribution of inputs across all cells for the training set is presented in gray.

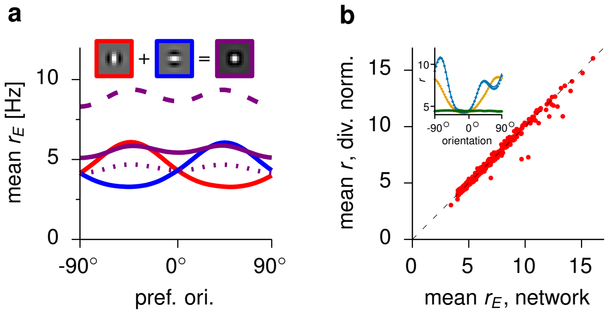


**Fig. S4 | Divisive normalisation in the optimized network. a,** The optimized network's response (solid purple) to a sum of two stimuli (insets), compared to its responses to individual stimuli presented separately (solid red/blue), their average (dotted purple), and sum (dashed purple). **b,** Scatter plot of neural responses vs. fit to a phenomenological divisive normalisation model (Eq. S9). Inset shows three representative average response profiles across the network (dots) and the phenomenological model's fit (lines). Note the close match between the network and the phenomenological model in all three cases.

The optimized network also retained a substantial amount of process noise that was larger in excitatory than in inhibitory neurons, and highly correlated both between the E and I cell of a pair and between cells with different tuning (up to a $\sim 30°$ tuning difference; Fig. S3a, bottom). Such correlated noise is detrimental for a precise encoding of the input (Abbott and Dayan, 1999; Averbeck et al., 2006) but it creates the ideal substrate for the network to modulate its variability in a stimulus dependent manner (Hennequin et al., 2018).

The optimized input transformation, capturing the nonlinear effects of upstream preprocessing of visual stimuli, had a threshold that was just below the distribution of receptive field outputs, ensuring that all stimulus-related information was transmitted in the input signal, and an exponent close to two, remarkably similar to that used by the cells of the network (Fig. S3b; cf. Eq. 9 and table S1).

### S3.2 Divisive normalization

Divisive normalization, or sublinear summation of neural responses, has been proposed as a canonical computation in cortical circuits (Carandini and Heeger, 2012). Under divisive normalization, the network's response

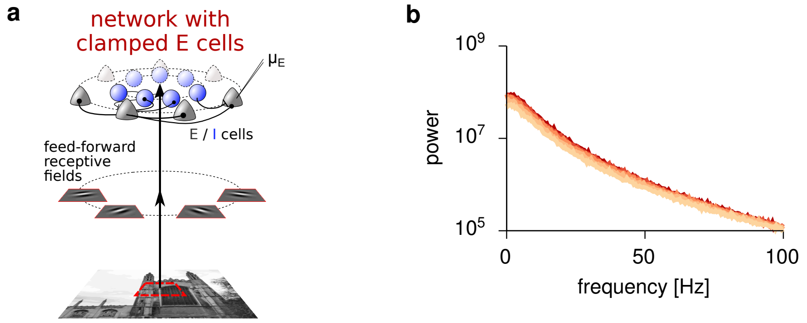


**Fig. S5 | Blocking oscillations in the optimized network by voltage-clamping E cells. a,** Illustration of the clamping experiment in the optimized network (cf. Fig. 1b). For each stimulus, each E cell's voltage was clamped to its mean voltage ($\mu_E$) obtained when the network was presented with the same stimulus without voltage clamp. **b,** LFP power spectra in the network after voltage-clamping of the E population at different contrast levels (colors as in Fig. 5c). Note the absence of gamma peak (cf. Fig. 5c, inset).

to the sum of two stimuli is smaller than the sum of its responses to the individual stimuli (it usually lies between the average and the sum). We found this phenomenon to occur in our network, too, when combining training stimuli (Fig. S4a). To verify that this was also the case for arbitrary stimuli, we also fitted a standard phenomenological model of divisive normalization (adapted from Carandini and Heeger, 2012) to the behavior of our trained network. Specifically, for each image patch in a set of 500 random stimuli (same procedure as used to generate the generalization dataset, Online Methods), the model attempted to capture the mean firing-rate responses $\langle \mathbf{r} \rangle$ of our trained network as a function of the feedforward input $\mathbf{h}$ alone (i.e. without regard to its recurrent dynamics) in the following functional form:

$$\langle r \rangle_i^{(\beta)} = b_2 + \frac{h_i^{(\beta)} + b_1}{\left(\mathbf{M}\,\mathbf{h}^{(\beta)}\right)_i + s^2} \tag{S9}$$

where $h_i^{(\beta)}$ and $\langle r \rangle_i^{(\beta)}$ were the feedforward input and the average firing rate of cell $i$ in response to stimulus $\beta$, respectively, and $b_1$, $b_2$ and $s$ were constant parameters. The parameter matrix $\mathbf{M}$ is responsible for normalization, by dividing the input $h_i$ by a mixture of competing inputs to other neurons. $\mathbf{M}$ was parameterized as a symmetric circulant matrix to respect the rotational symmetry of the trained network. In total, our model of divisive normalization had $3 + \left(N_E/2\right) + 1 = 29$ free parameters. Model fitting was performed via minimization of the average squared difference between network and model rates, plus an elastic energy regularizer for neighbouring elements of $\mathbf{M}$. We used the L-BFGS-B algorithm and Tensorflow for optimization.

We found that the divisive normalization model accurately captured the responses of our trained network, not only to the stimuli in the set used to fit the divisive normalization model, but also to a second set of 500 novel stimuli (Fig. S4b). The divisive normalization model also outperformed both a linear model and a model of subtractive inhibition (not shown). These results show comprehensively that, in line with empirical data, our trained network performed divisive normalization of its inputs under general conditions.

### S3.3 Network mechanism underlying oscillations

To determine whether oscillations in our network resulted from the interaction of E and I cells, or whether they arose within either of these populations alone, we conducted two simulated experiments. In each simulation, either of the two populations (E or I) was voltage-clamped to its (temporal) mean, as calculated from the original network, for each input in the training set. Thus, recurrent input from the clamped population was effectively held constant to its normal mean, but did not react to changes in the other population.

As expected for a network in the inhibition-stabilized regime, clamping of the I cells resulted in unstable runaway dynamics, precluding further analysis of oscillations. In contrast, the network with clamped E voltages (Fig. S5a) remained stable, but the peak in its LFP power spectrum characteristic of gamma oscillations was no longer present (Fig. S5b). This shows that gamma oscillations in the original network required interactions between E and I cells (i.e. they were generated by the so-called "PING" mechanism; Tiesinga and Sejnowski 2009).

#### S3.4 Computing Fano factors and conductances

Some of the comparisons we performed between our model and neurophysiological results involved quantities that were not explicitly modelled by our rate-based network. These quantities included spike count Fano factors and input conductances. Here, we describe how we estimated these quantities in our model, based on the membrane potentials and firing rates that were explicitly represented in the model equations (Eqs. 8 and 9).

Following Hennequin et al. (2018), spike count statistics were estimated assuming doubly-stochastic action potential generation. Specifically, spikes were generated by an inhomogeneous Gamma process with the time-varying rate given by $r_i(t)$ for each neuron $i$ and the shape of the interspike interval distribution controlled by an additional parameter, $K_{\mathrm{ISI}}$ (Table S1). Our results were qualitatively robust to the choice of this parameter, which primarily determined the overall magnitude of Fano factors – in particular $K_{\mathrm{ISI}} > 1$ was needed to achieve Fano factors $< 1$ at high contrast – but not their modulation by stimuli.

To estimate input conductances, we equated the total (excitatory or inhibitory) input current in our model to the (excitatory or inhibitory) current in a canonical conductance-based model (Dayan and Abbott, 2001):

$$\frac{1}{\tau_i}\left(\sum_{j\in\mathrm{E/I}} \mathbf{W}_{ij}\,\mathbf{r}_j(t)\right) \approx \frac{1}{C_\mathrm{m}}\,\mathbf{g}_i^{\mathrm{E/I}}(t)\,\left[V^{\mathrm{E/I}} - \left(\mathbf{u}_i(t) + V_\mathrm{rest}\right)\right] \tag{S10}$$

where $C_\mathrm{m}$ is the membrane capacitance, $V^{\mathrm{E/I}}$ denote the reversal potentials for E/I currents, and $V_\mathrm{rest}$ is a baseline (resting) potential added to the membrane potentials of our model. Solving for $\mathbf{g}^{\mathrm{E/I}}$, we obtained:

$$\mathbf{g}_i^{\mathrm{E/I}}(t) \approx \frac{C_\mathrm{m}\sum_{j\in\mathrm{E/I}} \mathbf{W}_{ij}\,\mathbf{r}_j(t)}{\tau_i\left[V^{\mathrm{E/I}} - \left(\mathbf{u}_i(t) + V_\mathrm{rest}\right)\right]} \tag{S11}$$

We chose $C_\mathrm{m} = 20.0$ pF, $V^\mathrm{E} = 0$ mV, $V^\mathrm{I} = -80$ mV, and $V_\mathrm{rest} = -65$ mV. Conductances in Fig. 5e are shown relative to their steady-state values during spontaneous activity.

#### S3.5 Quantifying oscillatoriness

To quantify oscillatoriness in neural responses along some direction in state space, we computed the corresponding projection of neural responses, and then fitted the following parametric function to its autocorrelogram:

$$C_\alpha(\Delta t) = \left[(1-\alpha) + \alpha\,\cos\!\left(2\pi f\,\Delta t\right)\right]\left[\frac{\tau}{\tau-\tau_\eta}e^{-\Delta t/\tau} - \frac{\tau_\eta}{\tau-\tau_\eta}e^{-\Delta t/\tau_\eta}\right] \tag{S12}$$

where $\tau$ and $\tau_\eta$ are two time constant parameters, $\alpha \in [0,1]$ quantifies the degree of oscillatoriness, and $f$ represents the dominant oscillation frequency. In Fig. 6d and Fig. S7b, the fit was performed using Tensorflow, for each principal component and each input. The form of Eq. S12 was motivated by noting that, for $\alpha = 0$, it reduces to

$$C_0(\Delta t) = \frac{\tau}{\tau-\tau_\eta}e^{-\Delta t/\tau} - \frac{\tau_\eta}{\tau-\tau_\eta}e^{-\Delta t/\tau_\eta} \tag{S13}$$

which is the autocorrelation function of the fluctuations in a single (isolated, and thus not oscillating) neuron with membrane time constant $\tau$ receiving noisy inputs with correlation time $\tau_\eta$ (this can be seen by integrating Eqs. S5-S7). More generally, for $\alpha > 0$, $C_0(\Delta t)$ (Eq. S13) determines the envelope of $C_\alpha(\Delta t)$ (Eq. S12):

$$(1-2\,\alpha)\;C_0(\Delta t) \le C_\alpha(\Delta t) \le C_0(\Delta t) \tag{S14}$$

### S4 Analyses of other networks

Most of the analyses we present in the main text (all simulation results except those in Fig. 5, right column) were based on a single optimised network. Here we present further analyses of a number of other networks to demonstrate the non-triviality, robustness and specificity of our results, and to dissect which components of our optimization approach were responsible for our main results.

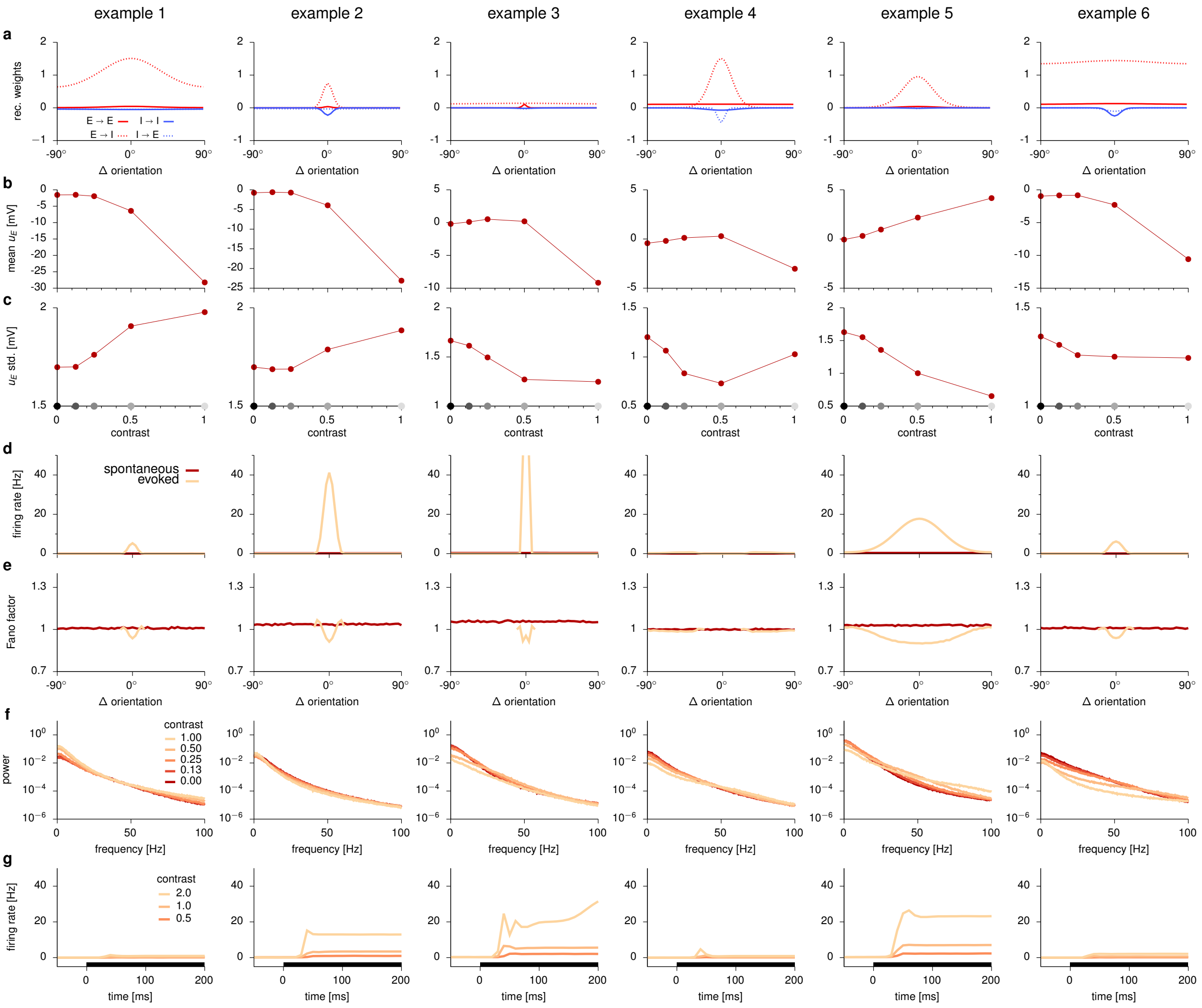


**Fig. S6 | Random networks.** Results for six example random networks are shown (columns, see details in text). **a,** Recurrent weights as a function of the difference in the preferred stimuli of two cells (cf. Fig. S3a, top). Different lines are for weights connecting cells of different types (legend). **b-c,** Mean (**b**) and standard deviation of membrane potential responses (**c**), averaged over the population, as a function of contrast (cf. Fig. 3a). Gray dots on x-axis indicate training contrast levels. **d-e,** Mean firing rate (**d**) and Fano factor (**e**) of neurons as a function of stimulus orientation (relative to their preferred orientation) during spontaneous (dark red) and evoked activity (light orange; cf. Fig. 5a-b). The peak mean rate of example 3 exceeded 50 Hz and is thus shown as clipped in this figure. Note that mean rate tuning curves (during evoked activity) were very narrow for most networks (all but example 5), resulting in 0 Hz rates and thus undefined Fano factors for stimuli further away from the preferred orientation. **f,** LFP power spectra at different contrast levels (colors as in Fig. 5c). **g,** Average rate response around stimulus onset at different contrast levels (colors as in Fig. 5d). Black bars show stimulus period. Note the absence of gamma peaks in **f** (cf. Fig. 5c, inset) and of transients in **g** (cf. Fig. 5d).

### S4.1 Random networks

The parametrization of our network was highly constrained (e.g. ring topology with circulant and symmetric weight and process noise covariance structure, 1:1 E:I ratio, fixed receptive fields). Therefore, we first wanted to know whether these constraints alone, without any optimization, were sufficient to generate the results we obtained in the optimized network. For this, we sampled weight matrices at random, by drawing each of the 8 hyperparameters of the weight matrix (Eq. 10) from an exponential distribution truncated between 0.1 and 10 times the values originally found by optimization. We discarded matrices that were either unstable or converged to a trivial solution (all mean rates equal 0). Less than 20% of the generated matrices satisfied these criteria, further confirming that optimization was actually non-trivial. Fig. S6 shows the first 6 cases that satisfied these criteria. These networks displayed a wide array of behaviors. For example, the standard deviation of responses could go up, down, or even be non-monotonic with contrast, while the range of mean rates also varied wildly.

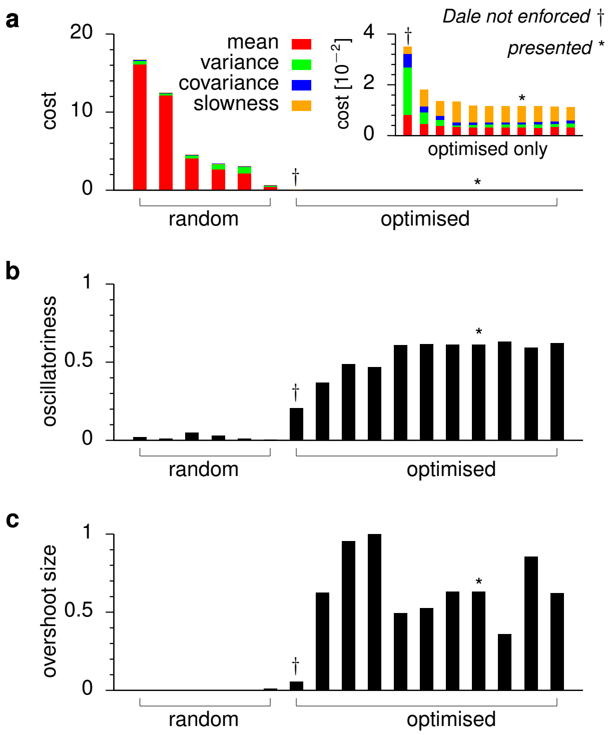


**Fig. S7 | Comparison of random and optimized networks in terms of cost and dynamical features.** Networks are ranked in all panels in order of decreasing total cost achieved by them (shown in **a**). **a,** Total cost (Eq. 17) computed for each of the random networks (Section S4.1) and for networks that were optimized for this cost (Sections S4.2 and S4.3). Each component of the cost is presented in a different color (legend, see Eqs. 18-21 for mathematical definitions). The inset shows the optimized networks only (note different y-scale). The network presented in the main text is indicated with $\star$, and the network optimized without enforcing Dale's principle is marked with †. **b-c,** Oscillatoriness (**b**) and transient overshoot size (**c**) for each network in **a**. Oscillatoriness was computed by numerical fits of Eq. S12 to the autocorrelogram of the "LFP" generated by the network (Online Methods, Section S3.5). Transients in population-average firing rates were quantified as the size of the overshoot normalized by the change in the steady state mean (see also Fig. 7a).

Correspondingly, the cost achieved by these random networks was 1-3 orders of magnitude higher than that achieved by the optimized network (Fig. S7a). As simple summary statistics of the overall dynamics of these networks, we also quantified their degree of oscillatoriness and the size of transient overshoots in them (Fig. S7b-c). Importantly, compared to optimized networks (see below) – including the one presented in the main text –, oscillations and transients were almost entirely absent from these networks. Thus, random networks behaved fundamentally differently than the optimized network.

### S4.2 Other optimized networks

Having established that random networks did not show the same behavior as our optimized network, we also sought to confirm the converse: that well-optimized networks reliably showed similar behavior. This was important because our cost function was highly non-convex. Therefore, any minimum our optimizer found, such as that corresponding to the network presented in the main text, had no guarantee of being the global minimum. In order to check that the results we obtained for the network presented in the main text were representative of the best achievable minima of the cost function, we trained further 10 networks starting from random initial conditions. None of these networks achieved substantially lower costs than the one we presented in the main text, and those whose final cost was at least approximately as low as that of the original network (9 out of 10; Fig. S7a) all had substantial oscillatoriness and transient overshoots (Fig. S7b-c).

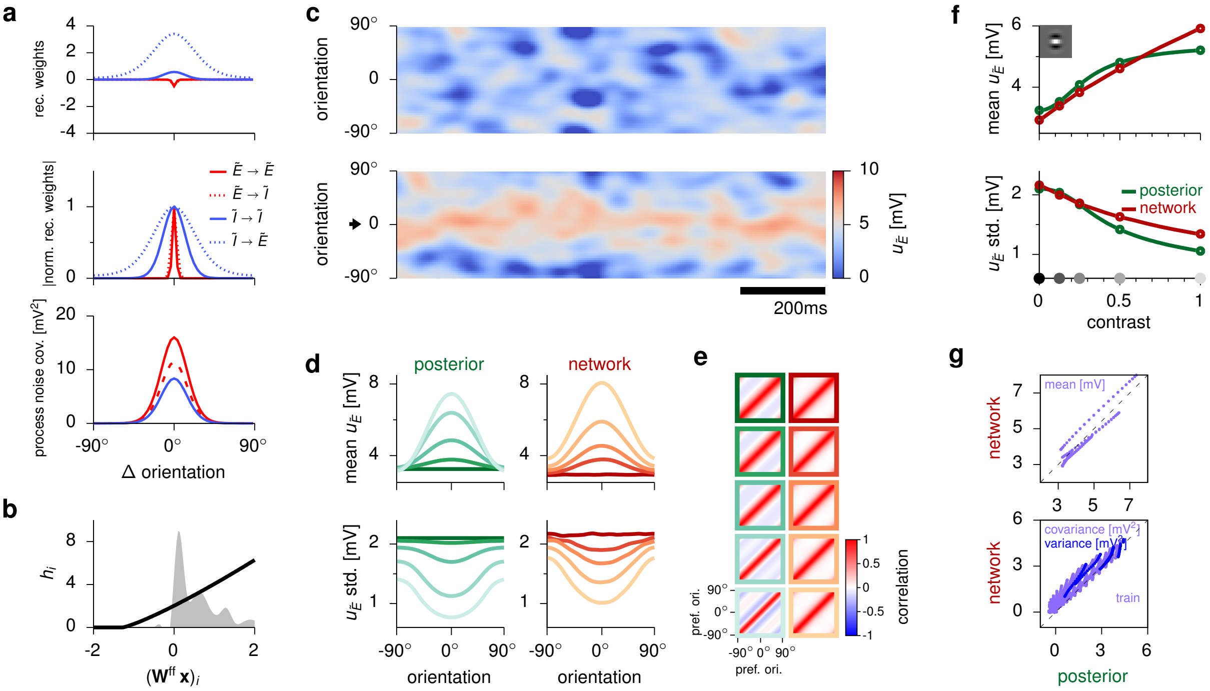


**Fig. S8 | Control network: Dale's principle not enforced.** **a,** Recurrent weights (top: raw weights, Eq. 8; middle: normalized absolute values) and process noise covariance (bottom, Eq. 11) between cells of different (here: notional) types (see legend) as a function of the difference in their preferred stimuli (cf. Fig. S3a, Fig. S6a). Note that the outgoing weights of the cells whose moments were constrained ($\tilde{E}$ cells) were actually negative. Therefore, in effect, these cells became inhibitory during training. **b,** Input nonlinearity (Eq. 14), converting feedforward receptive field activations $(\mathbf{W}^{\mathrm{ff}}\,\mathbf{x})_i$ into network inputs $h_i$ (black). For comparison, the distribution of inputs across all cells for the training set is presented in gray (cf. Fig. S3b). **c,** Sample population activity of $\tilde{E}$ cell membrane potentials $u_{\tilde{E}}$ (y-axis, ordered by their preferred orientation) at zero (top) and high (bottom) contrast (cf. Fig. 2a). The high contrast stimulus has a dominant orientation at 0° (arrow). **d,** Mean (top) and standard deviation (bottom) of latent variables under the ideal observer's posterior distribution (left, green) and of $\tilde{E}$ cell membrane potentials $u_{\tilde{E}}$ under the network's stationary distribution (right, red), ordered by their preferred orientation, for each stimulus in the training set (cf. Fig. 2c). **e,** Correlation matrices of the ideal observer's posterior distributions (left, green) and the network's stationary response distributions (right, red), for each stimulus in the training set (cf. Fig. 2d). Line colors in **d** and frame colors in **e** correspond to training stimuli with different contrast levels as in Fig. 2b-d. **f,** Mean and standard deviation of latent variables (green) and stationary network responses (red) averaged over the population, as a function of the contrast of stimuli in the training set (see inset in top panel for a representative stimulus; cf. Fig. 3a, Fig. S6b-c). Gray dots on x-axis indicate training contrast levels. **g,** Stationary means (top), variances (bottom, blue) and covariances (bottom, purple) during network activity (y-axis) versus under the posterior (x-axis). Each dot corresponds to the response of an individual cell (top, bottom blue) or cell-pair (bottom, purple) to one of the training stimuli (cf. Fig. 3b).

### S4.3 Control network optimized without enforcing Dale's principle

In order to see how much the biological constraints we used for the optimized network, and in particular enforcing Dale's principle, were *necessary* to achieve the performance and dynamical behavior of the original network, we optimized a network with the same cost function as for the original network (Eq. 17) but without enforcing Dale's principle. This meant that the signs of synaptic weights in each quadrant of the weight matrix (the $a_{\mathrm{XY}}$ coefficients in Eq. 10) were not constrained. Otherwise, optimization proceeded in the same way as before (Section S2). The training of this network proved to be much more difficult and prone to result in unstable networks, which we avoided by early stopping. Interestingly, the network optimized this way still obeyed Dale's principle (Fig. S8a, top[1]). Overall, the *stationary* behavior of this network was broadly similar to that of the originally

[1]Note that the notional labels $\tilde{E}$ and $\tilde{I}$ in Fig. S8a were swapped relative to what one would expect from the signs of the corresponding weights, but it was still the case that all outgoing synapses of any one cell had the same sign.

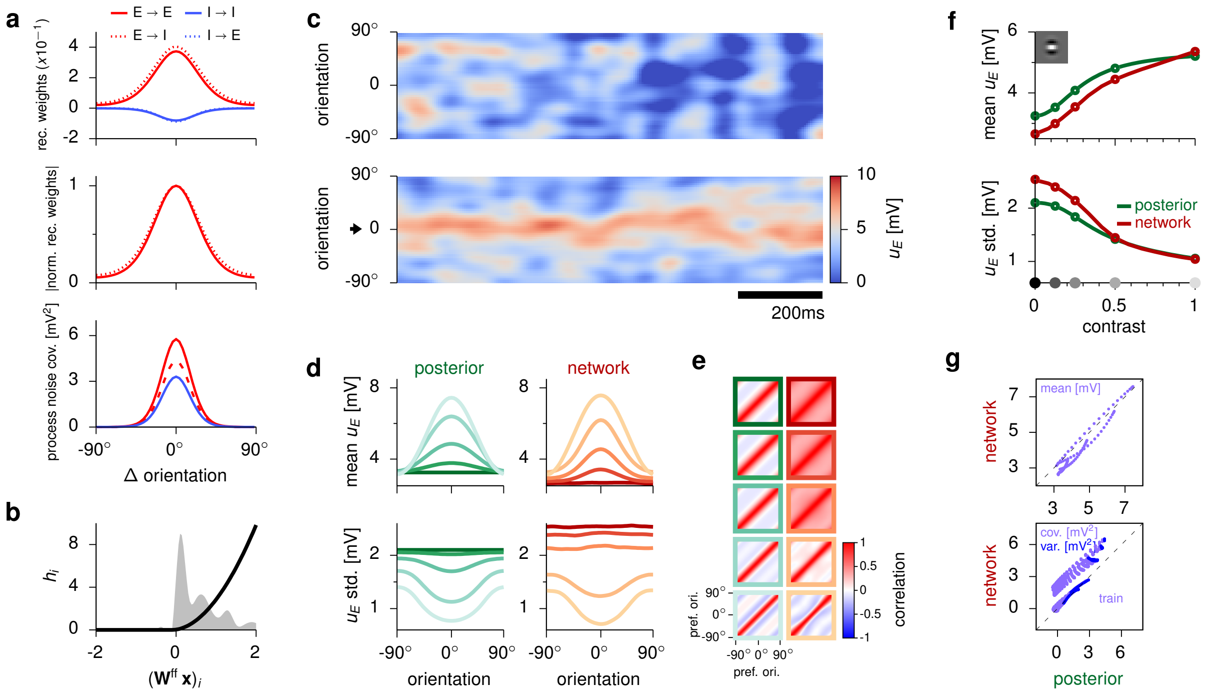


**Fig. S9 | Control network: no explicit slowness penalty.** Layout as in Fig. S8. Line segments between circles in **f** show generalization to the training image patch presented at untrained contrast levels (cf. Fig. 3a).

optimized network (Fig. S8, cf. Figs. 2 and 3). Therefore, it achieved a performance that was far better than the random networks' (Fig. S7a). However, it still performed significantly worse than networks optimized with Dale's principle enforced (Fig. S7a), and its *dynamics* were also qualitatively different: oscillations and transient overshoots were largely absent from it (Fig. S7b-c, Fig. S13).

### S4.4 Control networks optimized for other objectives

The optimized network presented in the main text displayed a suite of interesting, biologically relevant dynamical features. We have shown above that these features did not emerge in random networks without optimization, or with networks optimized for the same objective but without enforcing the same architectural constraints (Dale's principle) – but that they did emerge robustly when optimizing the same architecture for the same objective. However, these investigations left open the possibility that the architectural constraints of the original network (E-I structure, rotational symmetry, choice of membrane time constants, single cell nonlinearity, etc) alone might have been *sufficient* for generating the dynamical features of our interest, irrespective of our computational objective. Thus, we systematically "knocked out" or altered different terms of the original cost function (Eq. 17) to see which ones (if any) were critical for the results we report in the main text. Crucially, when training under these altered cost functions, we left everything else (network architecture, etc) in our original optimization approach unchanged.

We first tested whether the explicit slowness penalty in our cost function (Eq. 21) was necessary by setting $\epsilon_{\text{slow}} = 0$ in Eq. 17. We found that a network optimized without this term behaved largely identically to the originally optimized network (Figs. S9 and S13). As we note in the main text (Online Methods), this could be attributed to the fact that our optimization implicitly encouraged fast sampling by default, simply by using a finite averaging window for computing average moments of network responses, and in particular by including samples immediately or shortly following stimulus onset (Section S2).

Next, we tested whether the requirement to modulate the (co)variances of network responses in order to match those of the ideal observer's posterior was necessary for our main results. This was particularly important because modulations of response variability are a hallmark of a sampling-based probabilistic inference strategy (Orbán et al., 2016). Therefore, if networks not required to modulate their response variability had also shown the same dynamical features that we found in the originally optimized network that would have called into question how specific those features were for sampling-based inference. To see whether this was the case, we

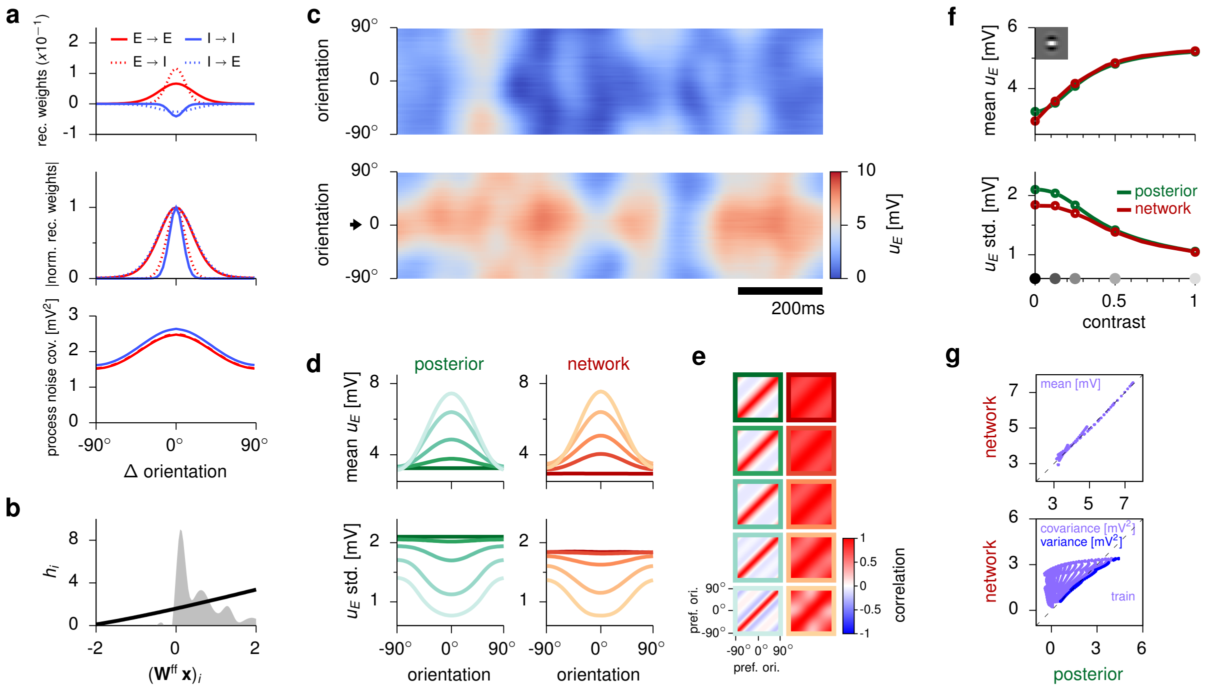


**Fig. S10 | Control network: mean and variance (but no covariance) matching.** Layout as in Fig. S8. Line segments between circles in **f** show generalization to the training image patch presented at untrained contrast levels (cf. Fig. 3a). Note complete failure to fit covariances (but not variances, or means) in **e** and **g**.

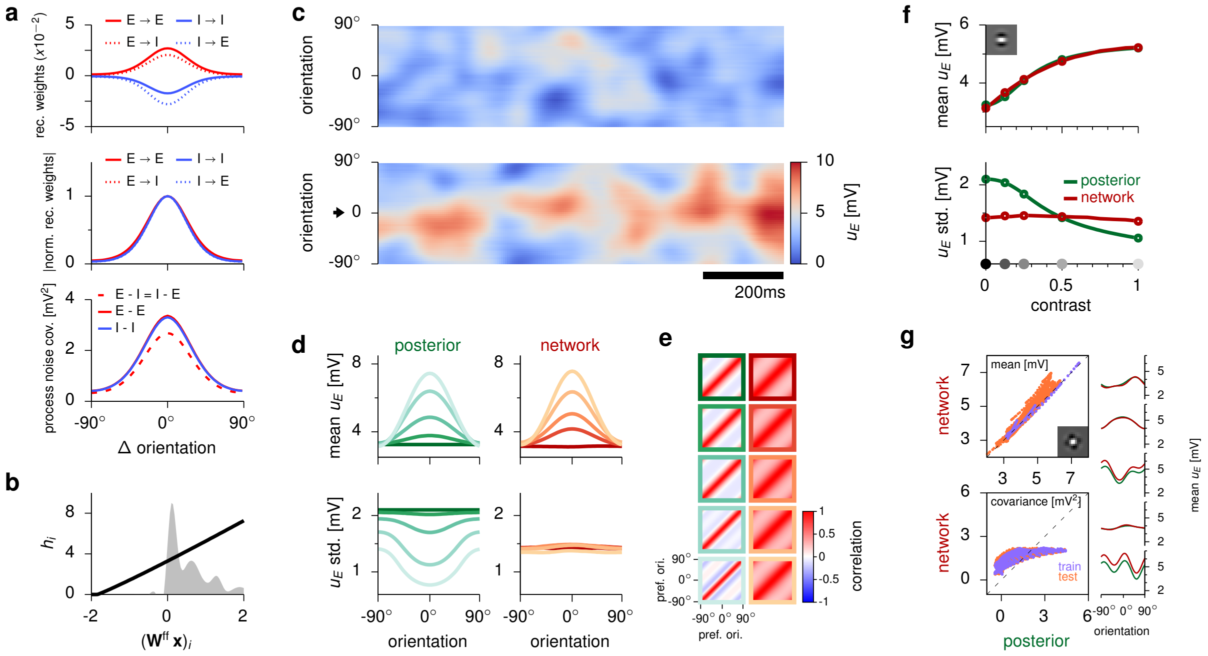


**Fig. S11 | Control network: mean matching only (i.e. no variance or covariance matching).** Layout as in Fig. S8. Line segments between circles in **f** show generalization to the training image patch presented at untrained contrast levels (cf. Fig. 3a). Orange dots in **g** correspond to novel, untrained stimuli in the test set (see inset in top panel for a representative stimulus; cf. Fig. 3b). No distinction is made (either for training or test stimuli) between variances and covariances. Insets in **g** (right) show examples of generalization to novel test stimuli (cf. Fig. 3c). Each row corresponds to a different stimulus, showing the corresponding means of latent variables under the GSM posterior (green) and stationary responses in the network (red). Note complete failure to fit (co)variances (but not means) in **d**-**g**.

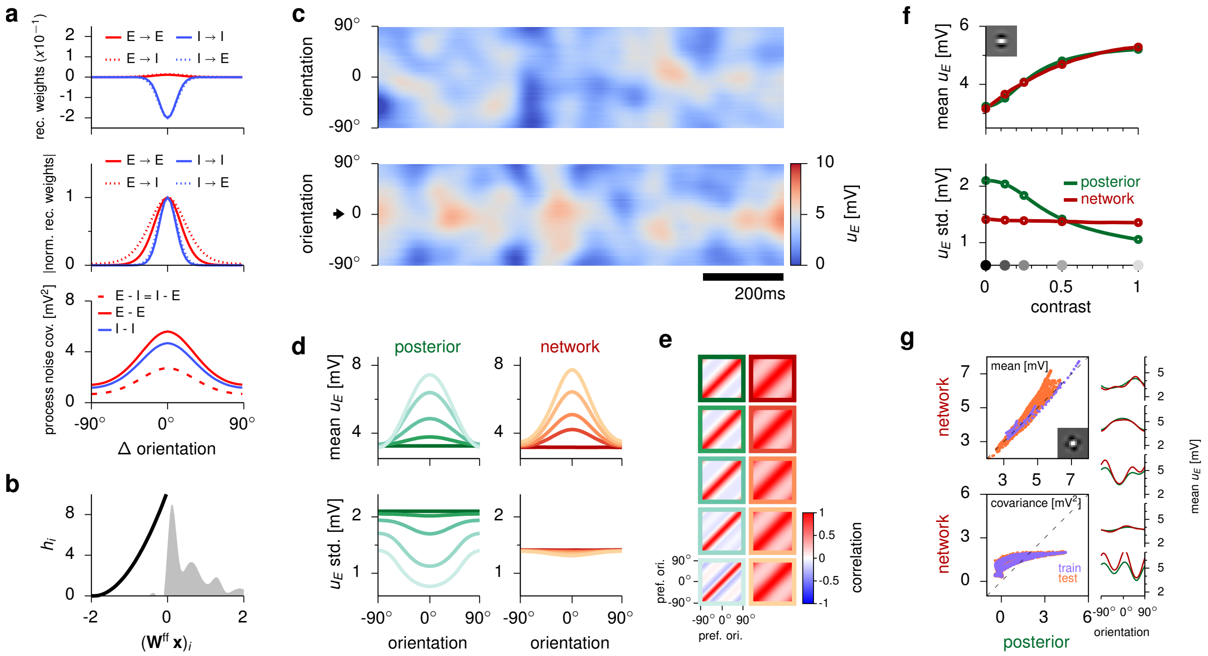

**Fig. S12 | Control network: enforcing constant Fano factors.** Layout as in Fig. S8. Line segments between circles in **f** show generalization to the training image patch presented at untrained contrast levels (cf. Fig. 3a). Orange dots in **g** correspond to novel, untrained stimuli in the test set (see inset in top panel for a representative stimulus; cf. Fig. 3b). No distinction is made (either for training or test stimuli) between variances and covariances. Insets in **g** (right) show examples of generalization to novel test stimuli (cf. Fig. 3c). Each row corresponds to a different stimulus, showing the corresponding means of latent variables under the GSM posterior (green) and stationary responses in the network (red). Note complete failure to fit (co)variances (but not means) in **d**-**g**.

either set $\epsilon_{\text{cov}} = 0$, thus foregoing the requirement of matching covariances but still requiring the network to match response means and variances (Fig. S10), or we set both $\epsilon_{\text{var}} = 0$ and $\epsilon_{\text{cov}} = 0$ in Eq. 17, thus requiring the network to match only response means (Fig. S11). (In both cases, we also set $\epsilon_{\text{slow}} = 0$ which did not have any strong effect on the network's dynamics, as shown above; Figs. S9 and S13.)

After optimization, these networks developed weak connection weights (the latter one an order of magnitude weaker than in the originally optimized network; Fig. S10a and Fig. S11a, top, cf. Fig. S3a), with virtually identical widths for E and I inputs onto both E and I cells (Fig. S10a and Fig. S11a, middle; cf. Fig. S3a), and an almost linear input transformation (Fig. S11b; cf. Fig. S3b). As a result, in both cases, we found that neither network modulated the moments it was not explicitly required to match (Fig. S10e and g, Fig. S11d-g). In fact, the network matching only the means was so weakly coupled that, as expected from an essentially feed-forward network, its response covariance tuning simply reflected its process noise covariance (cf. Fig. S11e and a, bottom). In contrast, in the network that did not match covariances, optimization found a solution such that all neurons were highly correlated, with a single, global mode of fluctuation. This solution seemed to be sufficient to achieve the target variances (but not covariances). These results meant that tuning curves (based on mean rates) in both networks were similar to those in the originally optimized network (Fig. S13a), but Fano factors were barely modulated in the network with only mean matching (Fig. S13b). This indicated that modulating (co)variances was not simply an unavoidable byproduct of the required mean modulation and the particular network architecture of our choice. Critically, unlike the originally optimized network, neither of these networks showed discernible oscillations or transient overshoots Fig. S13c-e). Therefore, within the space of models we tested, the objective to modulate response variability seemed necessary for obtaining the dynamical features reported in the main text.

While the control network with only mean matching displayed only weak stimulus-dependent modulation of Fano factors (Fig. S13b), it was not specifically trained to achieve this. This was interesting, because Fano factors do need to be specifically stimulus-independent for a class of models, (linear) probabilistic population codes (PPCs; Ma et al., 2006), that provide a conceptually very different link between neural variability and the representation of uncertainty than that provided by sampling, which we pursue here (Fiser et al., 2010; Orbán et al., 2016; Shivkumar et al., 2018). Therefore, we used our optimization-based approach to directly compare

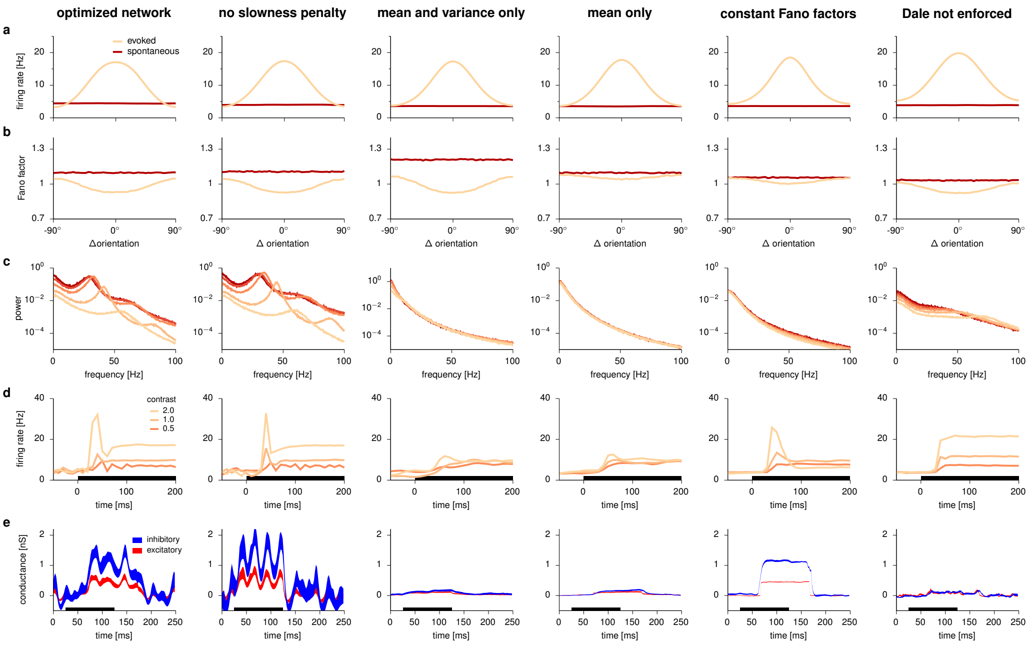


**Fig. S13 | Comparison of neural dynamics between the originally optimized network and the control networks. a-e,** As Fig. 5, comparing the originally optimized network of the main paper (left), to various control networks, from left to right: without slowness penalty, without covariance modulation (matching means and variances),
without covariance and variance modulation (matching means only), enforcing constant Fano factors, and with Dale's principle not enforced. For ease of comparison, only stimulus-dependent power spectra are shown for the optimized network of the main paper, without showing the dependence of gamma peak frequency on contrast (cf. Fig. 5c, middle), as most control networks had no discernible gamma peaks.

the circuit dynamics required by PPCs to those of our originally optimized network implementing sampling. For this, we trained a further control network whose goal was to match the mean modulation of the control network (resulting in realistic tuning curves), while keeping Fano factors constant. We achieved this by devising a set of target covariances that would result (together with the target mean responses used by all other networks) in constant Fano factors – assuming a doubly stochastic Poisson process (i.e. Cox process) for spike generation and an exponentially decaying autocorrelation function, for which an analytic expression relating rate moments and Fano factors exists (see Hennequin and Lengyel, 2016, for details). Training then proceeded exactly as for the other networks, with the same $\epsilon$ parameters in the cost function as for the originally optimized network, only employing the new covariance targets.

The resulting optimized network made use of strong inhibitory connections, large shared process noise (Fig. S12a), and strongly modulated inputs (Fig. S12b). The network was able to match mean responses in the training set and to generalize to novel stimuli (Fig. S12d, f-g), while keeping Fano factors relatively constant as required (Fig. S13b). (For consistency with previous results, we obtained Fano factors by numerically simulating the same type of doubly stochastic Gamma process as in the other networks, Section S3.4, thus violating the assumptions under which we computed the target covariances of the network – hence the remaining small modulations of Fano factors.) Critically, while the training procedure was the same as for the original network, only differing in the required variability modulation provided by the targets, this control network displayed no gamma-band oscillations (Fig. S13c). Inhibition-dominated transients did emerge, but were weaker than in the original network (Fig. S13d and e).

### S5 Mathematical analyses of oscillations and transients

In order to gain an understanding of the computational benefits of oscillations and transient overshoots in the network, we constructed a series of simplified, analytically tractable systems in which these features could be selectively controlled. In line with our training objective (Eq. 17) and all other analyses presented here, which were based on first- and second-order statistics, we modelled the statistics of neural responses as Gaussian processes (GPs; Williams and Rasmussen, 2006), $\mathbf{u}(t) \sim \mathrm{GP}\big(\boldsymbol{\mu}_{\mathrm{GP}}(\cdot), \boldsymbol{\Sigma}_{\mathrm{GP}}(\cdot,\cdot)\big)$, with mean, $\boldsymbol{\mu}_{\mathrm{GP}}(t) = \mathbb{E}\left[\mathbf{u}(t)\right]$, and covariance functions, $\boldsymbol{\Sigma}_{\mathrm{GP}}\left(t, t'\right) = \mathbb{C}\left[\mathbf{u}(t), \mathbf{u}\left(t'\right)\right]$, chosen to produce overshoots and/or oscillations in a controlled way.

### S5.1 Sampling accuracy of statistically stationary dynamics

We want to know how well a finite number of samples (i.e. responses from the network) – or more precisely: a continuum of temporally correlated samples collected over a finite time window of length $T$ – can represent the target distribution $\mathcal{P}$. For this, we quantify the discrepancy between the relevant target distribution, $\mathcal{P}$, and the (time-marginalized) distribution of responses produced by the network over a finite time window $T$, $\mathcal{Q}_T$. Specifically, we use the symmetrized Kullback-Leibler divergence ($\mathrm{D}_{\mathrm{SKL}}$) defined for two probability distributions $\mathcal{P}$ and $\mathcal{Q}_T$ as:

$$\mathrm{D}_{\mathrm{SKL}}\left[\mathcal{P}\|\mathcal{Q}_T\right] = \frac{1}{2}\left[\int \mathcal{P}(\mathbf{u}) \log\left(\frac{\mathcal{P}(\mathbf{u})}{\mathcal{Q}_T(\mathbf{u})}\right) \mathrm{d}\mathbf{u} + \int \mathcal{Q}_T(\mathbf{u}) \log\left(\frac{\mathcal{Q}_T(\mathbf{u})}{\mathcal{P}(\mathbf{u})}\right) \mathrm{d}\mathbf{u}\right] \tag{S15}$$

where $\mathrm{D}_{\mathrm{SKL}}\left[\mathcal{P}\|\mathcal{Q}_T\right] \geq 0$, with equality achieved if and only if $\mathcal{P}(\mathbf{u}) = \mathcal{Q}_T(\mathbf{u})\ \forall \mathbf{u}$. Note that all pairs of distributions studied here are approximately Gaussian: $\mathcal{P}$ is (approximately) Gaussian given the construction of our generative model (Eqs. 3, 6 and 7), and $\mathcal{Q}_T$ is also Gaussian under our GP approximation (see above). Therefore, both $\mathcal{P}$ and $\mathcal{Q}_T$ can be considered to be fully characterized by their means and covariances: $\boldsymbol{\mu}_{\mathcal{P}}$ and $\boldsymbol{\Sigma}_{\mathcal{P}}$ for the target distribution, and $\boldsymbol{\mu}_{\mathcal{Q}}(T) = \frac{1}{T}\int_0^T \mathbf{u}(t)\,\mathrm{d}t$ and $\boldsymbol{\Sigma}_{\mathcal{Q}}(T) = \frac{1}{T}\int_0^T \mathbf{u}(t)\ \mathbf{u}^{\mathsf{T}}(t)\,\mathrm{d}t - \boldsymbol{\mu}_{\mathcal{Q}}\,\boldsymbol{\mu}_{\mathcal{Q}}^{\mathsf{T}}$ for the response distribution, respectively. With these definitions, Eq. S15 takes the following approximate form:

$$\mathrm{D}_{\mathrm{SKL}}\left[\mathcal{P}\|\mathcal{Q}_T\right] \approx \frac{1}{4}\left[\boldsymbol{\mathcal{E}}^{\mathsf{T}}(T)\left(\boldsymbol{\Sigma}_{\mathcal{P}}^{-1} + \boldsymbol{\Sigma}_{\mathcal{Q}}^{-1}(T)\right)\boldsymbol{\mathcal{E}}(T) + \mathrm{Tr}\left(\boldsymbol{\Sigma}_{\mathcal{P}}^{-1}\,\boldsymbol{\Sigma}_{\mathcal{Q}}(T)\right) + \mathrm{Tr}\left(\boldsymbol{\Sigma}_{\mathcal{Q}}^{-1}(T)\ \boldsymbol{\Sigma}_{\mathcal{P}}\right) - 2\dim(\mathbf{u})\right] \tag{S16}$$

where $\boldsymbol{\mathcal{E}}(T) = \boldsymbol{\mu}_{\mathcal{Q}}(T) - \boldsymbol{\mu}_{\mathcal{P}}$ is the error in the sample mean of responses.

We now consider the special case when $\dim(\mathbf{u}) = 1$, i.e. $\mathbf{u}$ is a scalar $u$ (either a single cell, or an arbitrary 1-dimensional projection of the network's activity – see Footnote 3 at the end of this section as well as Sections S5.2.2–S5.2.4 for results pertaining to multivariate $\mathbf{u}$), and thus so are the corresponding means and variances in Eq. S16. Note that $\mu_{\mathcal{Q}}(T)$ and $\sigma^2_{\mathcal{Q}}(T)$ themselves are random variables (as they are based on the stochastic responses, $u(t)$, of the network). Thus, we compute the *expected* divergence between $\mathcal{P}$ and $\mathcal{Q}_T$, $\overline{\mathrm{D}}_{\mathrm{SKL}}\left[\mathcal{P}\|\mathcal{Q}_T\right] = \mathbb{E}\left[\mathrm{D}_{\mathrm{SKL}}\left[\mathcal{P}\|\mathcal{Q}_T\right]\right]$, over the distribution of $\mu_{\mathcal{Q}}(T)$ and $\sigma^2_{\mathcal{Q}}(T)$, given the "true" mean and covariance function of the GP, $\mu_{\mathrm{GP}}(t)$ and $\sigma^2_{\mathrm{GP}}\left(t, t'\right)$, producing the samples comprising $\mathcal{Q}_T$:

$$\overline{\mathrm{D}}_{\mathrm{SKL}}\left[\mathcal{P}\|\mathcal{Q}_T\right] \approx \frac{1}{4}\left[\mathbb{E}\left[\mathcal{E}^2(T)\left(\frac{1}{\sigma^2_{\mathcal{P}}} + \frac{1}{\sigma^2_{\mathcal{Q}}(T)}\right)\right] + \mathbb{E}\left[\sigma^2_{\mathcal{Q}}(T)\right]\frac{1}{\sigma^2_{\mathcal{P}}} + \mathbb{E}\left[\frac{1}{\sigma^2_{\mathcal{Q}}(T)}\right]\sigma^2_{\mathcal{P}} - 2\right] \tag{S17}$$

Here we study the dynamics of our network in the statistically stationary limit, i.e. past burn-in (see Section S5.3 for an analysis of transients). Formally, this means that $\mathbb{E}\left[u(t)\right] = \mu_{\mathrm{GP}}(t) = \mu_{\mathrm{GP}}$, $\mathbb{V}\left[u(t)\right] = \sigma^2_{\mathrm{GP}}(t,t) = \sigma^2_{\mathrm{GP}}$, and $\mathbb{C}\left[u(t), u\left(t'\right)\right] = \sigma^2_{\mathrm{GP}}\,C_{\mathrm{GP}}\left(t - t'\right)\ \forall t, t'$, where $C_{\mathrm{GP}}(\Delta t) = C_{\mathrm{GP}}(-\Delta t)$ is the autocorrelation function of the GP characterizing network responses. As we are primarily interested in how sampling accuracy specifically depends on the *dynamics* of neural responses, we also assume that the (time-marginalized) stationary distribution of responses has already been matched to the target distribution. This implies that $\mu_{\mathcal{P}} = \mathbb{E}\left[\mu_{\mathcal{Q}}(T)\right] = \mu_{\mathrm{GP}}$ and $\sigma^2_{\mathcal{P}} = \mathbb{E}\left[\sigma^2_{\mathcal{Q}}(T)\right] = \sigma^2_{\mathrm{GP}}\left(1 - \overline{C}_{\mathrm{GP}}(T)\right)$, where

$$\overline{C}_{\mathrm{GP}}(T) = \frac{1}{T^2}\int_0^T\int_0^T C_{\mathrm{GP}}\left(t - t'\right)\mathrm{d}t\,\mathrm{d}t' \tag{S18}$$

is the average autocorrelation of the network, i.e. the normalized area under the autocorrelation function. Therefore, note that moment matching does not constrain $C_{\mathrm{GP}}$ in any way, as any value of $\overline{C}_{\mathrm{GP}}(T)$ (smaller than 1) can be compensated for by the appropriate choice of $\sigma^2_{\mathrm{GP}}$ to match $\mathbb{E}\left[\sigma^2_{\mathcal{Q}}(T)\right]$ to $\sigma^2_{\mathcal{P}}$. For simplicity, we further assume that the sample mean, $\mu_{\mathcal{Q}}(T)$, and variance of network responses, $\sigma^2_{\mathcal{Q}}(T)$, are statistically independent given $\mu_{\mathrm{GP}}$, $\sigma^2_{\mathrm{GP}}$ and $C_{\mathrm{GP}}$. (Formally, this only holds for i.i.d. normal random variables, while for correlated variables produced by a stationary process, such as our sequence of network responses, only zero covariance can be proven.[2]) In addition, we ignore the effects of variability in the sample variance, $\sigma^2_{\mathcal{Q}}(T)$. Under these assumptions Eq. S17 (or at least a lower bound thereof) can be written as

$$\overline{\mathrm{D}}_{\mathrm{SKL}}\left[\mathcal{P}\|\mathcal{Q}_T\right] \approx \frac{\overline{\mathcal{E}}^2(T)}{2\,\sigma^2_{\mathcal{P}}} \tag{S19}$$

where $\overline{\mathcal{E}}^2(T) = \mathbb{E}\left[\mathcal{E}^2(T)\right] = \mathbb{V}\left[\mu_{\mathcal{Q}}(T)\right]$ is the mean squared error of the sample mean of responses, which is equal to the variance of the sample mean under our moment matching assumption (see above). Importantly, in this case, $\overline{\mathcal{E}}^2(T)$ can be shown to depend directly on the average autocorrelation of network responses:

$$\overline{\mathcal{E}}^2(T) = \sigma^2_{\mathcal{P}}\,\frac{\overline{C}_{\mathrm{GP}}(T)}{1-\overline{C}_{\mathrm{GP}}(T)} \tag{S20}$$

Substituting Eq. S20 to Eq. S19 yields[3]

$$\overline{\mathrm{D}}_{\mathrm{SKL}}\left[\mathcal{P}\|\mathcal{Q}_T\right] \approx \frac{1}{2}\,\frac{\overline{C}_{\mathrm{GP}}(T)}{1-\overline{C}_{\mathrm{GP}}(T)} \tag{S21}$$

As the area under the autocorrelogram must be finite, Eqs. S18 and S21 imply that $\overline{C}_{\mathrm{GP}}(T)$ and thus $\overline{\mathrm{D}}_{\mathrm{SKL}}$ tends to 0 as $T \to \infty$. In other words, in line with intuition, sufficiently long time windows allow the target distribution to be represented with arbitrary precision (once the stationary distribution is matched).

### S5.2 Oscillations

In this section we first analyze how oscillations in neural responses affect sampling accuracy in general, and then study how the dynamics of a network lead to such oscillations, and in particular to oscillations that are beneficial for sampling.

#### S5.2.1 Oscillations and sampling accuracy

The fact that there is a tight relationship between the average autocorrelation and sampling accuracy (Eq. S21) reveals a fundamental benefit of oscillations for sampling. Oscillations make the total (signed) area under the autocorrelogram smaller because an oscillatory autocorrelogram has as its envelope (and thus never exceeds) its non-oscillatory counterpart (Eq. S14).

In order to illustrate this fundamental effect of oscillations, in Fig. 6, we considered a stationary, moment-matched GP (see previous section) with an autocorrelation function that had the following parametric form:

$$C_{\mathrm{GP}}(\Delta t) = \left[(1-\alpha) + \alpha\,\cos\!\left(2\pi f\,|\Delta t|\right)\right]\left(1 + \frac{|\Delta t|}{\tau}\right)\,e^{-|\Delta t|/\tau} \tag{S22}$$

Note that this was a special case of the parametric function we fitted to the autocorrelogram of actual network responses (Eq. S12) with the time constant of process noise being the same as the membrane time constant:

[2] Alternatively, the results below can also be obtained without assuming the independence of the sample mean and variance, by using the expected (non-symmetrized) Kullback-Leibler divergence, $\overline{\mathrm{D}}_{\mathrm{KL}}\left(\mathcal{Q}\|\mathcal{P}\right)$, to measure the sampling accuracy of the network.

[3] Eq. S21 can be straightforwardly generalized to the original case of multivariate $\mathbf{u}$:

$$\overline{\mathrm{D}}_{\mathrm{SKL}}\left[\mathcal{P}\|\mathcal{Q}_T\right] \approx \frac{1}{2}\mathrm{Tr}\left[\left(\mathbf{I}-\overline{\mathbf{C}}_{\mathrm{GP}}(T)\right)^{-1}\overline{\mathbf{C}}_{\mathrm{GP}}(T)\right]$$

where, in analogy with Eq. S18,

$$\overline{\mathbf{C}}_{\mathrm{GP}}(T) = \frac{1}{T^2}\int_0^T\!\!\int_0^T \mathbf{C}_{\mathrm{GP}}\left(t-t'\right)\mathrm{d}t\,\mathrm{d}t'$$

such that

$$\mathbb{C}\left[\mathbf{u}(t),\mathbf{u}(t')\right] = \boldsymbol{\Sigma}^{1/2}_{\mathrm{GP}}\,\mathbf{C}_{\mathrm{GP}}\left(t-t'\right)\,\boldsymbol{\Sigma}^{1/2}_{\mathrm{GP}} \quad \text{with} \quad \mathbf{C}_{\mathrm{GP}}(-\Delta t) = \mathbf{C}^{\mathsf{T}}_{\mathrm{GP}}(\Delta t) \quad \text{and} \quad \mathbf{C}_{\mathrm{GP}}(0) = \mathbf{I}$$

in the stationary case.

$\tau_\eta = \tau$. (This was relevant for our network, as we had $\tau_E = \tau_\eta = 20$ ms for excitatory neurons.) Thus, just as in Section S3.5, $\alpha \in [0,1]$ determined the strength of oscillations in the system (which occurred at frequency $f$), with the $\alpha = 0$ case corresponding to the autocorrelation of an isolated single neuron. With this parametrization, it is easy to show that $\overline{\mathrm{D}}_{\mathrm{SKL}}\left[\mathcal{P}\|\mathcal{Q}\right] \to 0$ in Eq. S21 (because $\overline{C}_{\mathrm{GP}}(T) \to 0$) as $f \to \infty$, provided that $\alpha > 0$. Conversely, the more oscillatory the system is (i.e. the larger $\alpha$ is), the better its accuracy (the smaller its $\overline{\mathrm{D}}_{\mathrm{SKL}}\left[\mathcal{P}\|\mathcal{Q}\right]$) becomes for fixed, sufficiently high $f$.

As an example, we considered different specific values of $\alpha \in \{0, 0.2, 0.7\}$ at $f = 40$ Hz (Fig. 6a; we set $\mu_{\mathrm{GP}} = 3$ and $\sigma^2_{\mathrm{GP}} = 4$ without loss of generality as the analysis above suggested that $\overline{\mathrm{D}}_{\mathrm{SKL}}\left[\mathcal{P}\|\mathcal{Q}\right]$ is shift- and scale-invariant). We then used Eq. S17 to compute $\overline{\mathrm{D}}_{\mathrm{SKL}}\left[\mathcal{P}\|\mathcal{Q}\right]$ as a function of the length of the sampling time window, $T$, and the level of oscillatoriness, $\alpha$ (Fig. 6b).

#### S5.2.2 Oscillatoriness of high-dimensional network dynamics

In the preceding sections, we have used GPs with oscillatory autocorrelations as phenomenological models in order to understand how oscillatory components in network responses confer benefits for sampling. In this section, we build analytical insight into how the dynamics of the actual network gives rise to its spatio-temporal covariance, and its oscillatoriness, in the first place.

The first two moments of the stationary distribution of network activity are given by the steady-state solutions of the differential equations in Eqs. S5-S7:

$$\mathbf{0} = \left[\mathbf{T}^{-1}\,\boldsymbol{\Sigma}^*\right] + \left[\mathbf{T}^{-1}\,\boldsymbol{\Sigma}^*\right]^{\mathsf{T}} + \mathcal{J}\,\boldsymbol{\Sigma} + \boldsymbol{\Sigma}\,\mathcal{J}^{\mathsf{T}} \tag{S23}$$

$$\mathbf{0} = -\frac{1}{\tau_\eta}\,\boldsymbol{\Sigma}^* + \boldsymbol{\Sigma}^\eta\,\mathbf{T}^{-1} + \boldsymbol{\Sigma}^*\,\mathcal{J}^{\mathsf{T}} \tag{S24}$$

$$\frac{\mathrm{d}\boldsymbol{\Sigma}(\Delta t)}{\mathrm{d}\Delta t} = e^{-\Delta t/\tau_\eta}\,\left[\mathbf{T}^{-1}\boldsymbol{\Sigma}^*\right]^{\mathsf{T}} + \boldsymbol{\Sigma}(\Delta t)\,\mathcal{J}^{\mathsf{T}}, \qquad \forall \Delta t > 0 \tag{S25}$$

where we have dropped the primary time index $t$ everywhere as we are studying the stationary distribution here. The case $\Delta t < 0$ in Eq. S25 is resolved using $\boldsymbol{\Sigma}(-\Delta t) = \boldsymbol{\Sigma}^{\mathsf{T}}(\Delta t)$. Eq. S24 can be solved directly:

$$\boldsymbol{\Sigma}^* = \boldsymbol{\Sigma}^\eta\,\mathbf{T}^{-1}\,\left(\mathbf{I}/\tau_\eta - \mathcal{J}^{\mathsf{T}}\right)^{-1} \tag{S26}$$

and so Eq. S25 can be rewritten for convenience as:

$$\frac{\mathrm{d}\boldsymbol{\Sigma}^{\mathsf{T}}(\Delta t)}{\mathrm{d}\Delta t} = e^{-\Delta t/\tau_\eta}\,\mathbf{M} + \mathcal{J}\,\boldsymbol{\Sigma}^{\mathsf{T}}(\Delta t), \quad \text{where} \quad \mathbf{M} = \mathbf{T}^{-1}\,\boldsymbol{\Sigma}^* = \mathbf{T}^{-1}\,\boldsymbol{\Sigma}^\eta\,\mathbf{T}^{-1}\,\left(\mathbf{I}/\tau_\eta - \mathcal{J}^{\mathsf{T}}\right)^{-1} \tag{S27}$$

Now, we use the Laplace transform to obtain:

$$\mathcal{L}\left\{\boldsymbol{\Sigma}^{\mathsf{T}}\right\}(s) = (s\,\mathbf{I} - \mathcal{J})^{-1}\,\left[\boldsymbol{\Sigma}^{\mathsf{T}}(0) + \frac{1}{s + 1/\tau_\eta}\mathbf{M}\right] \tag{S28}$$

which is then solved by using the inverse Laplace transform:

$$\boldsymbol{\Sigma}^{\mathsf{T}}(\Delta t) = e^{\Delta t\,\mathcal{J}} * \left[\delta(\Delta t)\,\boldsymbol{\Sigma}^{\mathsf{T}}(0) + e^{-\Delta t/\tau_\eta}\,\mathrm{H}(\Delta t)\,\mathbf{M}\right] \tag{S29}$$

where $\mathrm{H}(\cdot)$ is the Heaviside function. This yields:

$$\boldsymbol{\Sigma}^{\mathsf{T}}(\Delta t) = e^{\Delta t\,\mathcal{J}}\,\boldsymbol{\Sigma}^{\mathsf{T}}(0) + \left(\mathbf{I}/\tau_\eta + \mathcal{J}\right)^{-1}\,\left[e^{\mathcal{J}\,\Delta t} - e^{-\Delta t\,\mathbf{I}/\tau_\eta}\right]\,\mathbf{M} \tag{S30}$$

and therefore

$$\boldsymbol{\Sigma}(\Delta t) = \boldsymbol{\Sigma}\,e^{\Delta t\,\mathcal{J}^{\mathsf{T}}} + \boldsymbol{\Sigma}^{*\mathsf{T}}\,\mathbf{T}^{-1}\,\left[e^{\Delta t\,\mathcal{J}^{\mathsf{T}}} - e^{-\Delta t\,\mathbf{I}/\tau_\eta}\right]\,\left(\mathbf{I}/\tau_\eta + \mathcal{J}^{\mathsf{T}}\right)^{-1} \tag{S31}$$

Finally, by diagonalizing the Jacobian $\mathcal{J} = \mathbf{V}\,\Lambda\,\mathbf{V}^{-1}$, where $\Lambda$ is a diagonal matrix of eigenvalues and $\mathbf{V}$ is the matrix whose columns contain the eigenvectors of $\mathcal{J}$, one obtains:

$$\boldsymbol{\Sigma}(\Delta t) = \boldsymbol{\Sigma}\,\mathbf{V}^{-\mathsf{T}}\,e^{\Delta t\,\Lambda}\,\mathbf{V}^{\mathsf{T}} + \boldsymbol{\Sigma}^{*\mathsf{T}}\,\mathbf{T}^{-1}\,\mathbf{V}^{-\mathsf{T}}\,\left[e^{\Delta t\,\Lambda} - e^{-\Delta t\,\mathbf{I}/\tau_\eta}\right]\,\left(\mathbf{I}/\tau_\eta + \Lambda\right)^{-1}\,\mathbf{V}^{\mathsf{T}} \tag{S32}$$

Eq. S32 exposes how the eigenvalues of the Jacobian (diagonal of $\Lambda$) with non-zero imaginary parts (first term of the equation and first term in square brackets) contribute to oscillatoriness in the sense of Eqs. S12 and S22. (Oscillations in the correlogram may be damped if the corresponding eigenvalues have comparatively large negative real parts.) This complex component competes with a pure (real) exponential decay (second term in square brackets), reflecting the autocorrelation of process noise. Thus, overall, Eq. S32 results in a partial degree of oscillatoriness (modulated by properties of the Jacobian), as seen in the real network.

##### S5.2.3 Optimal oscillatory high-dimensional dynamics

To understand how oscillatory activity in high-dimensional dynamics can be optimised for sampling, we use (an approximation of) our original measure of accuracy for multivariate distributions, the multivariate equivalent of Eq. S19:

$$\overline{\mathrm{D}}_{\mathrm{SKL}}\left[\mathcal{P}\|\mathcal{Q}_T\right] \approx \frac{1}{2}\mathbb{E}\left[\boldsymbol{\mathcal{E}}^{\mathsf{T}}(T)\ \boldsymbol{\Sigma}_{\mathcal{P}}^{-1}\ \boldsymbol{\mathcal{E}}(T)\right] \tag{S33}$$

Using the equivalence of a scalar expression and its own trace, and the cyclic property of the trace, this can be rewritten as

$$\overline{\mathrm{D}}_{\mathrm{SKL}}\left[\mathcal{P}\|\mathcal{Q}_T\right] \approx \frac{1}{2}\mathrm{Tr}\left[\mathbb{C}\left[\boldsymbol{\mu}_{\mathcal{Q}}(T)\right]\ \boldsymbol{\Sigma}_{\mathcal{P}}^{-1}\right] \tag{S34}$$

Here we extend the results of Aitchison et al. (2018) on a network receiving fast (temporally white) process noise, to the case of temporally correlated noise, which is more relevant to our optimized network. For analytical tractability, we have made two simplifying assumptions regarding process noise: that its temporal correlations are short (though non-negligible) compared to the intrinsic time constants of the network, and that it is spatially white (see below for mathematical definitions). Moreover, we assume that the dynamics of the inhibitory neurons are fast enough that their responses become slaved to those of the E neurons. This implies that we can describe our network with effective dynamics in the E population alone, i.e. the same space over which the target distributions are defined. In turn, this will help establish an analytical link between network dynamics and sampling performance.

In the following, we assume stationarity and fast process noise, $\tau_\eta \mathcal{J} \ll \mathbf{I}$, expanding expressions to first order in $\tau_\eta$, and define $\mathbf{Q}_\circ = 2\,\tau_\eta\,\mathbf{T}^{-1}\,\boldsymbol{\Sigma}^\eta\,\mathbf{T}^{-1}$. With these assumptions and definition, after substituting Eq. S26 into Eq. S31, we can rewrite the lagged-covariance of the network as:

$$\boldsymbol{\Sigma}(\Delta t) \approx \left(\boldsymbol{\Sigma} + \frac{\tau_\eta}{2}\,\mathbf{Q}_\circ\right) e^{\Delta t \mathcal{J}^{\mathsf{T}}} - \frac{\tau_\eta}{2}\,\mathbf{Q}_\circ\, e^{-\Delta t/\tau_\eta} \tag{S35}$$

Under our assumptions, we can also establish a relationship between $\mathcal{J}$ and $\boldsymbol{\Sigma}$ based on Eq. S23, via the following Lyapunov equation:

$$0 \approx \mathbf{Q}_\circ + \mathcal{J}\left(\boldsymbol{\Sigma} + \frac{\tau_\eta}{2}\,\mathbf{Q}_\circ\right) + \left(\boldsymbol{\Sigma} + \frac{\tau_\eta}{2}\,\mathbf{Q}_\circ\right)\mathcal{J}^{\mathsf{T}} \tag{S36}$$

Analogously to the case of temporally white process noise (Aitchison et al., 2018), we can parameterize any solution to the Lyapunov equation Eq. S36 as:

$$\mathcal{J} = -\left(\frac{1}{2}\,\mathbf{Q}_\circ + \mathbf{S}\right)\left(\boldsymbol{\Sigma} + \frac{\tau_\eta}{2}\,\mathbf{Q}_\circ\right)^{-1} \tag{S37}$$

where $\mathbf{S}$ is an arbitrary skew-symmetric matrix, which means that it can be diagonalized as

$$\mathbf{S} = \mathbf{U}\,\boldsymbol{\Omega}^{-1}\,\mathbf{U}^{\star} \tag{S38}$$

where $\mathbf{U}$ is a (complex) unitary matrix of eigenvectors, $\mathbf{U}^{\star}$ denotes its conjugate transpose, and $\boldsymbol{\Omega}$ is a diagonal matrix containing the inverses of the purely imaginary (or zero) eigenvalues of $\mathbf{S}$.

Eqs. S35 and S37 allow us to rewrite the covariance of $\boldsymbol{\mu}_{\mathcal{Q}}(T)$, the quantity relevant for $\overline{\mathrm{D}}_{\mathrm{SKL}}\left[\mathcal{P}\|\mathcal{Q}_T\right]$ (Eq. S34), as:

$$\mathbb{C}\left[\boldsymbol{\mu}_{\mathcal{Q}}(T)\right] = \frac{1}{T^2}\int_0^T\!\!\int_0^T \boldsymbol{\Sigma}(t-t')\,\mathrm{d}t\,\mathrm{d}t' \tag{S39}$$

$$\approx \frac{1}{T}\left[\left(\boldsymbol{\Sigma} + \frac{\tau_\eta}{2}\,\mathbf{Q}_\circ\right)\left(\frac{1}{2}\,\mathbf{Q}_\circ - \mathbf{S}\right)^{-1}\mathbf{Q}_\circ\left(\frac{1}{2}\,\mathbf{Q}_\circ + \mathbf{S}\right)^{-1}\left(\boldsymbol{\Sigma} + \frac{\tau_\eta}{2}\,\mathbf{Q}_\circ\right)\right] \tag{S40}$$

where we have assumed large $T$. To simplify the analysis, we follow Aitchison et al. (2018) and set $\mathbf{\Sigma}_\eta = \sigma^2\,\mathbf{I}$ and $\mathbf{Q}_\circ = 2\,\frac{\tau_\eta\,\sigma^2}{\tau^2}\,\mathbf{I}$. We thus have:

$$\mathbb{C}\left[\boldsymbol{\mu}_{\mathcal{Q}}(T)\right] \approx \frac{2\,\tau_\eta\,\sigma^2}{\tau^2\,T}\left[\left(\mathbf{\Sigma} + \frac{\tau_\eta^2\,\sigma^2}{\tau^2}\,\mathbf{I}\right)\left(\frac{\tau_\eta\,\sigma^2}{\tau^2}\,\mathbf{I} - \mathbf{S}\right)^{-1}\left(\frac{\tau_\eta\,\sigma^2}{\tau^2}\,\mathbf{I} + \mathbf{S}\right)^{-1}\left(\mathbf{\Sigma} + \frac{\tau_\eta^2\,\sigma^2}{\tau^2}\,\mathbf{I}\right)\right] \tag{S41}$$

which, to first order in $\tau_\eta$, reduces to

$$\mathbb{C}\left[\boldsymbol{\mu}_{\mathcal{Q}}(T)\right] \approx -\frac{2\,\tau_\eta\,\sigma^2}{\tau^2\,T}\,\mathbf{\Sigma}\,\mathbf{S}^{-2}\,\mathbf{\Sigma} \tag{S42}$$

For the moment matched case, $\mathbf{\Sigma}_{\mathcal{P}} = \mathbf{\Sigma}_{\mathcal{Q}} = \mathbf{\Sigma}^{1/2}\left(\mathbf{I} - \overline{\mathbf{C}}(T)\right)\mathbf{\Sigma}^{1/2} \approx \mathbf{\Sigma}$ (for sufficiently large $T$), which means

$$\mathbb{C}\left[\boldsymbol{\mu}_{\mathcal{Q}}(T)\right] \approx -\frac{2\,\tau_\eta\,\sigma^2}{\tau^2\,T}\,\mathbf{\Sigma}_{\mathcal{P}}\,\mathbf{S}^{-2}\,\mathbf{\Sigma}_{\mathcal{P}} \tag{S43}$$

We then substitute Eq. S43 into Eq. S34 to obtain

$$\overline{\mathrm{D}}_{\mathrm{SKL}}\left[\mathcal{P}\|\mathcal{Q}_T\right] \approx -\frac{\tau_\eta\,\sigma^2}{\tau^2\,T}\mathrm{Tr}\left[\mathbf{\Sigma}_{\mathcal{P}}\,\mathbf{S}^{-2}\right] \tag{S44}$$

Therefore, optimizing the network amounts to finding the $\mathbf{S}$ that minimizes the sampling error, $\overline{\mathrm{D}}_{\mathrm{SKL}}\left[\mathcal{P}\|\mathcal{Q}_T\right]$, given by Eq. S44. It is immediately clear that increasing the scale of $\mathbf{S}$ trivially decreases $\overline{\mathrm{D}}_{\mathrm{SKL}}\left[\mathcal{P}\|\mathcal{Q}_T\right]$. We thus ask what $\mathbf{S}$ achieves a set level of the sampling error, $\overline{\mathrm{D}}^*_{\mathrm{SKL}}$, with the minimal scale (determinant)? The optimal $\mathbf{S}$ can be obtained by solving the following optimization problem:

$$\underset{\mathbf{S}}{\operatorname{argmin}}\,\det(\mathbf{S})\ \text{s.t.}\ \overline{\mathrm{D}}_{\mathrm{SKL}}\left[\mathcal{P}\|\mathcal{Q}_T\right] = \overline{\mathrm{D}}^*_{\mathrm{SKL}} \tag{S45}$$

The solution to this optimization problem is:

$$\mathbf{S}^2 = -\frac{\tau_\eta\,\sigma^2\,N}{\tau^2\,T\,\overline{\mathrm{D}}^*_{\mathrm{SKL}}}\,\mathbf{\Sigma}_{\mathcal{P}} \tag{S46}$$

Finally, for the moment matched case we are studying here and assuming fast process noise[4], Eq. S37 can be rewritten as

$$\mathcal{J} \approx -\mathbf{S}\,\mathbf{\Sigma}_{\mathcal{P}}^{-1} \tag{S47}$$

Combining Eq. S47 with Eq. S46 and Eq. S38 allows us to express the Jacobian of the network as

$$\mathcal{J} \propto \mathbf{S}^{-1} = \mathbf{U}\,\mathbf{\Omega}\,\mathbf{U}^\star \tag{S48}$$

which is also a skew-symmetric matrix with purely imaginary (or zero) eigenvalues given by $\mathbf{\Omega}$ (though see Footnote 4). Thus, the optimal dynamics are rotational, resulting in oscillatory responses, with frequencies determined by the magnitude of the elements of $\mathbf{\Omega}$. Moreover, from Eqs. S38 and S46, we can derive a relationship between $\mathbf{\Omega}$ and $\mathbf{\Sigma}_{\mathcal{P}}$:

$$\det(\mathbf{\Sigma}_{\mathcal{P}}) \propto \det\left(\mathbf{S}^2\right) = \det{}^{-1}\left(\mathbf{\Omega}^2\right) \tag{S49}$$

This means that, overall, oscillation frequencies in the network should scale inversely with the target covariance. Specifically, at progressively higher contrast levels, as $\mathbf{\Sigma}_{\mathcal{P}}$ becomes smaller, this relationship predicts higher oscillation frequencies. This is exactly what we observe in the network (Fig. 5c).

This analysis also reveals another interesting property of the optimal dynamics. As $\mathcal{J}$ is skew-symmetric (Eq. S48 and footnote 4), not only does it have purely imaginary (or zero) eigenvalues, as we saw above, but its non-zero eigenvalues also come in positive /negative pairs of equal magnitude, with each pair defining a two-dimensional plane in which the dynamics are oscillatory. Thus, without loss of generality, in the following, we analyze the dynamics by focusing on one such plane at a time.

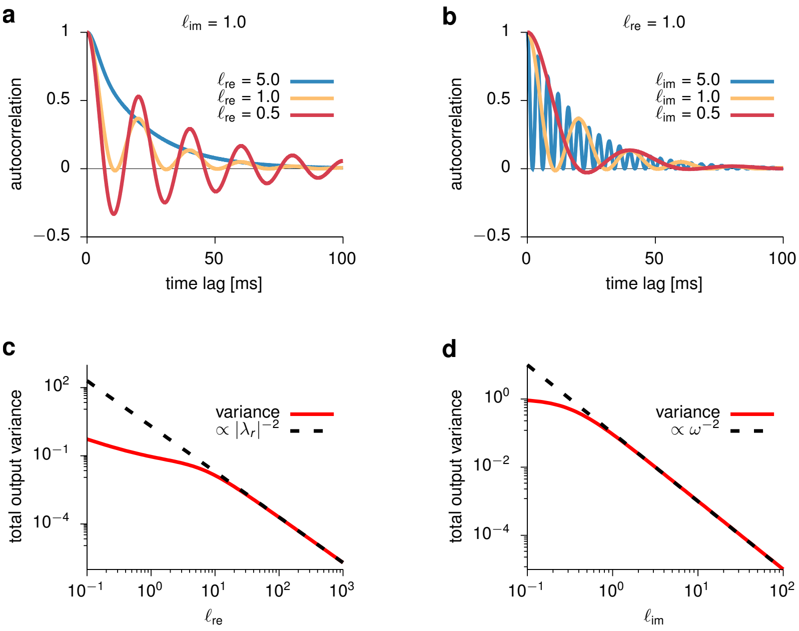


**Fig. S14 | Effects of the eigenvalues of the Jacobian in a two-dimensional system on response autocorrelation and variance. a,** Autocorrelation as a function of time lag, for different values of $\ell_{\text{re}}$ and constant $\ell_{\text{im}} = 1$ (see text following Eq. S50 for definitions). **b,** Same as **a**, but for different values of $\ell_{\text{im}}$ and constant $\ell_{\text{re}} = 1$. **c,** Total output variance as a function of $\ell_{\text{re}}$ for constant $\ell_{\text{im}} = 1$. **d,** Same as **c**, but as a function of $\ell_{\text{im}}$ for constant $\ell_{\text{re}} = 1$.

#### S5.2.4 Two-dimensional oscillatory dynamics

As seen in the previous section, optimal sampling encourages the network to operate as a collection of two-dimensional oscillators. To understand how the degree of oscillatoriness and the output variance depend on the eigenvalues of the Jacobian, we analyze here a reduced two-dimensional system, representing one of these planes in the full N-dimensional system. Each plane supports oscillations of a single frequency, given by the imaginary part of the corresponding eigenvalue ($\lambda_{\text{im}}$). Without loss of generality, we express the Jacobian as

$$\mathcal{J} = \begin{bmatrix} \lambda_{\text{re}} & -\lambda_{\text{im}} \\ \lambda_{\text{im}} & \lambda_{\text{re}} \end{bmatrix} \tag{S50}$$

with $\lambda_{\text{re}} < 0$ for stability. The eigenvalues of $\mathcal{J}$ are then given by $\lambda = \lambda_{\text{re}} \pm i\lambda_{\text{im}}$. Let us denote $\ell_{\text{re}} = -\lambda_{\text{re}}\,\tau_\eta$, and $\ell_{\text{im}} = \lambda_{\text{im}}\,\tau_\eta/(2\pi)$. To understand the effects of these eigenvalues on network oscillations, we used Eqs. S23, S26 and S32 to compute $\mathbf{\Sigma}^*$ and $\mathbf{\Sigma}$ and the autocorrelation function of the network (Fig. S14). We found that while $\ell_{\text{re}}$ controls the degree of oscillatoriness of the system, $\ell_{\text{im}}$ has little to no effect on it (and instead controls the oscillation frequency as expected; compare Fig. S14a-b). Note, however, that $\ell_{\text{re}}$ does not control the oscillatoriness of the system quite as independently from the envelope of decay in the same way as $\alpha$ did it in our parametric fit (Eqs. S12 and S22).

It is also instructive to analyze how the total output variance is related to the real and imaginary parts of the Jacobian. In particular, the total variance (given by the trace of $\mathbf{\Sigma}$) decreases with $\ell_{\text{re}}$, which in turns implies reduced oscillatoriness (Fig. S14c). This predicts a positive correlation between oscillatoriness and variance: larger variance should be associated with small $\ell_{\text{re}}$ and thus with a larger degree of oscillatoriness. This is consistent with what we observed across the different PCs in the full network (Fig. 6c-d). Moreover, the total variance also decays with the square of the oscillation frequency $\omega = \lambda_{\text{im}}$ (Fig. S14d), corroborating our earlier explanation of why oscillation frequencies tend to increase with contrast (as output variance decreases).

---

[4] Note that our approximation to $\mathcal{J}$ is only expanding to 0th order in $\tau_\eta$, rather than 1st order as until now – otherwise we would have an additional $-\frac{1}{2}\,\mathbf{Q}_\circ\,\mathbf{\Sigma}_{\mathcal{P}}^{-1}$ term in Eq. S47. This explains why we get purely oscillatory dynamics rather than partial oscillatoriness as derived in Section S5.2.2 and seen in the real network. Nevertheless, in this section our focus is on understanding specifically the oscillatory components of the dynamics, and so we work with this approximation here.

### S5.3 Transients

In this section we first analyze how transients in neural responses affect sampling accuracy in general, and then derive and study the analytical form of the optimal transient.

#### S5.3.1 Transients and sampling accuracy

To evaluate if and how transient overshoots benefit sampling accuracy over finite time windows, we followed a procedure analogous to the one we used for analyzing oscillations (Section S5.2). Specifically, we devised and compared a set of one-dimensional Gaussian processes to phenomenologically model the behavior of a single cell around the time of stimulus presentation, either respecting the transient behavior of our network, or breaking it in various ways. All of these GPs were required to sample from the spontaneous distribution of a representative network neuron before stimulus onset (time $t < 0$, representing the posterior distribution of the GSM appropriate for a 0-contrast stimulus), but to eventually converge to the (marginal) posterior distribution for the chosen stimulus (for $t \gg 0$). Our surrogate GPs differed only in their transient behavior. The GP that most faithfully captured our optimized network was constructed by taking the temporal evolution of the mean and variance of an actual neuron in the full network around stimulus onset, including overshoots ("with overshoots" in Fig. 7a, and Fig. S15a, red). For this, we chose the neuron whose preferred orientation matched that of the presented stimulus. Moreover, the autocorrelogram of this GP was also set to match that neuron's autocorrelogram (inset of Fig. S15a, right). Two additional GPs were constructed with the same autocorrelogram as the first GP, but different mean and variance time courses. In one, the mean and variance converged exponentially with a time constant of $\tau_\mathrm{E}$ ("exponential" in Fig. 7a, and Fig. S15a, dashed black). In the other, the mean and variance immediately jumped at stimulus onset to their new stationary values ("instantaneous" in Fig. 7a, and Fig. S15a, dashed gray). While such an instantaneous process is not realisable by any continuous dynamical system, it provides a useful lower bound on sampling error.

Sampling error was quantified by $\overline{\mathrm{D}}_\mathrm{SKL}$ (Eq. S17), computed at each time $t$ between the momentary target distribution (which switched abruptly from the spontaneous to the evoked posterior at $t = 0$) and the momentary empirical distributions represented by the samples from each of our three processes in a 100 ms-long sliding time window (ending at $t$). Transient overshoots in the network were indeed found to benefit burn-in behavior: the GP which faithfully captured our network's transient behavior led to faster reduction in $\overline{\mathrm{D}}_\mathrm{SKL}$ following stimulus onset than the exponentially converging surrogate. Indeed, transient overshoots produced near-optimal convergence, i.e. almost as fast as the instantaneous system (Fig. 7b and Fig. S15b[5]).

#### S5.3.2 Optimal transient responses for continual estimation

Any system performing inference via sampling in real time faces the challenge of approximating a time-varying posterior using samples over a finite time window. In the previous section, we have shown that transient overshoots are useful in this setting. To explain this effect, here we analyzed the related (albeit simpler) task of generating (one dimensional[6]) time-varying neural activity $u(t)$ whose running average, $\mu_\mathcal{Q}(t)$[7], most closely tracked a time-varying target signal $\mu_\mathcal{P}(t)$ (e.g. the posterior mean in a continual inference scenario). We took the running average to be the convolution of $u(t)$ with a kernel $k(t)$:

$$\mu_\mathcal{Q}(t) = \big(u * k\big)(t)\,. \tag{S51}$$

More specifically, our goal was to find the $u(t)$ that minimized:

$$\tilde{\mathrm{D}} = \int \left[\big(\mu_\mathcal{Q}(t) - \mu_\mathcal{P}(t)\big)^2 + \epsilon_\mathrm{smooth}\left|u'(t)\right|^2\right] \mathrm{d}t \tag{S52}$$

[5] Fig. 7b only shows the term of $\overline{\mathrm{D}}_\mathrm{SKL}$ that depends on the sample mean, $\mu_\mathcal{Q}(T)$ (via $\mathcal{E}^2(T)$): $\overline{\mathrm{D}}^{\mu}_\mathrm{SKL} = \frac{1}{4}\mathbb{E}\left[\mathcal{E}^2(T)\,\left(\sigma_\mathcal{P}^{-2} + \sigma_\mathcal{Q}^{-2}(T)\right)\right]$. Fig. S15b shows the full $\overline{\mathrm{D}}_\mathrm{SKL}$ (Eq. S17). Expectations in both cases were taken across time, as well as across trials (see also Footnote 7 and Eq. S52).

[6] As before, the results generalize to multidimensional responses.

[7] Note that here we have slightly overloaded our previous notation, as $t$ in $\mu_\mathcal{Q}(t)$ refers to the *point in time* up until which we average responses, rather than the *length of time* over which we average, as $T$ did in the similar notation $\mu_\mathcal{Q}(T)$ used in Sections S5.1 and S5.2. The duration of the averaging window here is implicit in the definition of $k(t)$.

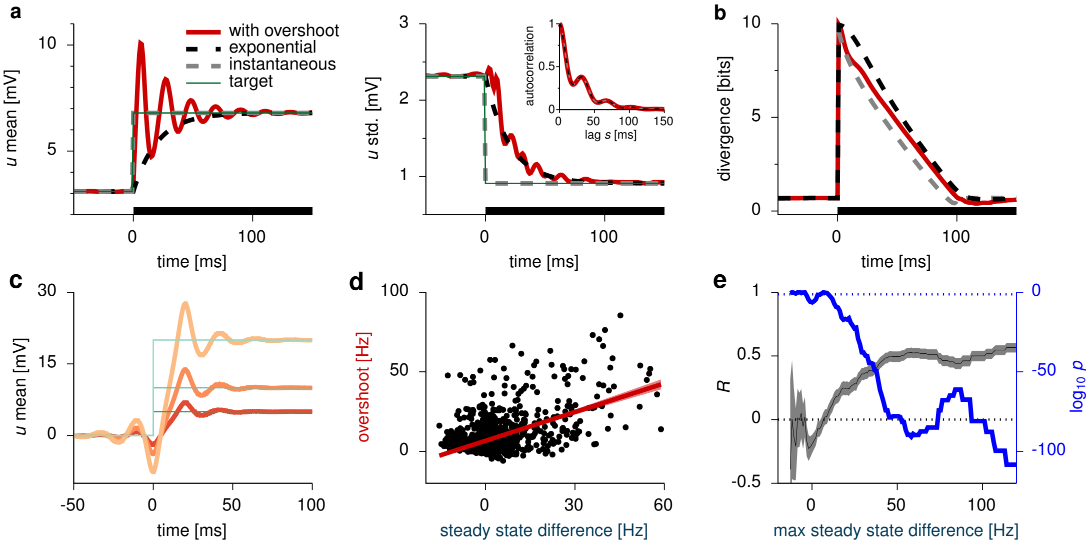


**Fig. S15 | Transients support continual inference (cont'd). a-b,** Analysis of transients in the response of a single neuron. (Left panel in **a** is reproduced from Fig. 7a.) **a,** Temporal evolution of the mean (left) membrane potential ($u_E$), and the membrane potential standard deviation (right) in three different neural responses (thick lines) with identical autocorrelations (matched to neural autocorrelations in the full network, inset, cf. Fig. 6) but different time-dependent means (left) and standard deviations (right). Thin green line shows the time-varying target mean (left) and standard deviation (right). **b,** Total divergence (Eq. S17 and footnote 5) between the target distribution at a given point in time and the distribution represented by the neural activity sampled in the preceding 100 ms, for each of the three responses (colors as in **a**). In comparison, note that Fig. 7b only shows the mean-dependent term of the divergence (Footnote 5). Black bars in **a**-**b** show stimulus period. **c,** Optimal response trajectories (red lines) for continual estimation of the mean of a target distribution (thin green lines). Different shades of red indicate optimal trajectories corresponding to three different target levels (Eq. S54). **d-e,** Relationship between overshoot magnitude and steady state difference: analysis of experimental recordings from awake macaque V1 (Ecker et al., 2010). **d,** Overshoot magnitude versus steady state difference: same as Fig. 7e, but restricting the analysis to steady state differences below 60 Hz (to exclude outliers). Red line shows linear regression (±95% confidence bands). The correlation between overshoot size and steady state difference is still significant ($R^2 = 0.27$, $p < 0.001$). **e,** Systematically changing the maximal steady state difference (x-axis) used for restricting the analysis of the correlation between overshoot size and steady state difference (panel **d** here and Fig. 7e) reveals that the correlation is robust (black line ±95% confidence intervals) and remains highly significant (blue line, showing corresponding p-values; note logarithmic scale) for all but the smallest threshold (and thus smallest sample size). Horizontal dotted lines show $R = 0$ correlation (black) and $p = 0.05$ significance level (blue) for reference.

where the first term penalized the average squared distance between $\mu_{\mathcal{Q}}(t)$ and the time-varying target $\mu_{\mathcal{P}}(t)$ (and as such, it was closely related to $\overline{\mathrm{D}}^{\mu}_{\mathrm{SKL}}$, Footnote 5), and the second term encouraged temporal smoothness by penalizing high (squared) temporal derivatives, $u'(t)$, such that the optimal $u(t)$ could have been produced by a continuous dynamical system. By taking the Fourier transform of Eq. S52, and making use of Plancherel's theorem, we obtain:

$$\tilde{\mathrm{D}} = \int \left( \left| \hat{u}(f)\ \hat{k}(f) - \hat{\mu}_{\mathcal{P}}(f) \right|^2 + \epsilon_{\mathrm{smooth}}\ (2\pi f)^2\ \left| \hat{u}(f) \right|^2 \right) \mathrm{d}f \tag{S53}$$

where $\hat{\cdot}$ denotes the Fourier transform. Setting $\frac{\mathrm{d}\tilde{\mathcal{D}}}{\mathrm{d}u}$ to zero, we obtain the optimal $\hat{u}$:

$$\hat{u}(f) = \frac{\hat{\mu}_{\mathcal{P}}(f)\ \hat{k}^*(f)}{\left| \hat{k}(f) \right|^2 + 4\,\epsilon_{\mathrm{smooth}}\,\pi^2 f^2}, \tag{S54}$$

where $\cdot^*$ denotes the complex conjugate. The optimal $u(t)$ can then be obtained from Eq. S54 via the inverse Fourier transform.

As an example, we considered a prospective box-car filter ($k(t) = 1/T$ for $t \in [0, T]$ and 0 otherwise), and the Heaviside step function as the target signal. We then computed the trajectories $u(t)$ (Fig. S15c, solid lines in shades of red) that minimized the cost function given by Eq. S52 for $\epsilon_{\text{smooth}} = 1$, $T = 20$ ms, and three different values for the target $\mu_{\mathcal{P}}$ (5, 10, and 20 mV, dashed green lines), emulating different contrast levels. We observed a transient overshoot in the optimal response that very closely resembled those observed in the optimized network: its magnitude scaled with the value of the target mean, and it was followed by damped oscillations. Although the intuition for this result is most straightforward for the simple box-car kernel used here, qualitatively similar results (with less ringing following the overshoot) were obtained for example for an exponentially decaying, "leaky" kernel (not shown).

### S6 Online repositories

Code and parameters for the numerical experiments: bitbucket.org/RSE_1987/ssn_inference_numerical_experiments
Code for the optimization procedure: bitbucket.org/RSE_1987/ssn_inference_optimizer

| Parameter | Description | Value |
|---|---|---|
| | Generative model | |
| $N_\mathrm{x}$ | Number of observed variables (pixels) | 256 |
| $N_\mathrm{y}$ | Number of latent variables | 50 |
| $\mathbf{A}$ | Matrix of Gabor oriented filters | (see Fig. S1 and repository) |
| $\mathbf{C}$ | Prior covariance matrix | (see Fig. S1 and repository |
| $\sigma_\mathrm{x}$ | Standard deviation of the pixel noise | 10.0 |
| $K$ | Shape parameter of the gamma prior on contrast | 2.0 |
| $\vartheta$ | Scale parameter of the gamma prior on contrast | 2.0 |
| $\alpha_\mathrm{nl}$ | Scaling of the non-linear transformation | 2.4 |
| $\beta_\mathrm{nl}$ | Baseline of the non-linear transformation | 1.9 |
| $\gamma_\mathrm{nl}$ | Power of the non-linear transformation | 0.6 |
| | Network | |
| $N$ | Total number of neurons | 100 |
| $N_\mathrm{E}$ | Number of E cells | 50 |
| $N_\mathrm{I}$ | Number of I cells | 50 |
| $\tau_\mathrm{E}$ | Membrane time constant of E cells | 20ms |
| $\tau_\mathrm{I}$ | Membrane time constant of I cells | 10ms |
| $\tau_\eta$ | Process noise time constant | 20ms |
| $\mathbf{W}$ | Weight matrix | (optimized, see repository) |
| $n$ | Exponent in the single cell transfer function | 2 |
| $k$ | Scaling factor in the single cell transfer function | 0.3 |
| $\mathbf{\Sigma}_\eta$ | Process noise covariance | (optimized, see repository) |
| $\mu_0$ | Mean of the initial distribution | $\mathbf{0}$ |
| $\mathbf{\Sigma}_0$ | Covariance of the initial distribution | $4\,\mathbf{I}$ |
| $dt$ | simulation time-step | 0.2 ms |
| | Input | |
| $\mathbf{W}^\mathrm{ff}$ | Feed-forward weights | $[\mathbf{A}\,\mathbf{A}]^\mathrm{T}/15.0$ |
| $\alpha_\mathrm{h}$ | Input scaling | 1.96 (optimized) |
| $\beta_\mathrm{h}$ | Input baseline | 0.10 (optimized) |
| $\gamma_\mathrm{h}$ | Input power | 2.03 (optimized) |
| | Fano factor calculation | |
| $T$ | Observation window | 100 ms |
| $K_\mathrm{ISI}$ | Shape parameter of interspike interval gamma distribution | 1.15 |
| | Input conductance calculation | |
| $C_\mathrm{m}$ | Membrane capacitance | 20.0 pF |
| $V_\mathrm{rest}$ | Resting membrane potential (reference level for **u**) | -65 mV |
| $V^\mathrm{E}$ | E reversal potential | 0 mV |
| $V^\mathrm{I}$ | I reversal potential | -80 mV |
| | Optimization | |
| $\epsilon_\mathrm{mean}$ | Cost weight for the error of the mean | $4.0 \cdot 10^{-5}$ |
| $\epsilon_\mathrm{var}$ | Cost weight for the error of the variance | $8.0 \cdot 10^{-5}$ |
| $\epsilon_\mathrm{cov}$ | Cost weight for the error of the covariance | $8.0 \cdot 10^{-7}$ |
| $\epsilon_\mathrm{slow}$ | Cost weight for the slowness penalty | $4.0 \cdot 10^{-7}$ (ADF only) |
| $T_\mathrm{max}$ | Stimulus presentation time/ integration end time | 500 ms |
| $T_\mathrm{min}$ | Integration start time | 0 to 450 ms |
| $\tau_\mathrm{max}$ | Autocorrelation integration time | 100 ms |
| $N_\mathrm{trials}$ | Number of trials for stochastic optimization | 50 |
| $dt'$ | simulation time-step during optimization | 0.2 ms |

**Table S1 |** List of parameter values.
